# Some Statistical Consideration in Transcriptome-Wide Association Studies

**DOI:** 10.1101/812677

**Authors:** Haoran Xue, Wei Pan, for the Alzheimer’s Disease Neuroimaging Initiative

**Author notes:** Data used in preparation of this article were obtained from the Alzheimer’s Disease Neuroimaging Initiative (ADNI) database (adni.loni.usc.edu). As such, the investigators within the ADNI contributed to the design and implementation of ADNI and/or provided data but did not participate in analysis or writing of this report. A complete listing of ADNI investigators can be found at: http://adni.loni.usc.edu/wp-content/uploads/how_to_apply/ADNI_Acknowledgement_List_Sep23.pdf.

## Abstract

Transcriptome-wide association study (TWAS) has become popular in integrating a reference eQTL dataset with an independent main GWAS dataset to identify (putatively) causal genes, shedding mechanistic insights to biological pathways from genetic variants to a GWAS trait mediated by gene expression. Statistically TWAS is a (two-sample) 2-stage least squares (2SLS) method in the framework of instrumental variables analysis for causal inference: in Stage 1 it uses the reference eQTL data to impute a gene’s expression for the main GWAS data, then in Stage 2 it tests for association between the imputed gene expression and the GWAS trait; if an association is detected in Stage 2, a (putatively) causal relationship between the gene and the GWAS trait is claimed. If a non-linear model or a generalized linear model (GLM) is fitted in Stage 2 (e.g. for a binary GWAS trait), it is known that using only imputed gene expression, as in standard TWAS, in general does not lead to a consistent (i.e. asymptotically unbiased) estimate for the causal effect; accordingly, a variation of 2SLS, called two-stage residual inclusion (2SRI), has been proposed to yield better estimates (e.g. being consistent under suitable conditions). Our main goal is to investigate whether it is necessary or even better to apply 2SRI, instead of the standard 2SLS. In addition, due to the use of imputed gene expression (i.e. with measurement errors), it is known that in general some correction to the standard error estimate of the causal effect estimate has to be applied, while in the standard TWAS no correction is applied. Is this an issue? We also compare one-sample 2SLS with two-sample 2SLS (i.e. the standard TWAS). We used the ADNI data and simulated data mimicking the ADNI data to address the above questions. At the end, we conclude that, in practice with the large sample sizes and small effect sizes of genetic variants, the standard TWAS performs well and is recommended.

## 1 Introduction

Genome-wide association studies (GWAS) have been successful in identifying thousands of trait-associated genetic variants, mostly single nucleotide polymorphisms (SNPs). However, since most of the identified trait-associated SNPs are in the non-coding region of the genome, there is a lack of mechanistic understanding of how these SNPs influence the traits. It is hypothesized that many genetic variants influence complex traits through transcriptional regulation (He et al., 2013), which can be used to identify *causal* genes. For this purpose, PrediXcan (Gamazon et al., 2015) and transcription-wide association study (TWAS) (Gusev et al., 2016), simply called **TWAS** from now on, were recently proposed to uncover *putatively* causal genes by integrating a main GWAS dataset with a reference gene expression or expression quantitative trait (eQTL) dataset. TWAS has since become popular and successful in applications to common diseases like T2D and cancer, and to complex traits like BMI, lipids and height, convincingly showing the power of integrating GWAS and eQTL data to gain biological insights. Statistically, TWAS applies the (two-sample) two-stage least squares (2SLS) method for causal inference Xu, Wu, Wei & Pan, 2017a), closely related to Mendelian randomization (MR) (Zhao, Wang, Hemani, Bowden, & Small, 2019). Since TWAS is a gene-based method by testing genes one by one, for the purpose of presentation we can consider only one gene. In Stage 1, one builds a prediction model for the genetic component of the gene’s expression level, called “genetically regulated expression (GReX)”, by using only cis-acting genotypes around the gene based on a reference eQTL dataset. In stage 2, for a given separate main GWAS dataset, based on the genotype of each subject, we can “impute” his/her gene expression (i.e. GReX) using the predictive model built in Stage 1. Then we test the association between the imputed gene expression and the GWAS trait. If there is an association, then, under suitable modeling assumptions (Xu, Wu, Wei, & Pan, 2017; Mancuso et al., 2019; Hu et al., 2019; Wainberg et al., 2019), it is claimed that the gene is (putatively) causal to the trait: some causal SNPs affect the trait through the mediating effects of the gene’s expression.

As to be reviewed next, if the GWAS trait is quantitative, a linear model is usually used in Stage 2 to test the association between the imputed gene expression level and the trait, which is exactly 2SLS. However, in practice, a non-linear model (with an additive error term) may be used; or, more often, since in many GWAS the trait is not quantitative, e.g. being binary as an indicator of a disease status, a generalized linear model (GLM), e.g. a logistic regression model, is instead fitted. For such a non-linear model in Stage 2, it is known that the usual 2SLS, specifically called Two-Stage Predictor Substitution (2SPS), may not be consistent; instead, for a non-linear model with an additive error term, Two-Stage Residual Inclusion (2SRI) is consistent and should be applied (Terza, Basu, & Rathouz, 2008; MacKenzie, Tosteson, Morden, Stukel & O’Malley, 2014), while for a logistic regression model, an equivalent method to 2SRI has been proposed (Palmer et al., 2008, 2011). Hence, an important question is whether it is indeed suitable to apply 2SPS for binary traits in Stage 2 as in the current practice of TWAS. From now on, we use 2SLS as a *generic* term covering both 2SPS and 2SRI. In addition, TWAS is based on a two-sample 2SLS (2S-2SLS), in which two separate and independent datasets are used in Stages 1 and 2 respectively; there is a corresponding one-sample 2SLS (1S-2SLS) with the two datasets in the two stages collected from the same set of subjects (Angrist & Krueger, 1991). This is convenient with two separate eQTL and GWAS datasets. However, in practice, there are situations when both eQTL and GWAS are collected on the same set (or largely overlapping sets) of subjects. In these situations, should be split the dataset into two non-overlapping subsets before applying 2S-2SLS, or apply 1S-2SLS to the whole dataset? To answer this question, we need to investigate how 1S-2SLS performs in the context of TWAS. We will use both a real dataset and simulated data to address these questions.

## 2 Methods

### 2.1 The True Models in Instrumental Variables Analysis

The true (causal) model is illustrated with a directed acyclic graph (DAG) in Figure 1, where *Y, Z* and *U* are the gene, GWAS trait and unobserved confounders respectively, and a directed edge solid between *U* and *Z, U* and *Y, SNP*_*j*_ and *Y* (for *j* = 1, …, *m*) represents a direct causal effect. Our goal is to test whether *Y* has a direct effect on *Z*, and possibly estimate its effect size.

**Figure 1:**
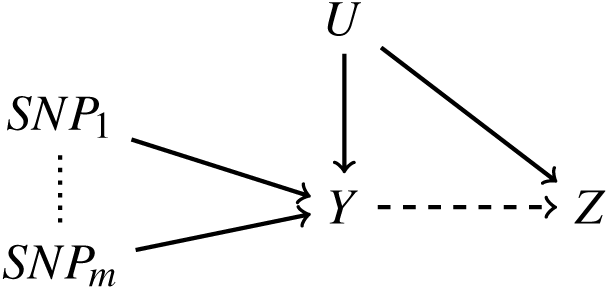
True model.

For individual *i, i* = 1, …, *n*, with gene expression level *Y*_*i*_, *SNP*s *SNP*_1,*i*_, …, *SNP*_*m,i*_, and binary trait *Z*_*i*_. Let *p*_*i*_ be the probability of *Z*_*i*_ = 1. Our the true model for Stage 1 is:

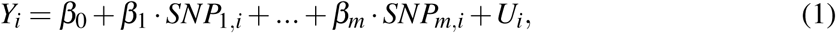

where, as usual, throughout, we use additive coding for each SNP: *SNP*_*j,i*_ = 0, 1 or 2 for *j* = 1, …, *m* and *i* = 1, …, *n*. The true model for Stage 2 is:

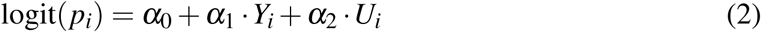

with *p*_*i*_ = Pr(*Y*_*i*_ = 1). A main challenge is that we do not observe the confounder *U*_*i*_ that is correlated with both *X*_*i*_ and *Y*_*i*_.

The above true model is the typical one adopted in the instrumental variables (IVs) analysis for causal inference, in which the SNPs are taken as IVs. The focus is statistical inference on the causal effect *α*_1_. A main benefit is consistent estimation and inference for *α*_1_ at the expense of three assumptions with IVs, as in MR: 1) the SNPs/IVs are associated with *Y*; 2) the SNPs/IVs are not associated with *U*; 3) Conditional on *Y*, the SNPs/IVs are not associated with *Z*. If any of the above three modeling assumptions is violated, biased inference for *α*_1_ results, as discussed in the context of TWAS (Mancuso et al., 2019; Wainberg et al., 2019). Since it is not the focus of this paper while it is quite challenging to deal with, we will assume that the three assumptions hold in the following as in standard scenarios.

### 2.2 One-sample versus Two-sample Approaches

All methods we are going to introduce consist of two stages. If we have both eQTL and GWAS data from the same sample of subjects, we have two choice of how to use the data. First, we can use the whole sample for both Stages 1 and 2, which is denoted as **one-sample** approach. Alternatively, we can randomly split the whole sample into half-half (or in whatever desired ratio), using the first half for Stage 1, and the other half for Stage 2, which is denoted as **two-sample** approach.

Denote *I*_1_, *I*_2_ ⊆ {1,*…, n*} be the index sets for samples used in Stages 1 and 2 respectively. For the one-sample strategy, *I*_1_ = *I*_2_ = {1,*…, n*}; for the two-sample approach, we have *I*_1_ *∩ I*_2_ = ø.

### 2.3 TWAS: Four Methods to Implement It

We implement TWAS in four different ways in Stage 2, depending on whether a LM or GLM is fitted to the binary GWAS trait and how to use imputed gene expression, leading to four different methods, denoted as GLM 2SPS, GLM 2SRI, LM 2SPS and LM 2SRI. The (standard) TWAS corresponds to LM 2SPS. All of these four methods are variants of 2SLS, consisting of two stages; they share fitting the same LM in Stage 1, but differ in fitting different models in Stage 2.

In Stage 1, we fit a LM by regressing gene expression *Y* on the SNPs, *SNP*_1_, …, *SNP*_*m*_:

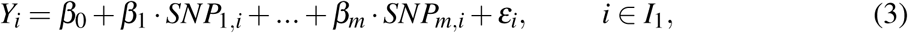

where *ε*_*i*_ is assumed to be a random noise with mean 0 and independent of the SNPs. We obtain the OLS estimates 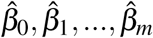, and predict (or impute) gene expression *Ŷ*_*i*_’s for *i* ∈ *I*_2_. We also estimate *Û*_*i*_ = *Y* − *Ŷ*_*i*_ for confounders *U*_*i*_, *i* ∈ *I*_2_.

In Stage 2, for GLM-2SPS, we fit a logistic regression model using *Ŷ*_*i*_:

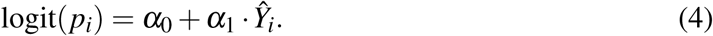

In contrast, for GLM-2SRI, we fit a logistic model using both *Y*_*i*_ and *Û*_*i*_:

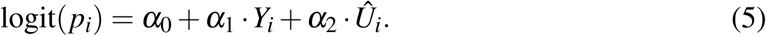

Note the use of *Y*_*i*_, not *Ŷ*_*i*_, in the above model, which will not be possible for usual two-sample scenarios.

As alternatives, we can fit two LMs in Stage 2. The first corresponds to LM-2SPS:

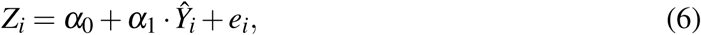

where *e*_*i*_ is assumed to be a random noise with mean 0 and independent of the SNPs. The second is for LM-2SRI:

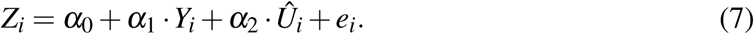

For each method, we draw inference on the causal effect *α*_1_ after obtaining its estimate 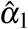 and standard error se 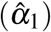. For GLM-2SRI and LM-2SPS, there are existing methods for correct se 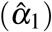 as to be discussed next, while for others we will use the output from standard software fitting their corresponding models in Stage 2.

It is noted that, although we have one-sample ADNI data with the availability of the genotype, gene expression and GWAS trait (AD) data for each subject, to mimic a two-sample approach, we split the whole sample of the ADNI data into two non-overlapping samples. An advantage of the two-sample approach is the relaxed assumption of the availability of the gene expression and the GWAS trait in the two samples respectively, to which the two-sample LM- or GLM-2SPS method (as the standard TWAS) can be applied; however, the two-sample LM- or GLM-2SRI does require the availability of the gene expression data in both samples (but only the availability of the GWAS trait in the second sample).

Given the true data generating models (1) and (2), and the working models (3) and (4), although the estimation of the causal effect *α*_1_ in (2) is not consistent in general, the test for whether *α*_1_ = 0 is valid and consistent (xDai & Zhang, 2015). Furthermore, we note that GLM-2SRI with model (5) is equivalent to the so called “Adjusted IV Estimator” (Palmer, Thompson, Tobin, Sheehan & Burton, 2008), also called “Control Function Estimator” (Palmer et al., 2011) for binary traits, in which *Y*_*i*_ is replaced by *Ŷ*_*i*_ in (5).

### 2.4 Corrections for se 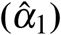

For one-sample GLM 2SRI, we correct the *se* 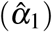 with following equation (Terza, 2018):

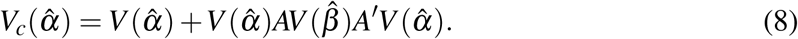

Here *V* 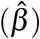 is the original covariance matrix of 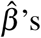 in the first stage, and *V* 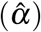 is the original covariance matrix of 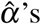 in the second stage; throughout, the “original” one means the usual covariance matrix directly output from fitting a standard model in Stage 1 or 2. *V*_*c*_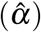 is the corrected covariance matrix for 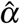. We define:

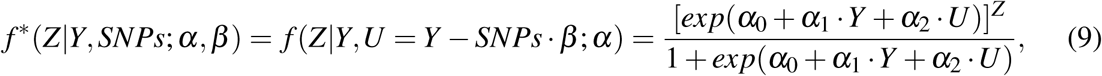

and let 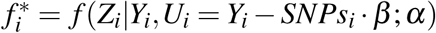 and 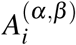:

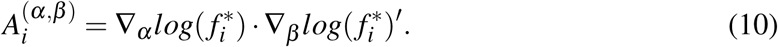

Then *A* is calculated as

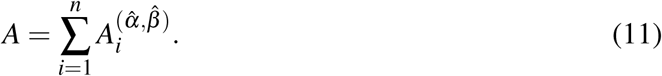

We can notice that the corrected *se*_*c*_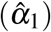 is larger than the original *se* 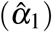. Though the correction was originally designed for one-sample GLM-2SRI, we will also apply it to two-sample GLM-2SRI.

For one-sample LM-2SPS, the usual standard error estimate requires the homoskedasticity assumption on the error term in a LM; to allow heterskedastic errors, a robust or sandwich-type standard error estimator has been proposed (Baiocchi, Cheng, & Small, 2014; Imbens & Angrist, 1994; Angrist & Pischke, 2009). We can use the `robust.se()` function in R package **ivpack** to obtain the robust standard error of 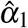 in LM 2SPS (Baiocchi, Cheng, & Small, 2014).

For two-sample LM-2SPS, to account for the statistical uncertainty or estimation error of estimating/imputing *Y*_*i*_ as *Ŷ*_*i*_ in Stage 1, we can correct the usual *se* (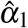) by inflating it with a factor no smaller than 1 (Inoue, Atsushi & Solon, 2010):

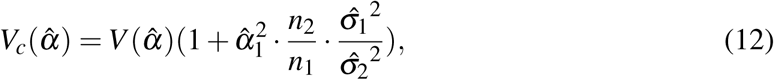

where *V* 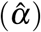 is the original covariance matrix of 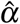, *n*_1_ and *n*_2_ are the sample sizes in Stages 1 and 2 respectively, and 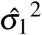 and 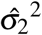 are the estimated variances for the error terms in the two LMs in the two stages respectively.

We thus obtain the corrected standard errors (SEs) for both one-sample and two-sample LM-2SPS, and for both one-sample and two-sample GLM-2SRI. For other methods, we are not aware of any existing methods to correct their SEs and thus just use the standard/uncorrected SEs from the output from fitting their corresponding models in Stage 2.

### 2.5 ADNI Data

Data used in the preparation of this article were obtained from the Alzheimer’s Disease Neuroimaging Initiative (ADNI) database (`adni.loni.usc.edu`). The ADNI was launched in 2003 by the National Institute on Aging (NIA), the National Institute of Biomedical Imaging and Bioengineering (NIBIB), the Food and Drug Administration (FDA), private pharmaceutical companies and non-profit organizations, as a 60 million, 5-year public private partnership. The primary goal of ADNI has been to test whether serial magnetic resonance imaging (MRI), positron emission tomography (PET), other biological markers, and clinical and neuropsychological assessment can be combined to measure the progression of mild cognitive impairment (MCI) and early Alzheimer’s disease (AD). Determination of sensitive and specific markers of very early AD progression is intended to aid researchers and clinicians to develop new treatments and monitor their effectiveness, as well as lessen the time and cost of clinical trials. The Principal Investigator of this initiative is Michael W. Weiner, MD, VA Medical Center and University of California — San Francisco. ADNI is the result of efforts of many co-investigators from a broad range of academic institutions and private corporations, and subjects have been recruited from over 50 sites across the U.S. and Canada. The initial goal of ADNI was to recruit 800 subjects but ADNI has been followed by ADNI-GO and ADNI-2. To date these three protocols have recruited over 1500 adults, ages 55 to 90, to participate in the research, consisting of cognitively normal older individuals, people with early or late MCI, and people with early AD. The follow up duration of each group is specified in the protocols for ADNI-1, ADNI-2 and ADNI-GO. Subjects originally recruited for ADNI-1 and ADNI-GO had the option to be followed in ADNI-2. For up-to-date information, see www.adni-info.org.

For real data analysis and for generating simulated data, we used the data from Alzheimer’s Disease Neuroimaging Initiative (ADNI) (Shen et al., 2014), including its gene expression, whole genome sequencing (WGS) and trait data. After cleaning and merging, we had a sample size 712. To mimic a case-control study, we treated 247 Cognitively Normal (CN) individual as controls, and the remaining 465 individuals with Alzheimer’s Disease (AD) or Mild Cognitive Impairment (MCI) as cases. The expression levels of 17256 genes on the autosomes were used. For each gene, we defined its cis-region by expanding 100kb upstream and downstream its coding region (i.e. from its TSS and TES) respectively. We excluded *SNP*s with MAF ≤ 0.05 or with missing values. We also pruned the SNPs to ensure that any of their pairwise Pearson correlations in absolute values was no more than 0.9. Finally, if there were still more than 30 *SNP*s in the cis-region of the gene, we chose and only kept the top 30 SNPs (as IVs) with the largest absolute values of the correlations with the gene’s expression level; if there were less than or equal to 30 *SNP*s, we kept all of them as IVs. In this way, we had *m* ≤ 30 *SNP*s as IVs so that the ordinary least squares (OLS) estimation could be applied in Stage 1.

For some genes, if their expression levels are not associated with their cis-SNPs, then using their cis-SNPs to predict their expression levels in Stage 1 would violate the first IV assumption, leading to the use of invalid IVs. As alternatives, for each gene we tested the association in equation (1) with the null hypothesis *H*_0_: *β*_1_ = *β*_2_ = … = *β*_*m*_ = 0. Under *H*_0_ (and the normality assumption for *Y*), the coefficient of determination *R*^2^ follows an *F*-distribution with degrees of freedom (*m, n m* 1). Thus, we performed the *F*-test on each gene in Stage 1, and only retained the genes with p-values less than some cut-off (0.05 or 0.1) before applying the methods.

### 2.6 Simulation Set-ups

To further study the methods, we conducted simulation studies by using the ADNI data to mimic realistic scenarios. We randomly selected gene **PSPH** on chromosome 7 to study. We first fitted a LM in the first stage with 30 *SNP*s we selected:

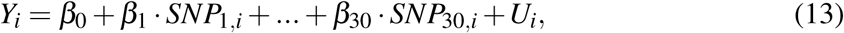

for *i* = 1, …, 712, to obtain 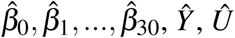 and *var*(*Û*). In the second stage, we first fitted a model like GLM 2SRI:

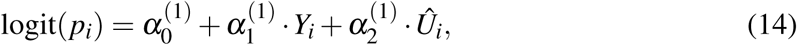

for *i* = 1, …, 712, to obtain 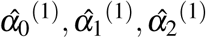 for a non-null model (with a causal effect 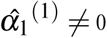.For a null model, we fitted a logistic regression model with only (*Û*):

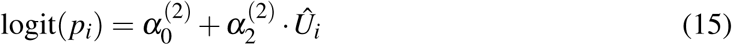

to obtain 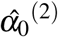 and 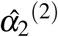. In the following simulation set-ups, we used these estimated coefficients to generate simulated data.

In each simulation set-up, we generated simulated data in the following steps.

For *i* = 1, …, *R* · 712:

1. Generate *U*_*i*_’s as i.i.d. Normal(0, *var* (*Û*));
2. Generate 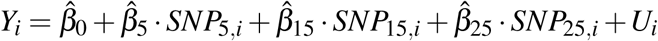;
3. Generate logit(*p*_*i*_). If *S*_*Y*_ = 0, then logit 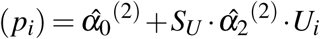; otherwise, logit 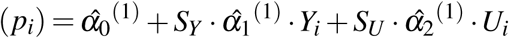;
4. Generate *Z*_*i*_ ∼ Bernoulli(*p*_*i*_).

Here *R* controlled the sample size, *S*_*Y*_ the effect size of the gene expression *Y*, and *S*_*U*_ the effect size of the confounder *U*. The values of the *SNP*s were drawn from the ADNI data: for *i* = *k* + *l* · 712, *k* = 1, …712, and *l* = 1, …, *R* − 1, we defined *SNP*_*j,i*_ = *SNP*_*j,k*_, meaning that we replicated the *SNP* data *R* times to possibly increase the sample size. We tried various combinations of (*R, S*_*Y*_, *S*_*U*_):

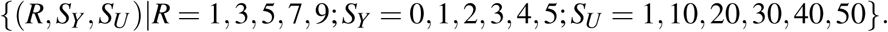

Note that we chose 3 *SNP*s as the true IVs: *SNP*_5_, *SNP*_15_, and *SNP*_25_. though we would use all 30 SNPs To mimic the real situation with valid IVs unknown, we would select 10 top SNPs as IVs from the 30 candidate SNPs as for the real data analysis. Then we fitted the LM in the first stage, and applied the four methods from formulas (4), (5), (6), and (7) in the second stage. Because we knew both *Y* and *U* with simulated data, we also considered the ideal (but not practical) *Oracle* method by fitting the true model (2) in the second stage.

For each simulated dataset, we applied the methods with both one-sample and two-sample approaches; for the former, we split a simulated dataset into half/half for the two stages respectively. For each simulation set-up, we generated 1000 independent simulated datasets.

### 2.7 Data Availability

The ADNI data are available to the approved user on the project web site http://adni.loni.usc.edu/. Some sample R code and data are available at https://github.com/xue-hr/twas_methods.

## 3 Results

### 3.1 ADNI Data Analysis

We applied the methods to 17256 genes for 712 individuals in ADNI data. For each of the four models, we tried both one-sample and two-sample approaches, obtaining the p-values for *α*_1_ for the 17256 genes, and draw Q-Q plots for these 17256 p-values in Figure 2.

**Figure 2:**
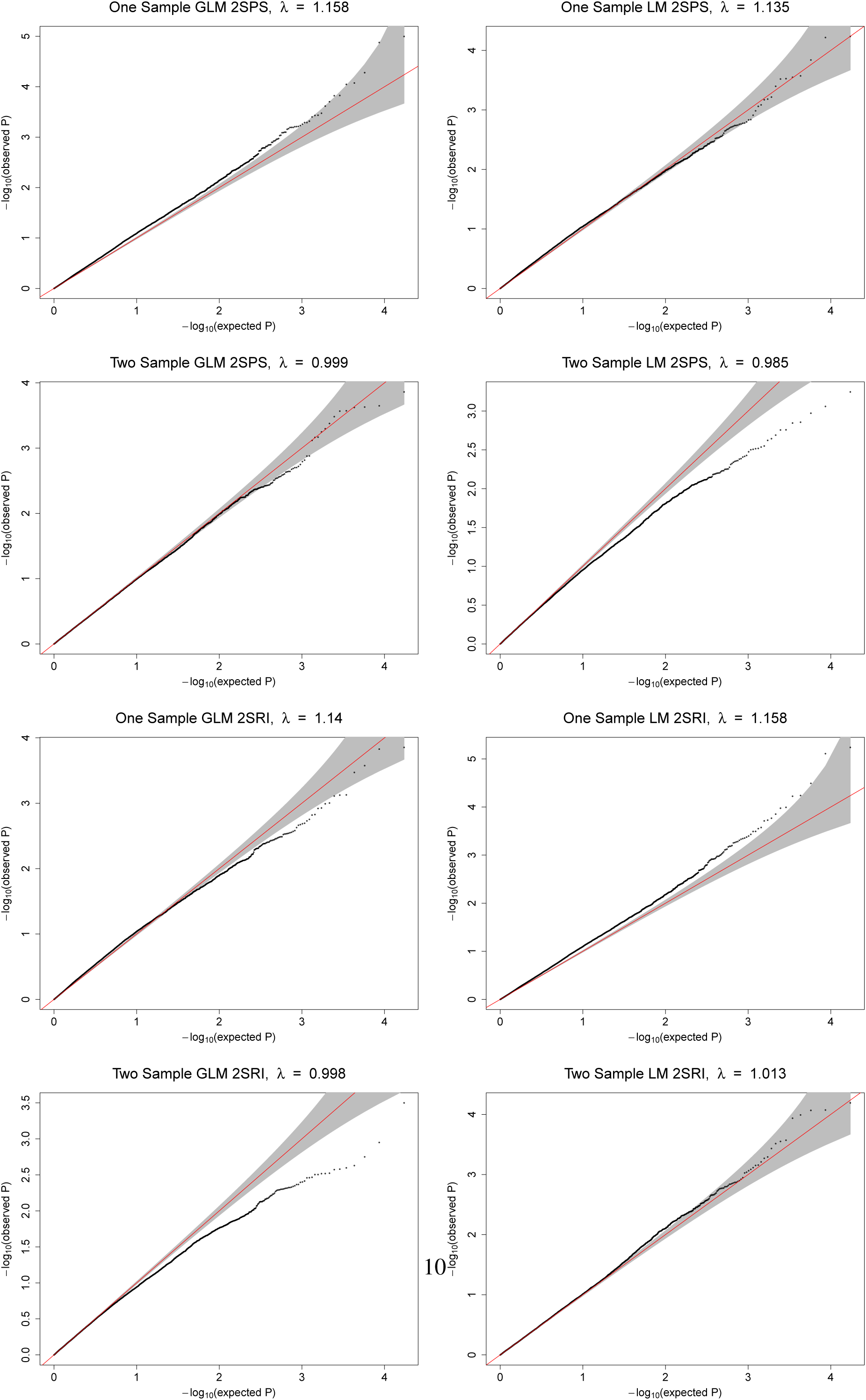
The ADNI data analysis: Q-Q plots of the obtained p-values of 17256 genes from each method versus the expected p-values under the null hypothesis of no association.

First, it is clear that all four methods with the one-sample approach led to inflated type I errors with inflated genomic control factors 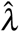 (Devlin & Roeder, 1999), likely due to not accounting for SNP selection in Stage 1 with the same sample as that in Stage 2. Among the four methods, 1S-GLM-2SPS and 1S-LM-2SRI was most liberal with much inflated type I errors, while 1S-GLM-2SRI was conservative at the left tail of the p-value distribution (i.e. perhaps over-estimating the more significant/smaller p-values); in contrast, 1S-LM-2SPS performed almost ok. Second, while all four methods with the two-sample strategy did not yield inflated 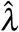, 2S-LM-2SPS and 2S-GLM-2SRI were too conservative, especially at the left tail of the p-value distribution; in contrast, 2S-GLM-2SPS performed almost ideally, followed by 2S-LM-2SRI.

We also conducted an F-test for a possible association between each gene’s expression levels and its cis-SNPs in Stage 1; we only retained the 9102 (or 10564) genes with a p-value < 0.05 (or 0.1) before applying the methods. The resulting Q-Q plots (Supplementary Materials) show the same patterns as discussed above; in particular, the inflation of the Type I error rates by the one-sample approaches was even more evident.

### 3.2 Simulation Results: Two-sample Approaches

#### 3.2.1 Type I errors

For the null case with no causal effects (i.e. *α*_1_ = 0 or *S*_*Y*_ = 0), the empirical Type I error rates at the nominal significance level of 0.05 for the four methods and the Oracle are shown in Figure 3 based on 1000 simulations; for comparison, the nominal significance level is marked with a gray horizontal line (in each corresponding figure). If the confounding is not severe with *S*_*U*_ = 1, all the methods performed satisfactorily. However, with more severe confounding with *S*_*U*_ = 50, first, LM-2SRI consistently had inflated Type I error rates, while GLM-2SRI also had an inflated type I error for the small sample size (with *R* = 1), which however disappeared with increasing sample sizes. In contrast, GLM-2SPS and LM-2SPS always controlled their Type I error rates satisfactorily.

**Figure 3:**
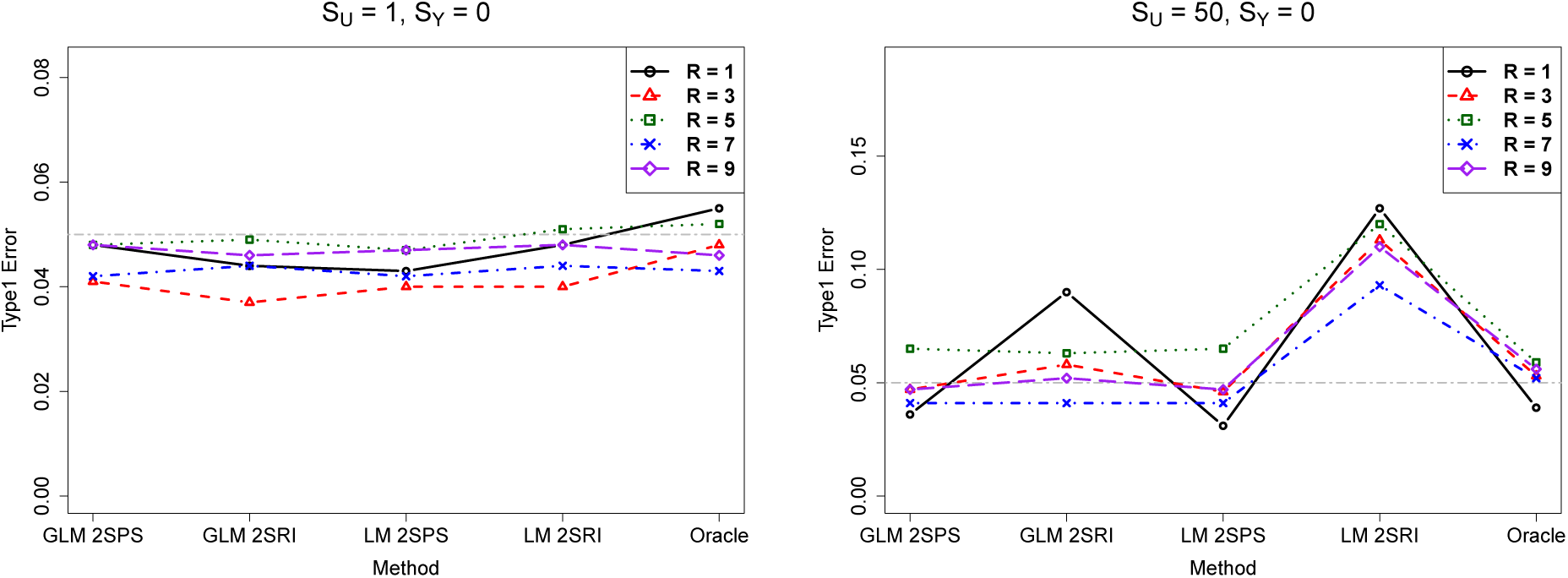
Simulations with the two-sample approaches: empirical Type I error rates of various methods.

#### 3.2.2 Power

In the presence of causal effects, as shown in Figure 4, with small confounding (with *S*_*U*_ = 1), all the methods performed similarly. However, with severe confounding (with *S*_*U*_ = 50), GLM-2SRI was low-powered, while GLM-2SPS and LM-2SPS performed similarly. Note that the high power of LM-2SRI was likely due to its inflated Type I error rates.

**Figure 4:**
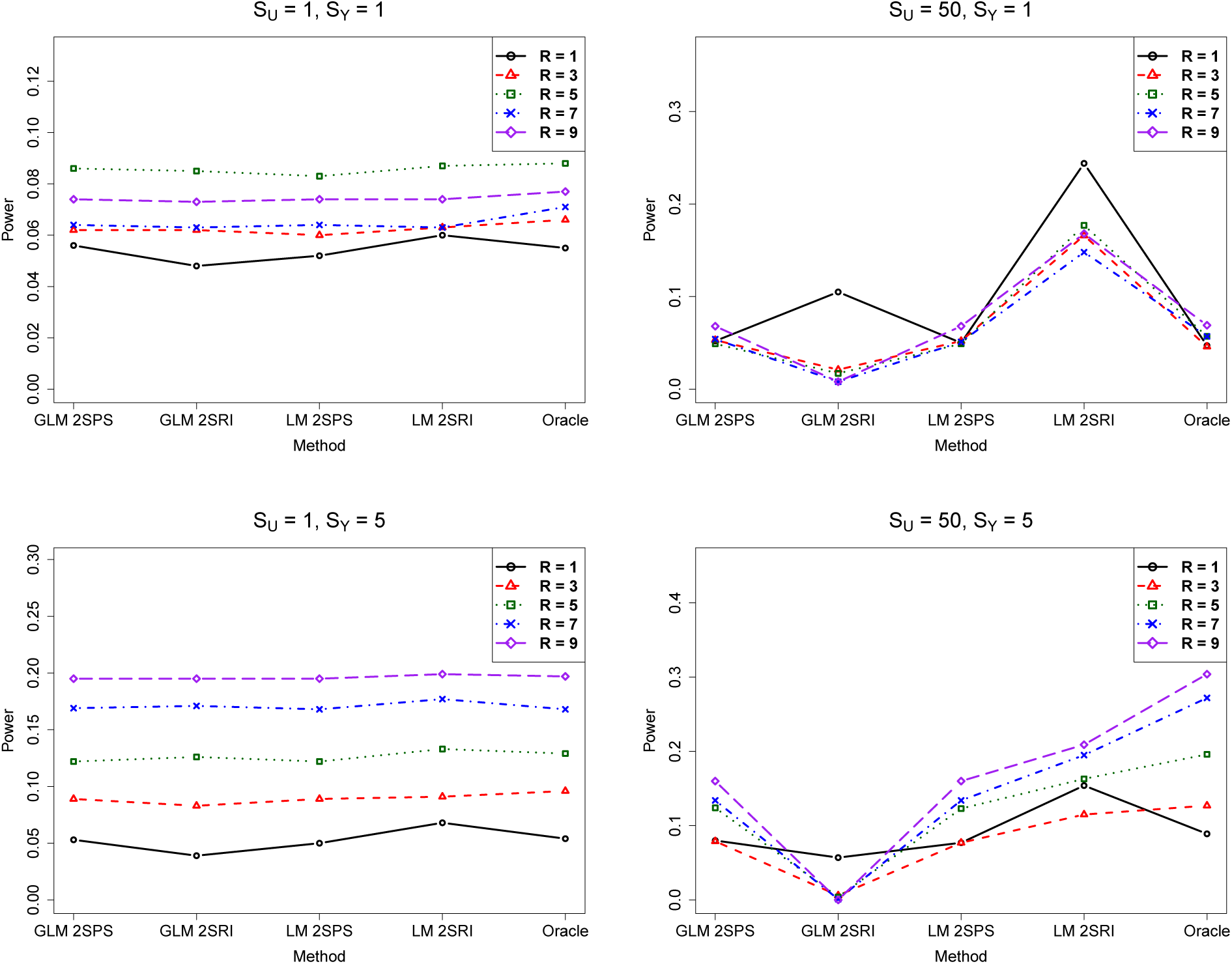
Simulations with the two-sample approaches: empirical power of various methods.

#### 3.2.3 Biases

We could compare the true causal effect size *α*_1_ with its estimate 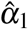 from each method to assess the extent of the bias, if any, by each method. Since the true model was a GLM in Stage 2, we only compared the GLM-based methods. For a specific simulation setting, based on 1000 simulations, we estimated the bias as 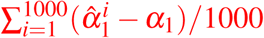, where 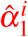 was the estimate of the true *α*_1_ in the *i*^*th*^ simulation.As shown in Figure 5, first, in the null case (i.e. *S*_*Y*_ = 0), with small confounding (i.e. small *S*_*U*_), the three methods performed similarly. However, with a large *S*_*U*_, at a small sample size GLM-2SPS and GLM-Oracle, but not GLM-2SRI, performed well; as the sample size increases, GLM-2SRI became less biased. Second, in the non-null case (i.e. *S*_*Y*_ ≠ 0), GLM-2SRI was less biased than GLM-2SPS in most cases, but not for *S*_*U*_ = 50, *S*_*Y*_ = 5 and *R* = 1 (i.e. a small sample size), in which GLM-2SRI was more biased than GLM-2SPS.

**Figure 5:**
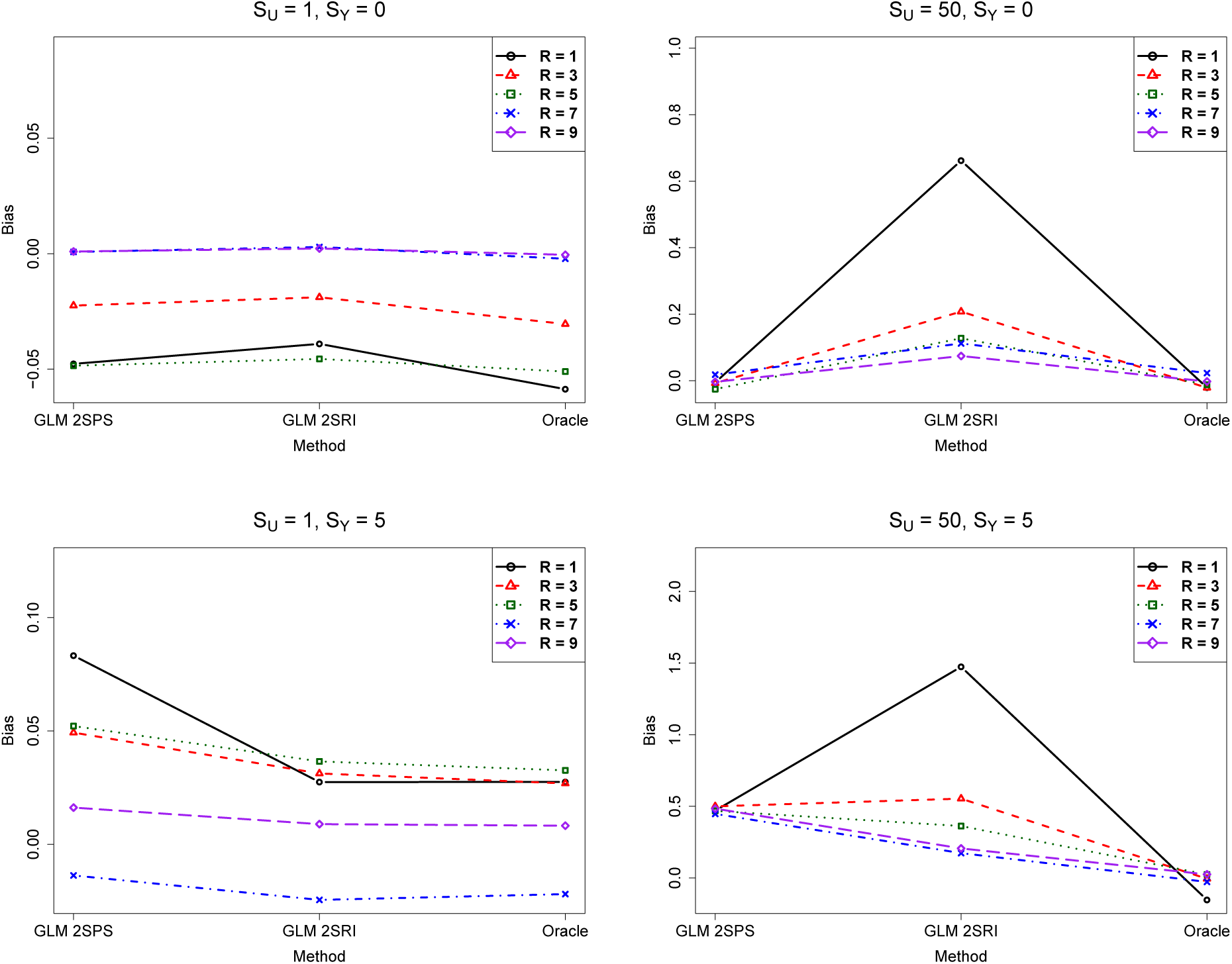
Simulations with the two-sample approaches: biases of various methods.

#### 3.2.4 Comparison of the original and corrected standard error estimates

For GLM-2SRI and LM-2SPS, we compared their original and corrected standard error estimates. Figure 6 shows that, first, with a small *S*_*U*_, the corrected SEs were slightly larger than the original SEs, but the difference became much smaller as the sample size *R* increased. On the other hand, under more severe confounding with a larger *S*_*U*_, the corrected SEs were always much larger than the original one for GLM-2SRI, regardless of the sample size; in comparison, the differences between the two for LM-2SPS were much smaller, and tended to disappear as the sample size increased, across all the simulation settings (also see Supplementary figures). These results suggest that it is perhaps necessary to use the corrected SE for GLM-2SRI, but not so for LM-2SPS (especially for large sample sizes).

**Figure 6:**
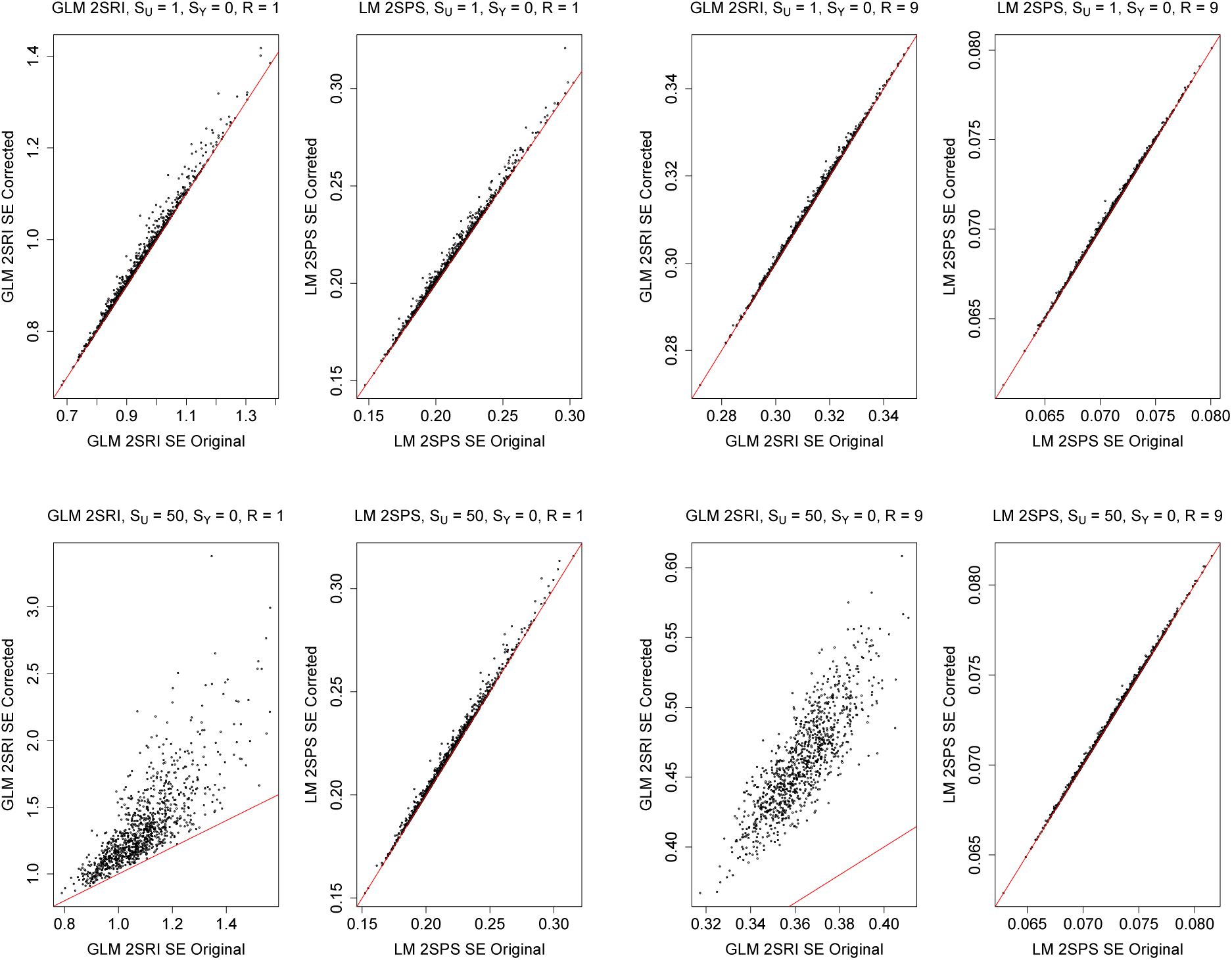
Simulations with the two-sample approaches: comparison of the original and corrected standard error estimates.

#### 3.2.5 Results For Other Settings

We have only shown some representative results for the simulation set-ups with *S*_*Y*_ = 0 or 5, and *S*_*U*_ = 1 or 50. The results for other set-ups are shown in the Supplementary Materials.

### 3.3 Simulation Results: One-sample Approaches

We reached similar conclusions on the relative performance of the four methods for the one-sample case as for the two-sample case shown earlier, except that GLM-2SRI performed better and were more in par with GLM-2SPR and LM-2SPR. Again LM-2SRI yielded inflated Type I errors. The detailed results are shown in the Supplementary Materials. It is noted that the three one-sample approaches performed well was presumably due to our “weak” selection of SNPs: we only selected 10 SNPs from 30 candidate SNPs as IVs; if more candidate SNPs were included while more or less SNPs were allowed to be selected as IVs, the effects of the selection bias would be larger as shown for the real data analysis.

## 4 Conclusions and Discussion

In summary, presumably due to selection bias, one-sample approaches may lead to inflated type I errors and thus are not recommended. Among the two-sample approaches, GLM-2SPS, followed by LM-2SPS, performed best; note that these two methods are used as default in practice for TWAS. Two-sample GLM-2SRI performed well if the sample size was large enough; otherwise it could be conservative, and even with large biases. In contrast, two-sample LM-2SRI did not perform well across all simulations, and should not be used.

Why did 2SPS perform well, even better than 2SRI for non-linear logistic regression model in Stage 2 in our study? Does this contradict the general theory of 2SRI? A quick answer to the second question is no. In retrospect, the reason for the first question is simple: it is related to the currently well accepted practice of applying a linear model to a binary trait in GWAS, because a linear model can approximate well the corresponding logistic (or other non-linear) regression model due to the small effect sizes of SNPs (Zhao, Wang, Hemani, Bowden & Small, 2019). Furthermore, in 2SRI, with the high correlation between the observed gene expression (*Y*_*i*_ in our notation) and the residual/estimated confounding (*Û*_*i*_ = *Y*_*i*_ − *Ŷ*_*i*_) due to the often low predictivity of a gene’s expression level by its cis-SNPs, fitting the Stage-2 model in 2SRI requires a larger sample size for it to perform well. Finally, it is not feasible to even apply 2SRI with two separate samples of eQTL and GWAS data, because of no observed gene expression levels (*Y*_*i*_’s) in the typical GWAS data.

We also note that in practice of using TWAS, the statistical uncertainty (i.e. estimation error) in imputing gene expression in Stage 1 is ignored. Although this uncertainty can be taken account using the corrected SE estimator, our numerical study suggested its negligible effects. Hence again the usual practice with TWAS of no correction appears to be fine.

We emphasize that our main conclusion (that the standard TWAS performs well) holds only under the conditions with the large sample size and small effect sizes of genetic variants on complex traits and common diseases. Otherwise, for example, in extensions of TWAS to molecular traits or other endophenotypes (Xu, Wu, Wei, Pan & Alzheimer’s Disease Neuroimaging Initiative, 2017b; Wu et al., 2018), on which genetic variants (or other IVs) may have much larger effect sizes, cautions should be taken: 2SPS as adopted in the standard TWAS may not be even consistent for a non-linear model in Stage 2.

There are other limitations with the current study, including the following two important and challenging issues. First, instead of OLS estimation, penalized regression methods, such as Lasso or elastic net (Zou & Hastie, 2005), or Bayesian methods, are often used in the first stage for TWAS in practice (Gamazon et al., 2015; Gusev et al., 2016). The benefits include selecting relevant SNPs as valid IVs, avoiding biases of weak IVs, and obtaining better estimates to impute gene expression better, which presumably would lead to better inference (e.g. more precise estimates and higher power) for the causal parameter in the second stage. Some large sample properties, e.g. square-root-*n* consistency, of such Lasso or post-Lasso procedures, have been established for sparse models under suitable conditions (Belloni et al., 2012). However, for finite (especially small) samples, these methods may *not* yield imputed gene expression leve;s orthogonal to or uncorrelated with the confounders, leading to possibly biased inference on the causal parameter with 2SPS in Stage 2. An alternative approach is jackknife instrumental variable estimation, which predicts/imputes each observation *i* by fitting a Stage 1 model with other observations after excluding observation *i*, but it is computationally intensive (Angrist, Imbens, & Krueger, 1999; Hansen & Kozbur, 2014). Furthermore, given the relatively large sample size in TWAS, it is not clear how biased the causal parameter inference would be if no correction is applied. Second, in our simulations, we only considered “weak selection” of IVs: we selected 10 SNPs as IVs from 30 candidate ones containing 3 causal SNPs (i.e. 3 valid IVs), which was expected to select at least one valid IV with a high probability while having a relatively small selection bias. The more difficult cases with few or no valid IVs, or more practically as in TWAS with a larger set of candidate SNPs/IVs, may render it necessary to use penalized regression or other more sophisticated methods in Stage 1, introducing some challenges as discussed earlier. More work is needed.

## Supporting Information

In the Supplementary Materials, we provide more analysis results for the ADNI data (Figures S1-S2), simulations for the two-sample approaches (Figures S3-S44), and simulations for the one-sample approaches (Figures S45–S86).

## Acknowledgements

We thank the reviewers for many helpful comments and suggestions. WP would like to thank Dr. Todd MacKenzie for first introducing 2SRI to him. This work was supported by NIH grants R21AG057038, R01HL116720, R01GM113250, R01GM126002 and R01HL105397, and by the Minnesota Supercomputing Institute at the University of Minnesota.

