## Supplementary_materials for "Some Statistical Consideration in Transcriptome-Wide Association Studies"

July 22, 2019

[3]Data used in preparation of this article were obtained from the Alzheimer's Disease Neuroimaging Initiative (ADNI) database (adni.loni.usc.edu). As such, the investigators within the ADNI contributed to the design and implementation of ADNI and/or provided data but did not participate in analysis or writing of this report. A complete listing of ADNI investigators can be found at: http://adni.loni.usc.edu/wp-content/uploads/how_to_apply/ADNI_Acknowledgement_List_Sep23.pdf.

### 1 More About Real Data Results

As we mentioned in section 3.1, we perform $F$-tests in the first stage to ensure the first IV assumption is satisfied. For $p$-value cutoff 0.05, there are 9102 genes with $p$-value smaller than it, we show Q-Q plots of them in Figure S1; for $p$-value cutoff 0.1, there are 10564 genes with $p$-value smaller than it, we show Q-Q plots of them in Figure S2.

Figure S1: The ADNI data analysis: Q-Q plots of the obtained p-values of 9102 genes from each method versus the expected p-values under the null hypothesis of no association

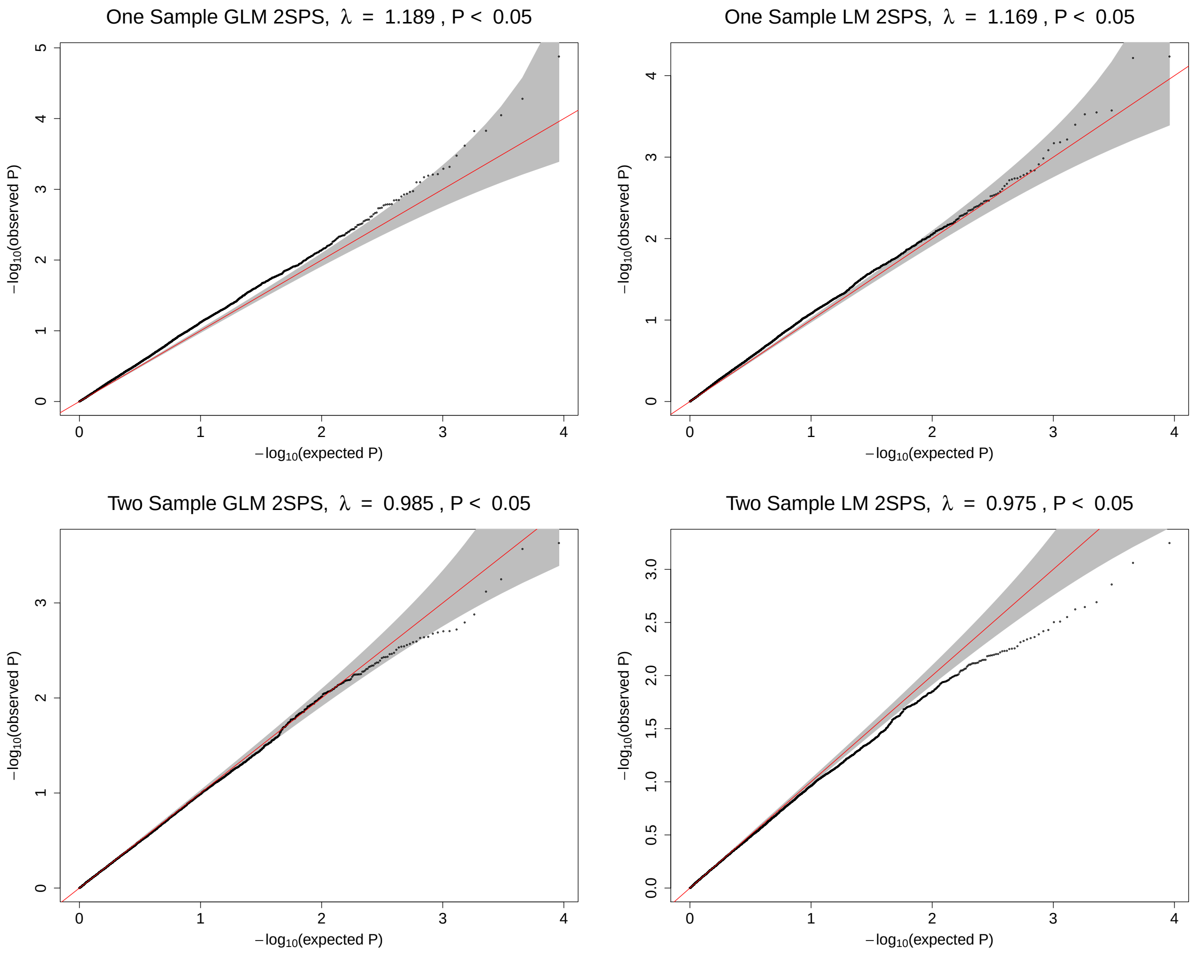

Figure S1 (Cont.): The ADNI data analysis: Q-Q plots of the obtained p-values of 9102 genes from each method versus the expected p-values under the null hypothesis of no association

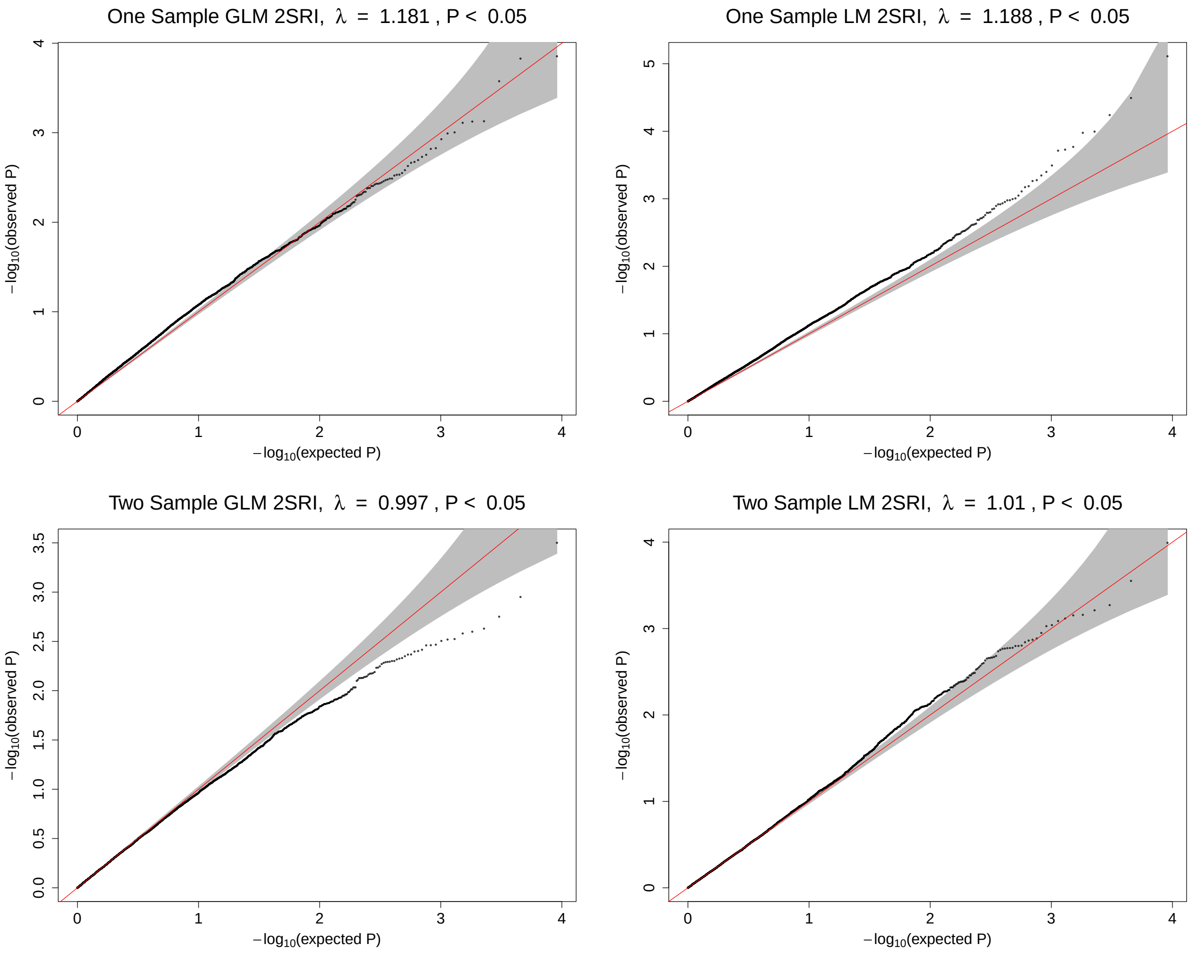

Figure S2: The ADNI data analysis: Q-Q plots of the obtained p-values of 10564 genes from each method versus the expected p-values under the null hypothesis of no association

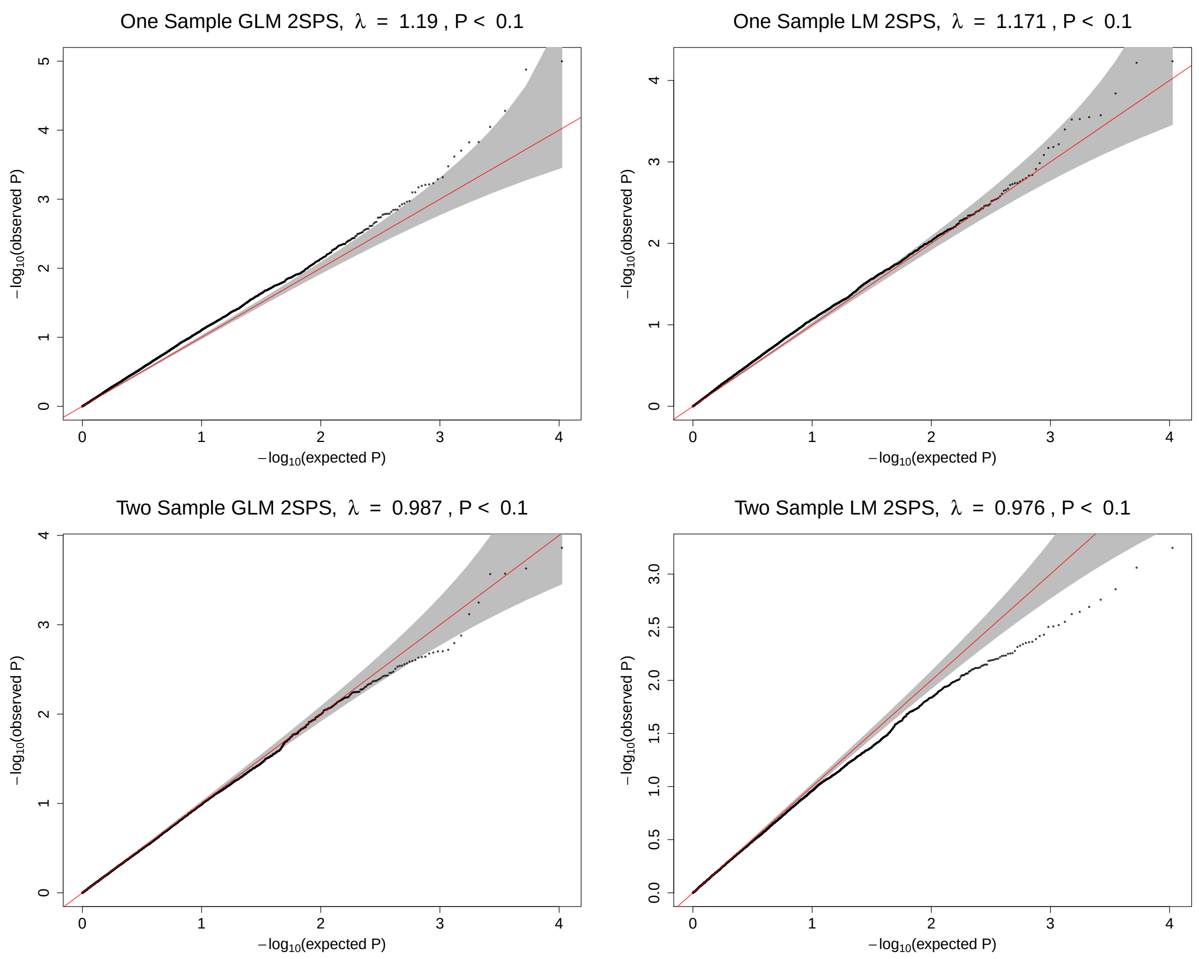

Figure S2 (Cont.): The ADNI data analysis: Q-Q plots of the obtained p-values of 10564 genes from each method versus the expected p-values under the null hypothesis of no association

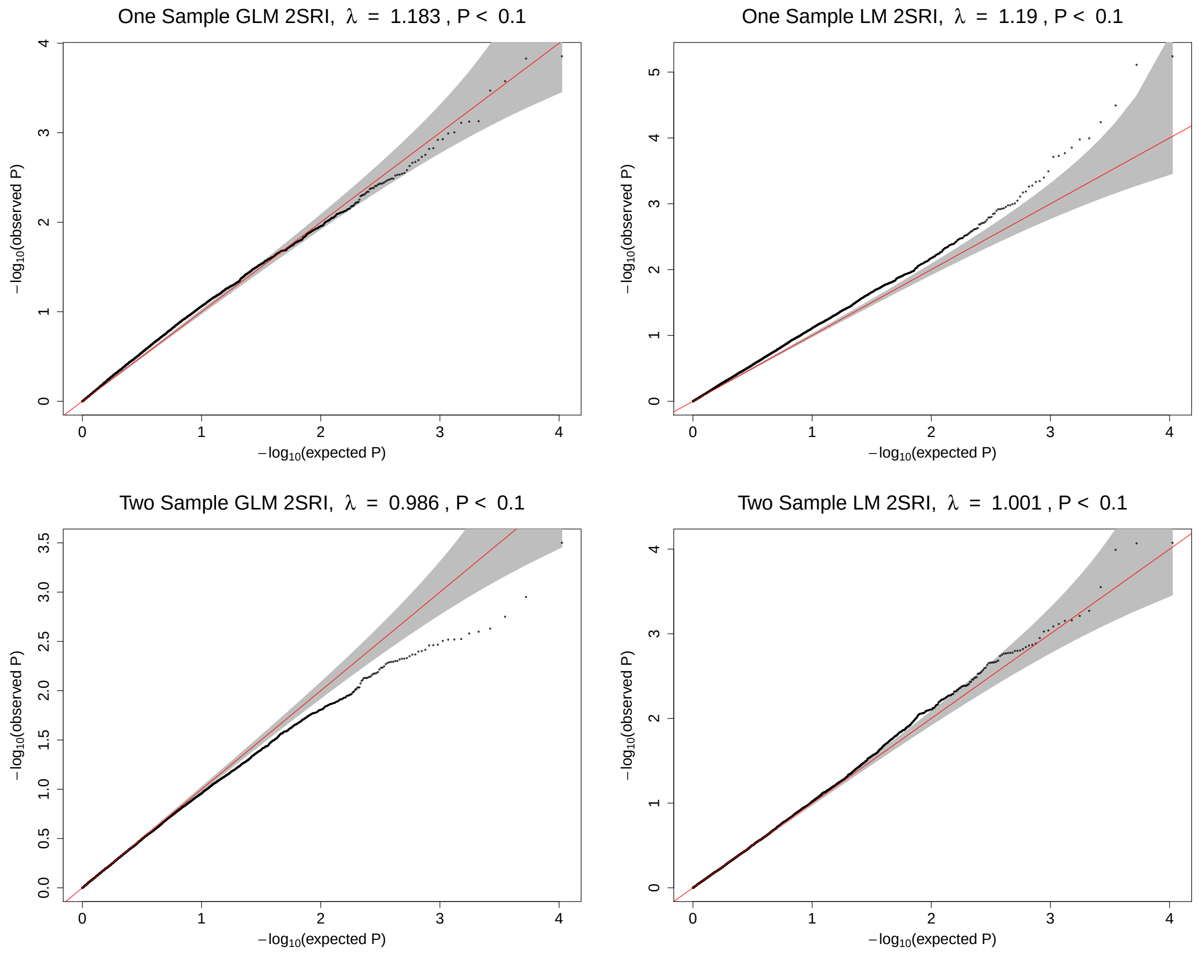


#### 2 More About Simulation Results

In section 3.2, we showed part of our simulation results about comparing Type I Error, Power, Bias, and Standard Error for our methods. Here we show the full results for combinations $\{(R, S_Y, S_U) | R = 1,3,5,7,9; S_Y = 0,1,2,3,4,5; S_U = 1,10,20,30,40,50\}$.

#### 2.1 Two-sample Approaches

##### 2.1.1 Type I Error

Figure S3: Simulations with the two-sample approaches: empirical Type I error rates of various methods.

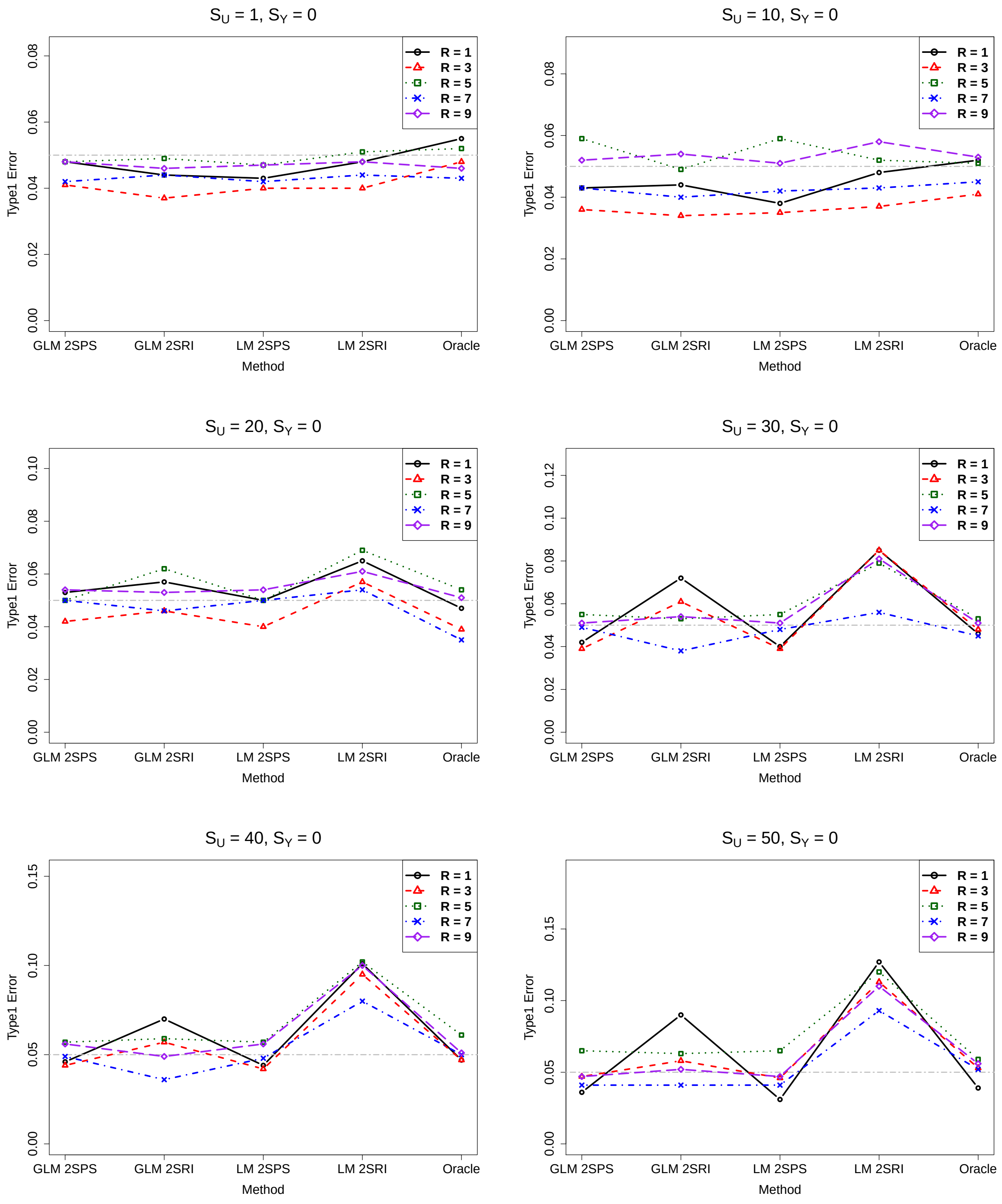

##### 2.1.2 Power

Figure S4: Simulations with the two-sample approaches: empirical power of various methods, $S_Y = 1$.

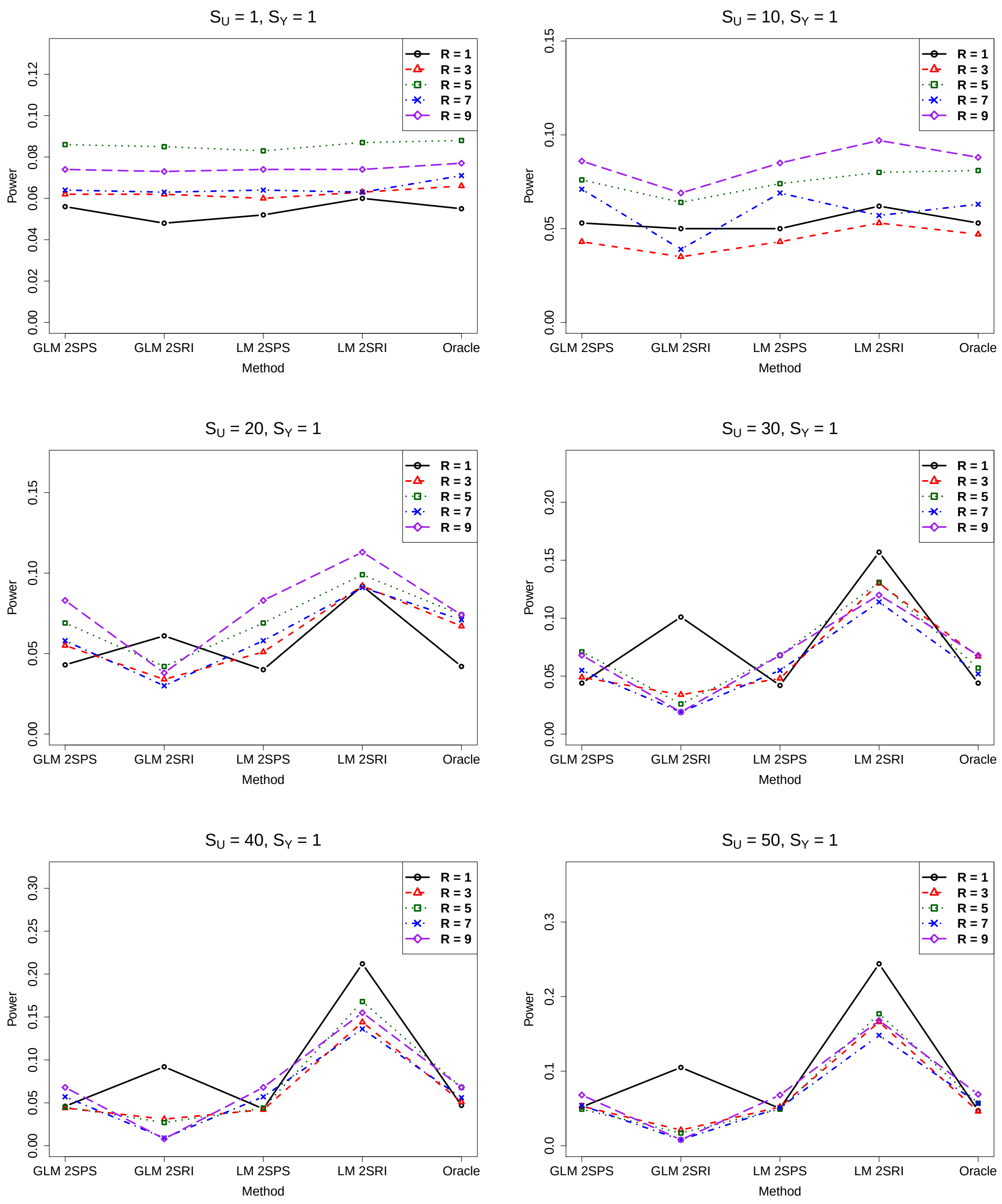

Figure S5: Simulations with the two-sample approaches: empirical power of various methods, $S_Y = 2$.

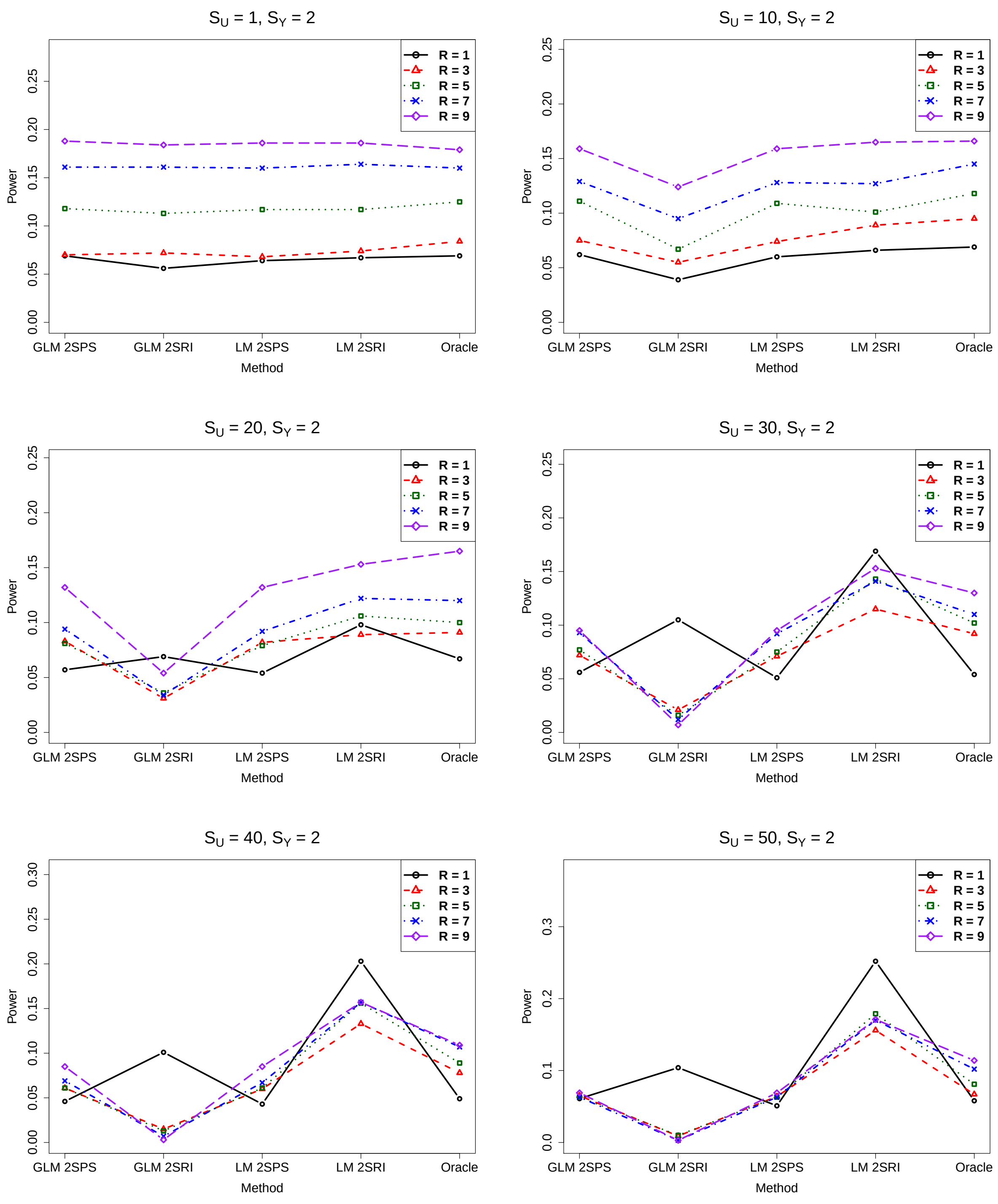

Figure S6: Simulations with the two-sample approaches: empirical power of various methods, $S_Y = 3$.

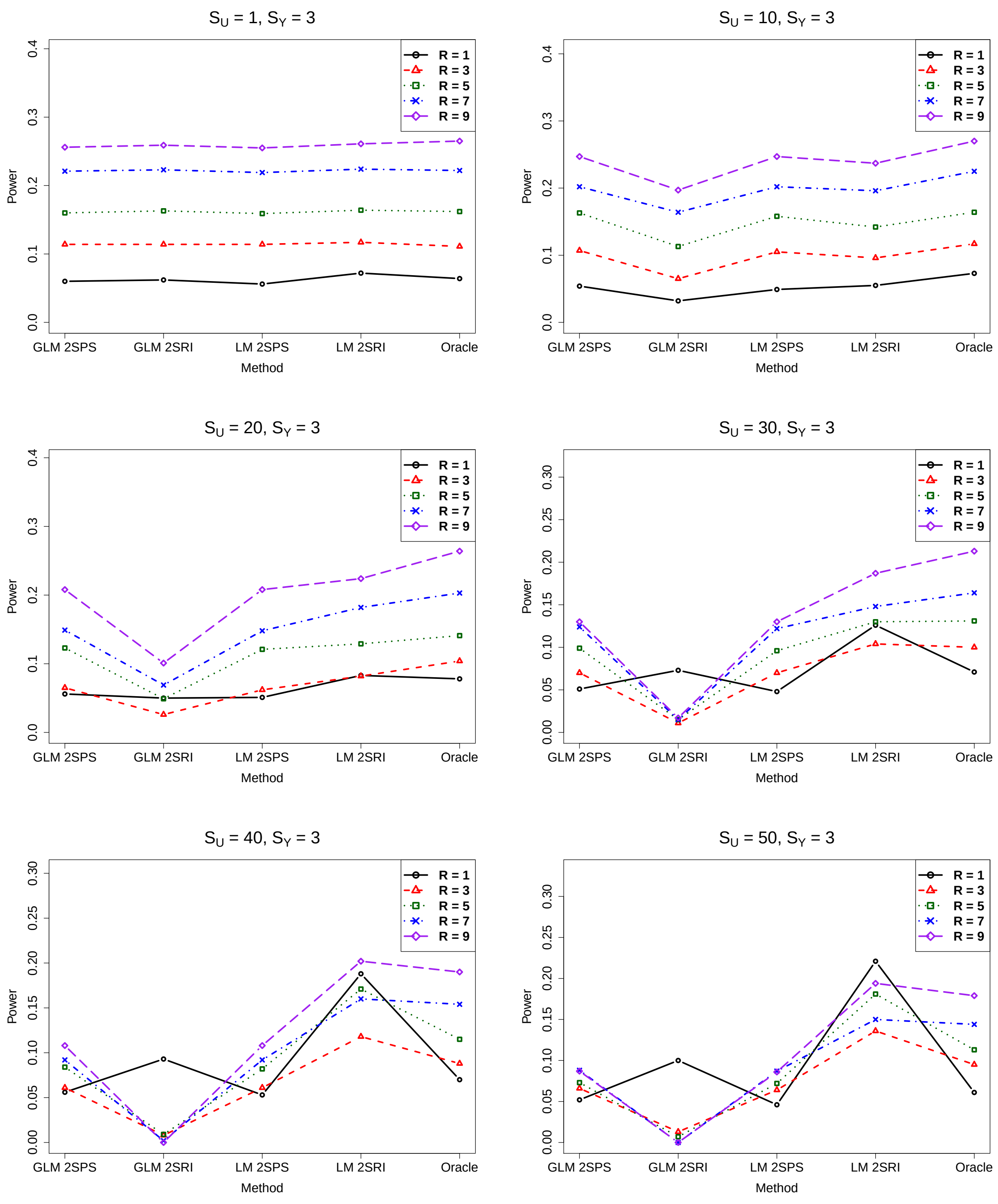

Figure S7: Simulations with the two-sample approaches: empirical power of various methods, $S_Y = 4$.

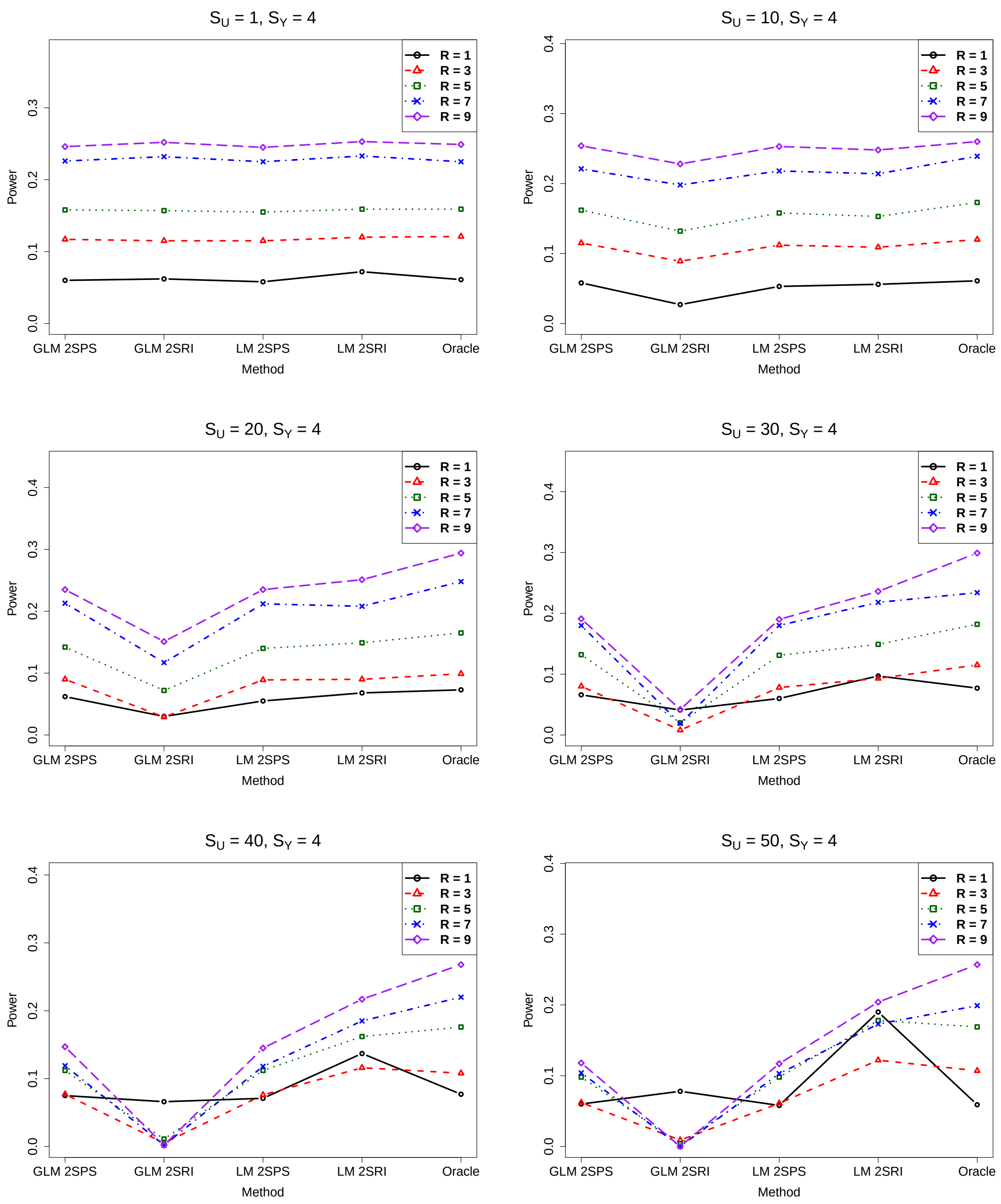

Figure S8: Simulations with the two-sample approaches: empirical power of various methods, $S_Y = 5$.

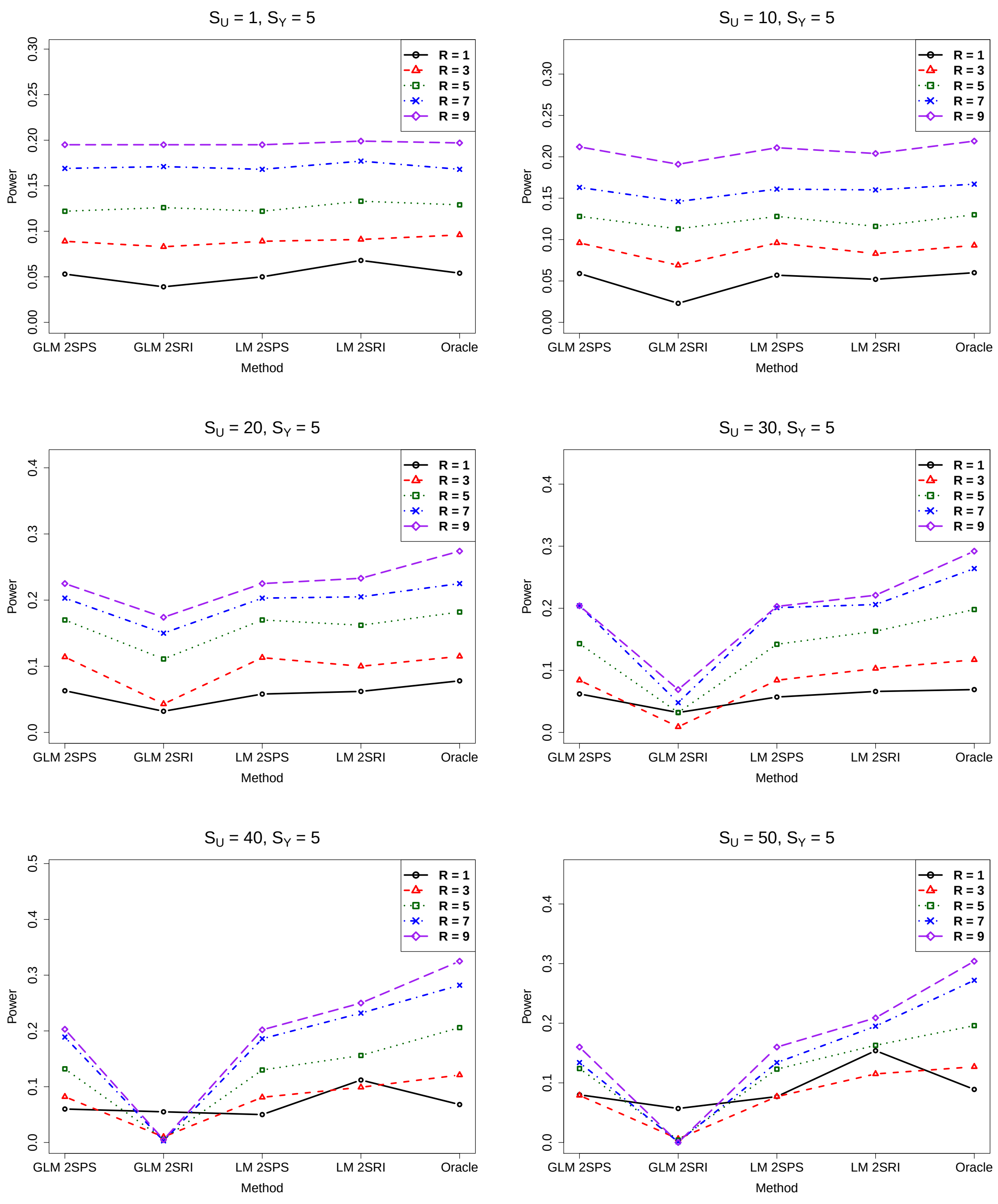

##### 2.1.3 Bias

Figure S9: Simulations with the two-sample approaches: biases of various methods, $S_Y = 0$

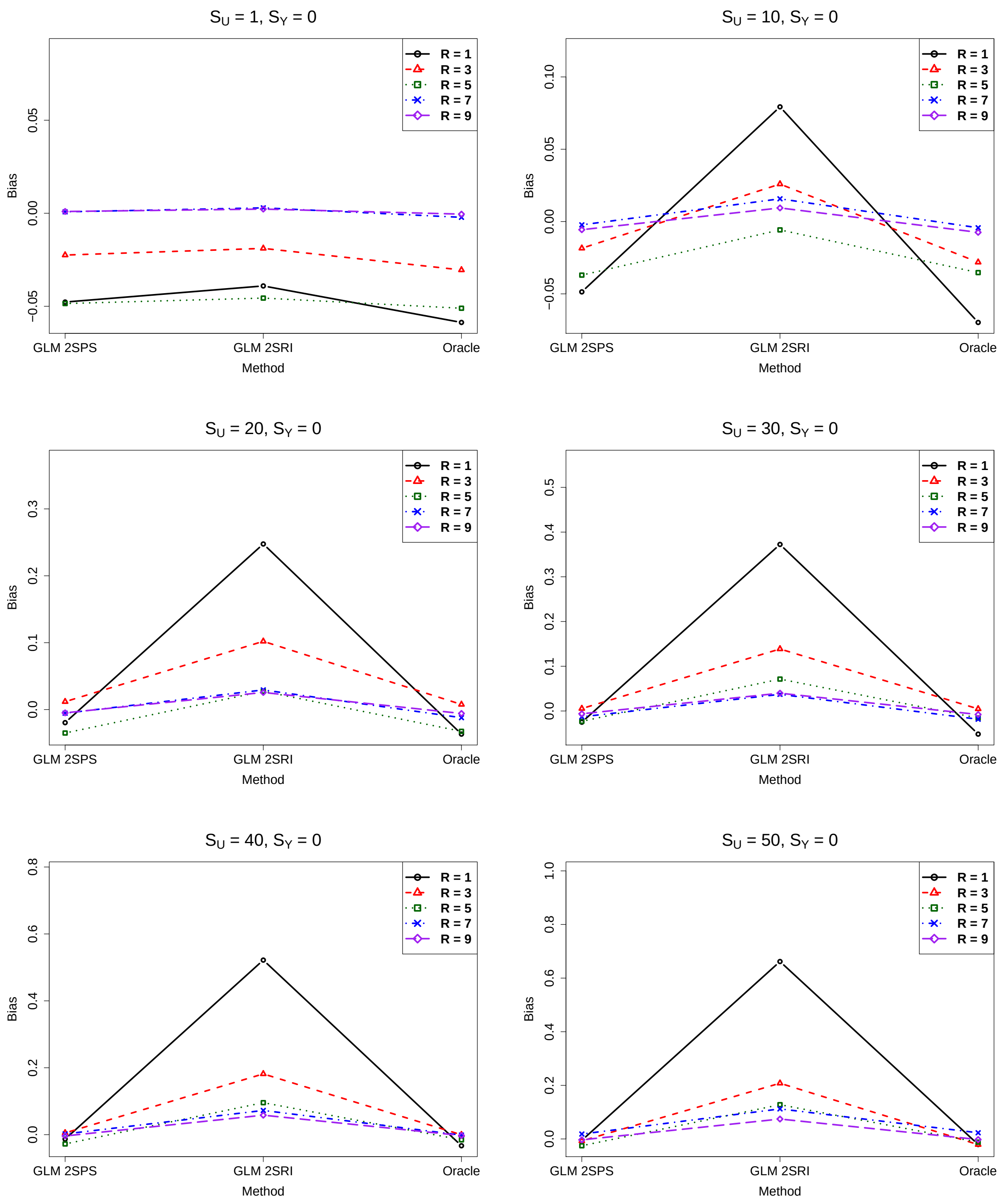

Figure S10: Simulations with the two-sample approaches: biases of various methods, $S_Y = 1$

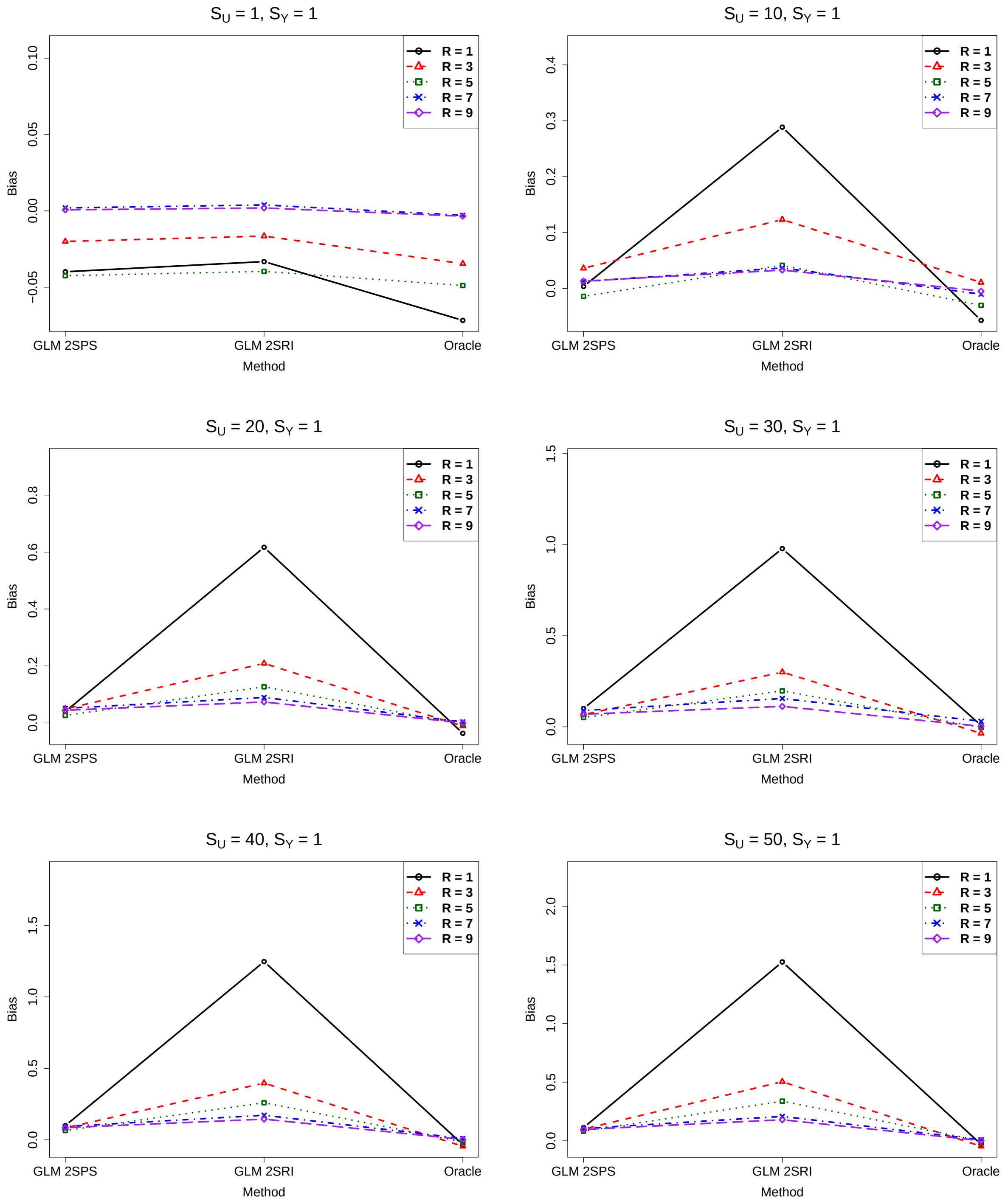

Figure S11: Simulations with the two-sample approaches: biases of various methods, $S_Y = 2$

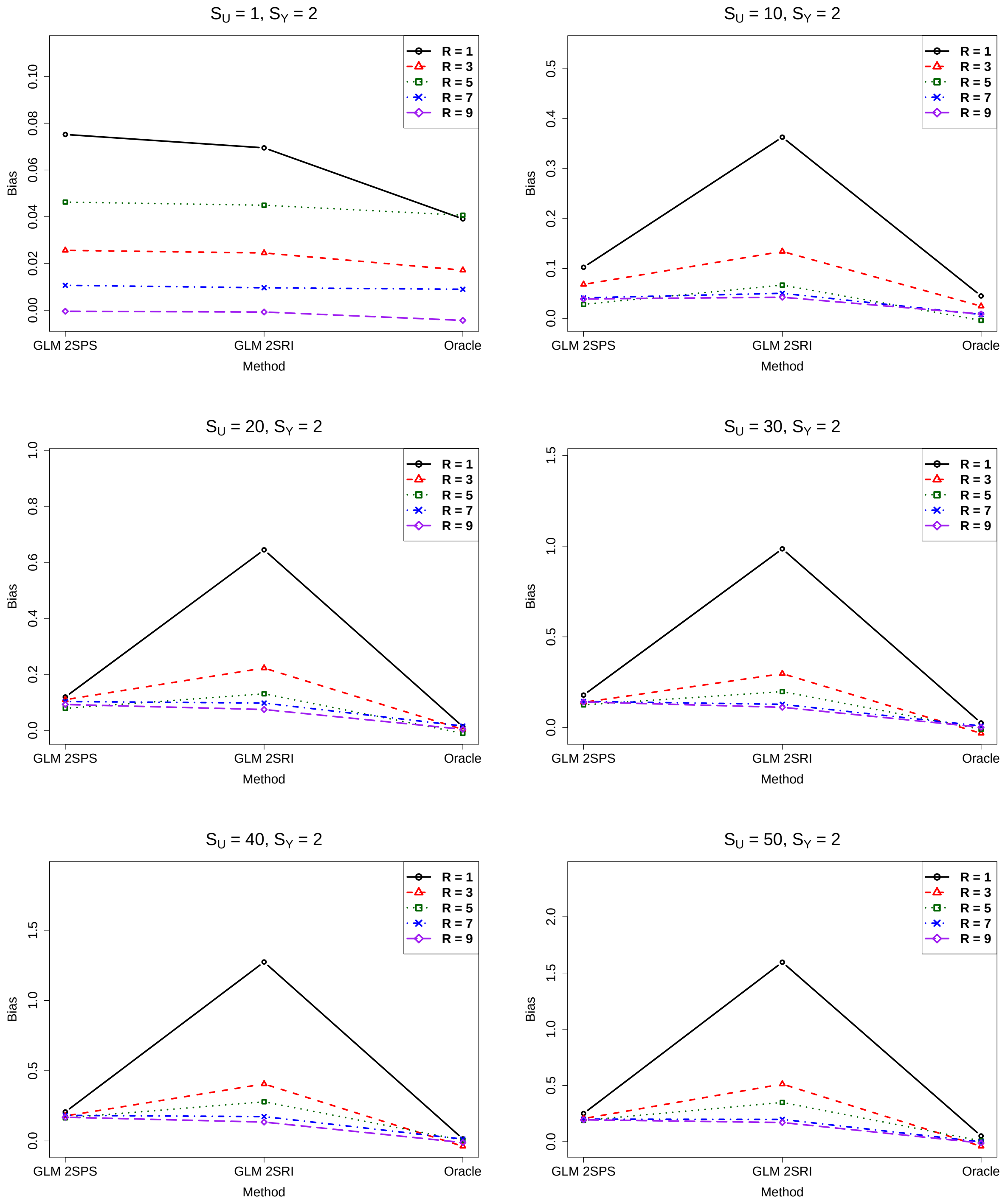

Figure S12: Simulations with the two-sample approaches: biases of various methods, $S_Y = 3$

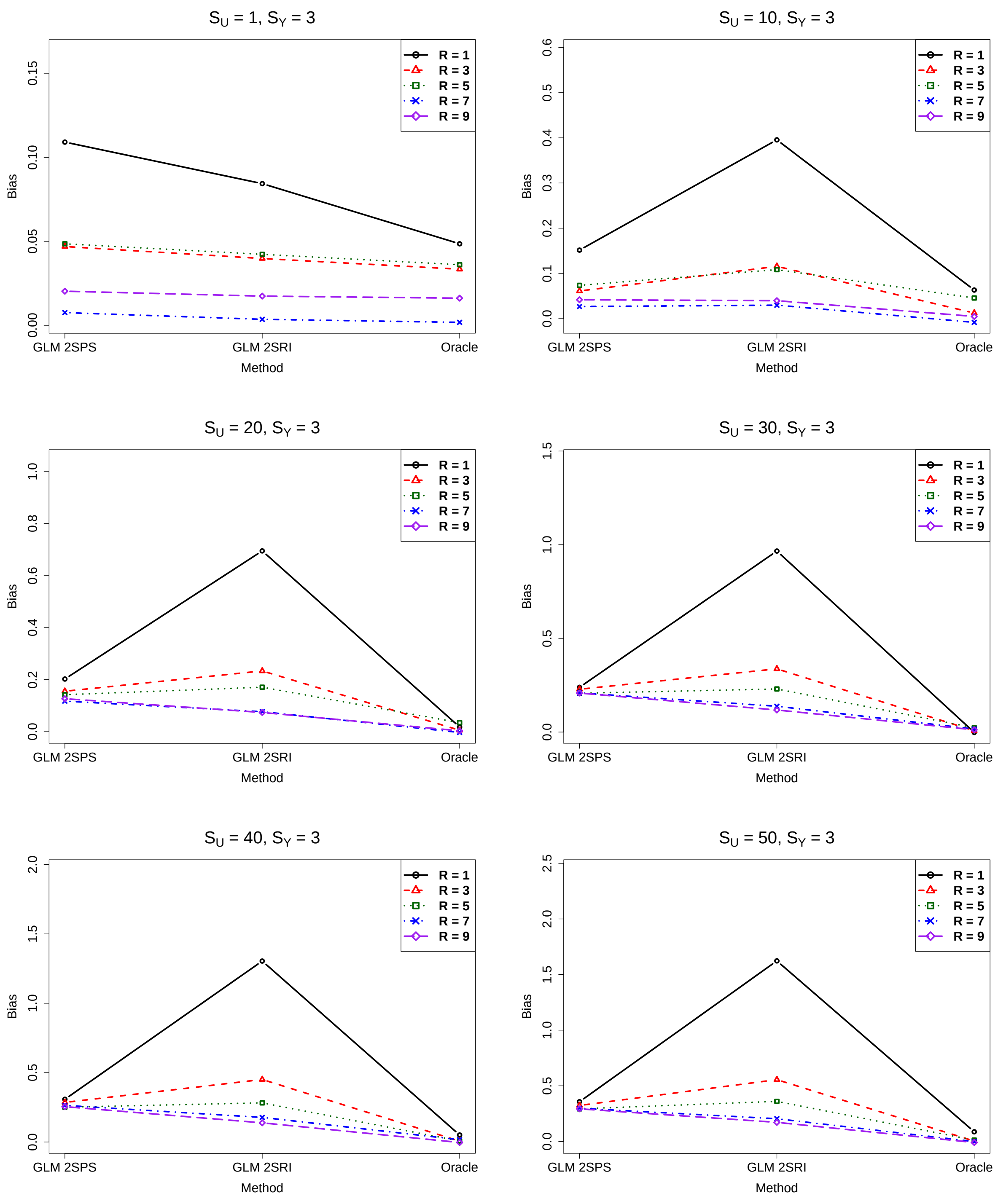

Figure S13: Simulations with the two-sample approaches: biases of various methods, $S_Y = 4$

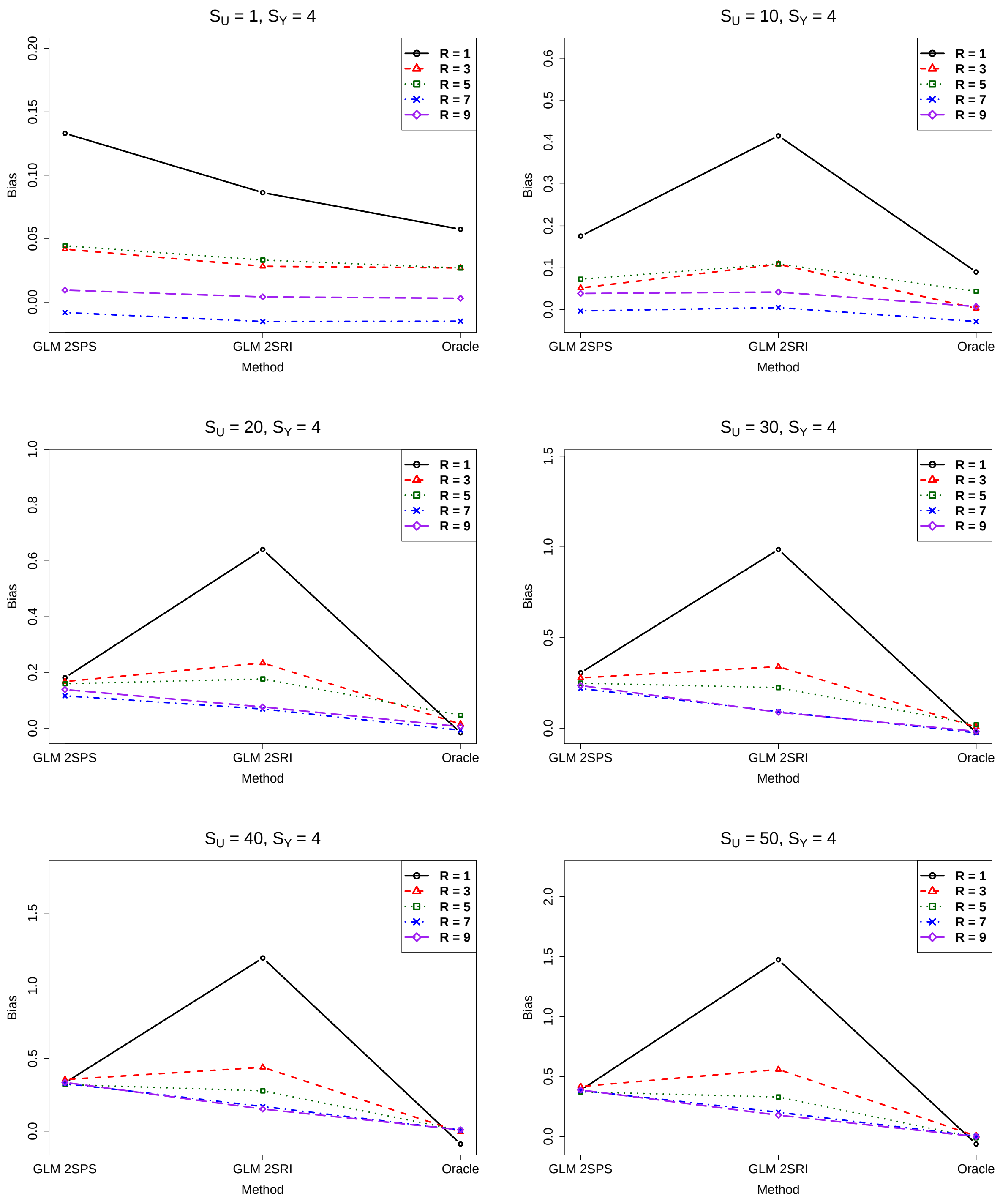

Figure S14: Simulations with the two-sample approaches: biases of various methods, $S_Y = 5$

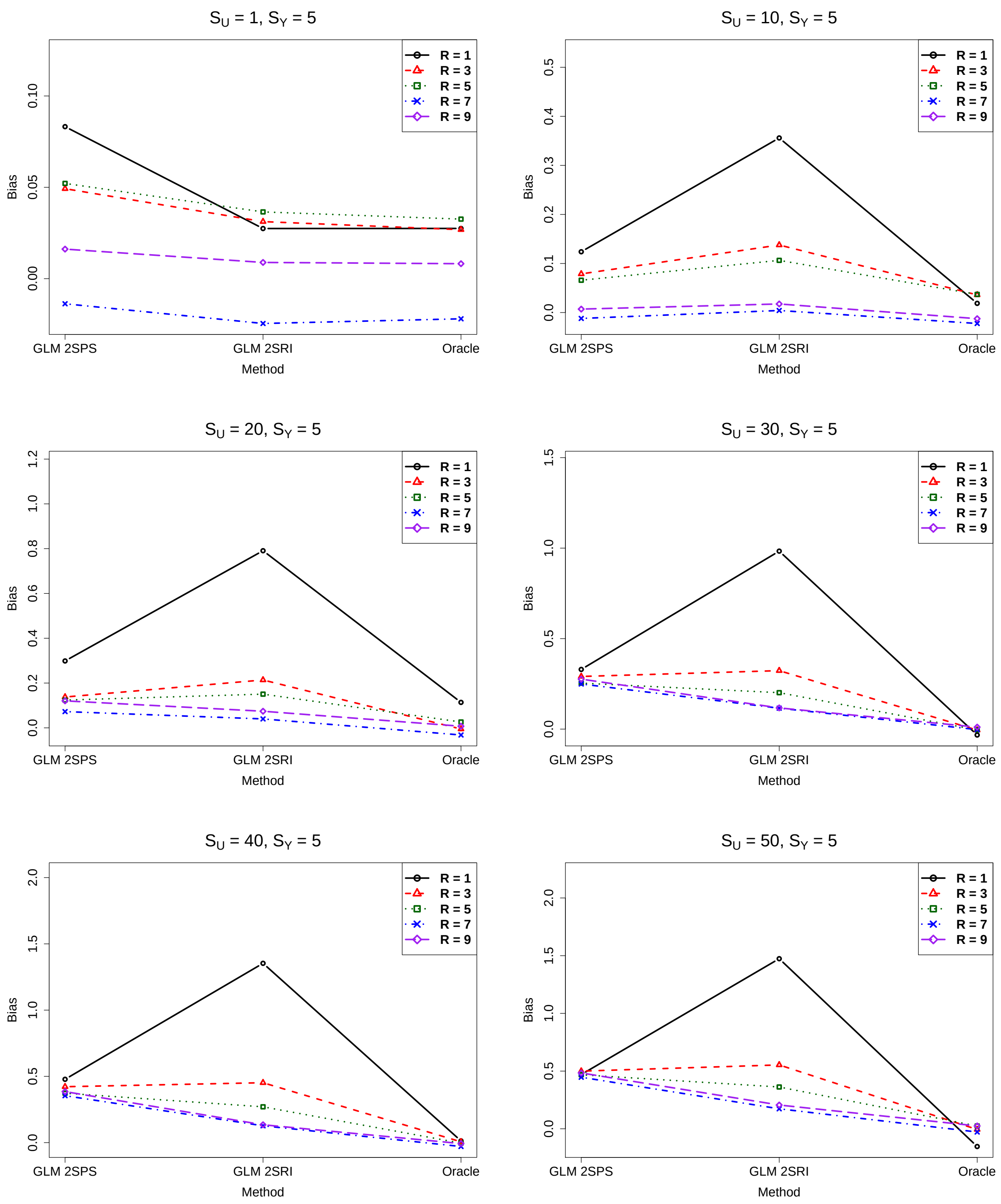

##### 2.1.4 Comparison of the original and corrected standard error estimates

Figure S15: Simulations with the two-sample approaches: comparison of the original and corrected standard error estimates, $S_Y = 0, R = 1$.

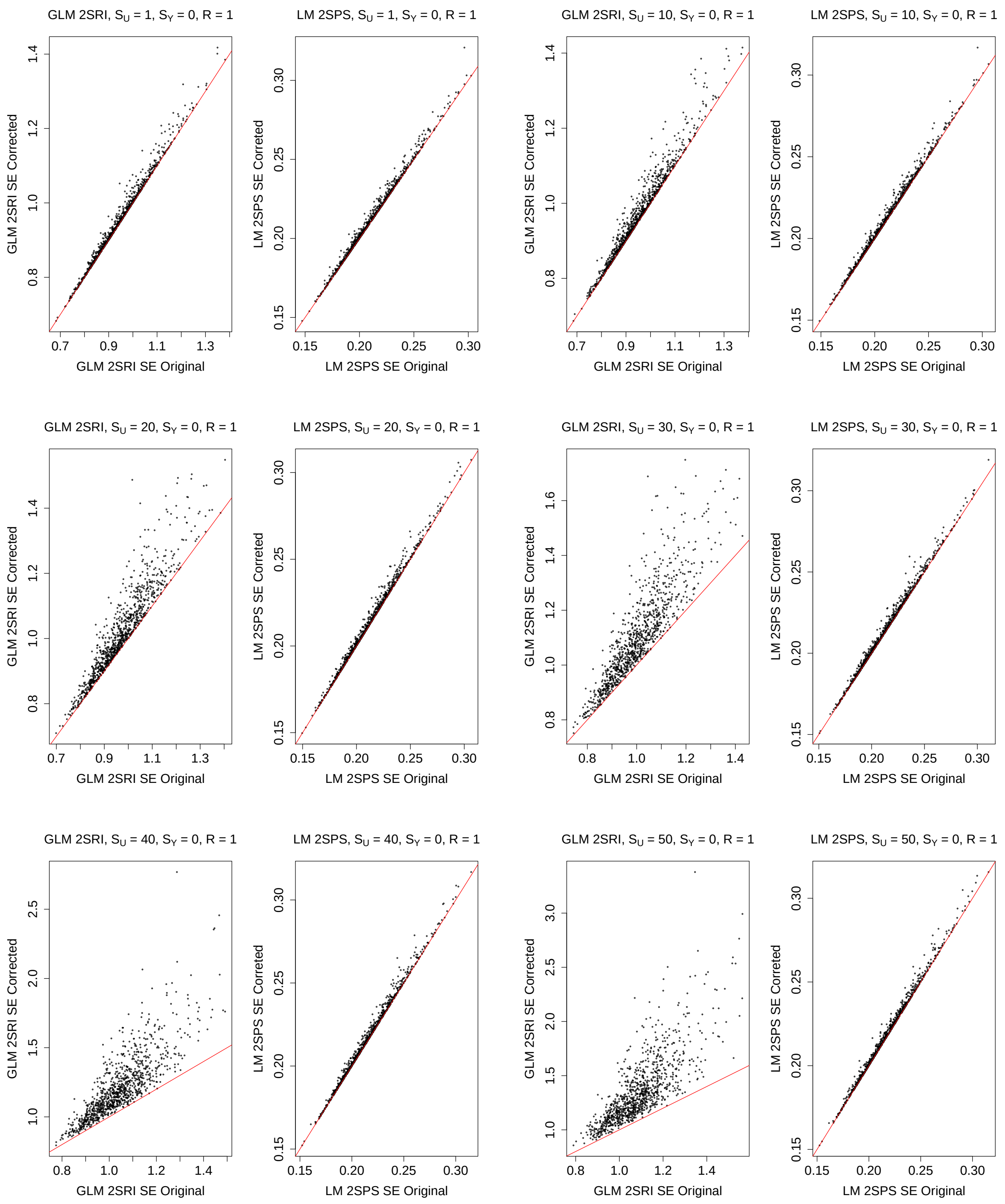

Figure S16: Simulations with the two-sample approaches: comparison of the original and corrected standard error estimates, $S_Y = 1, R = 1$.

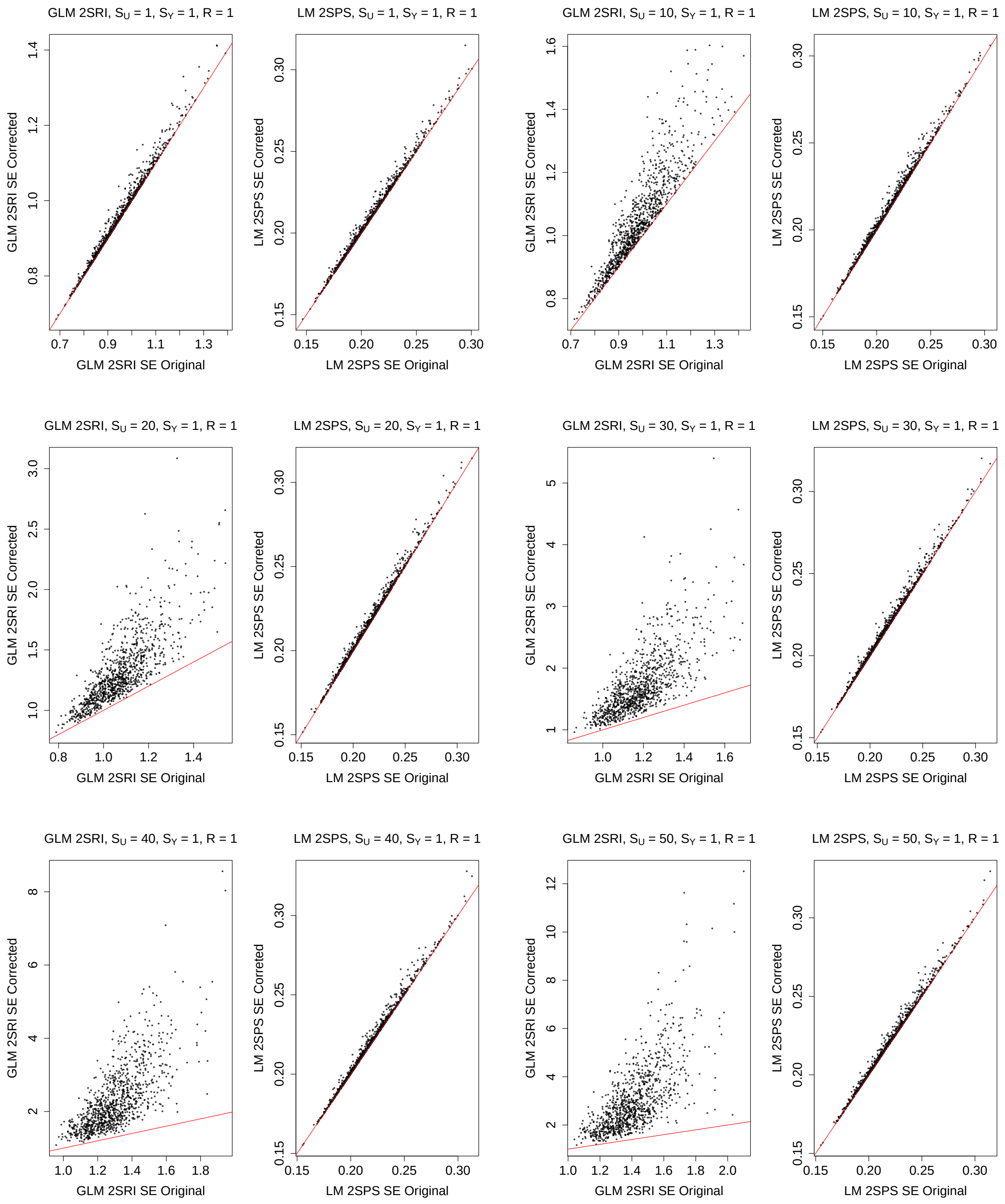

Figure S17: Simulations with the two-sample approaches: comparison of the original and corrected standard error estimates, $S_Y = 2, R = 1$.

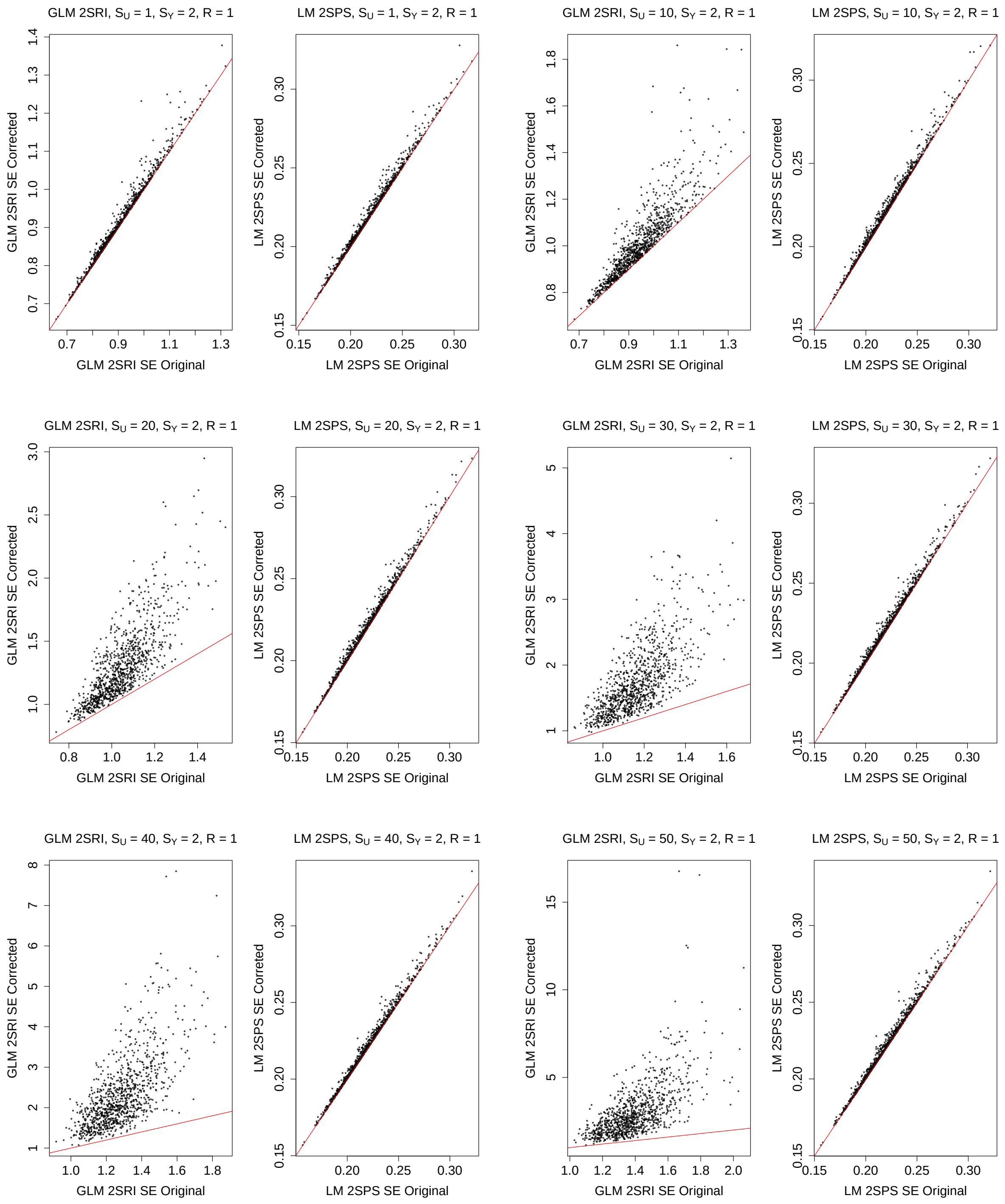

Figure S18: Simulations with the two-sample approaches: comparison of the original and corrected standard error estimates, $S_Y = 3, R = 1$.

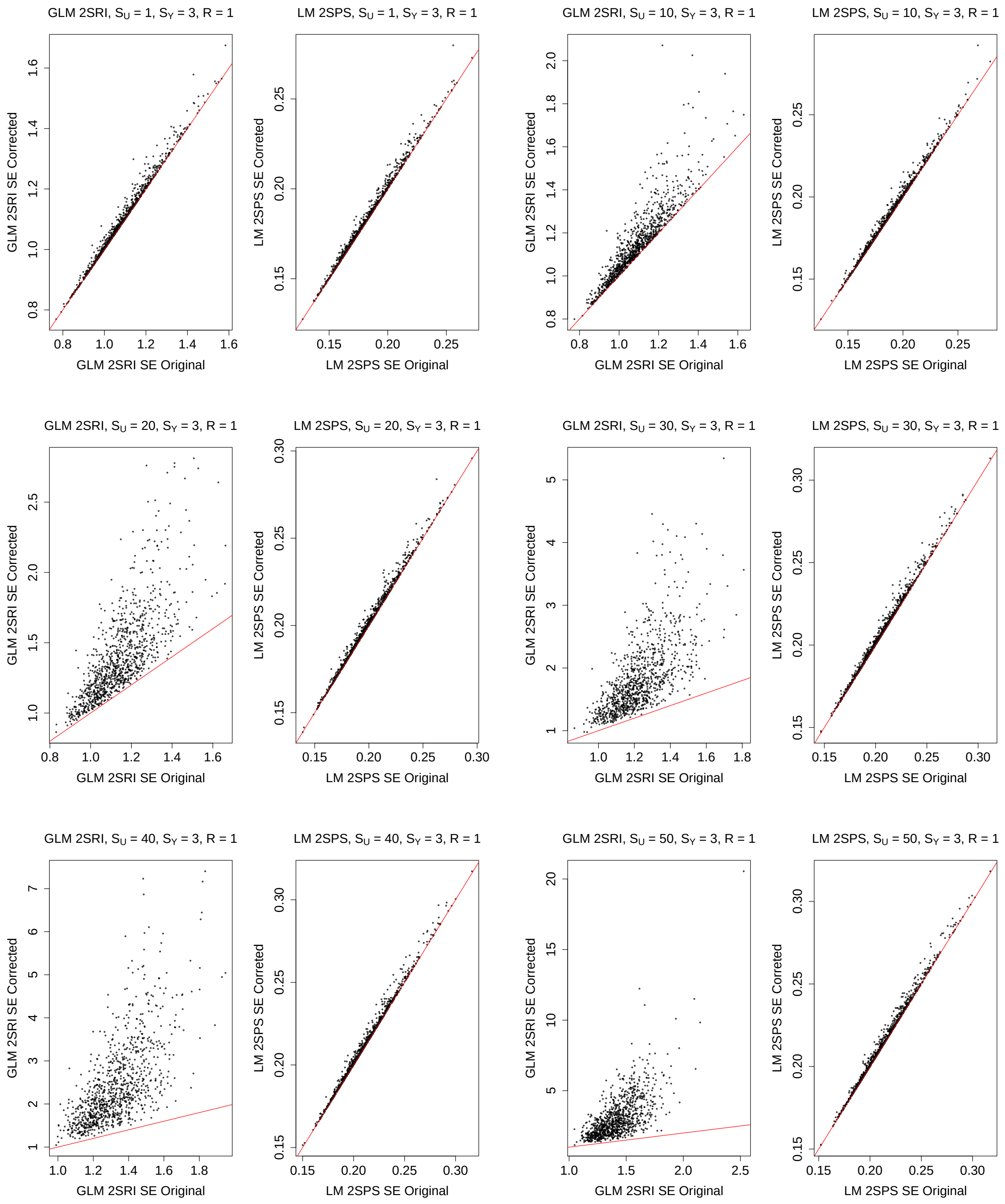

Figure S19: Simulations with the two-sample approaches: comparison of the original and corrected standard error estimates, $S_Y = 4, R = 1$.

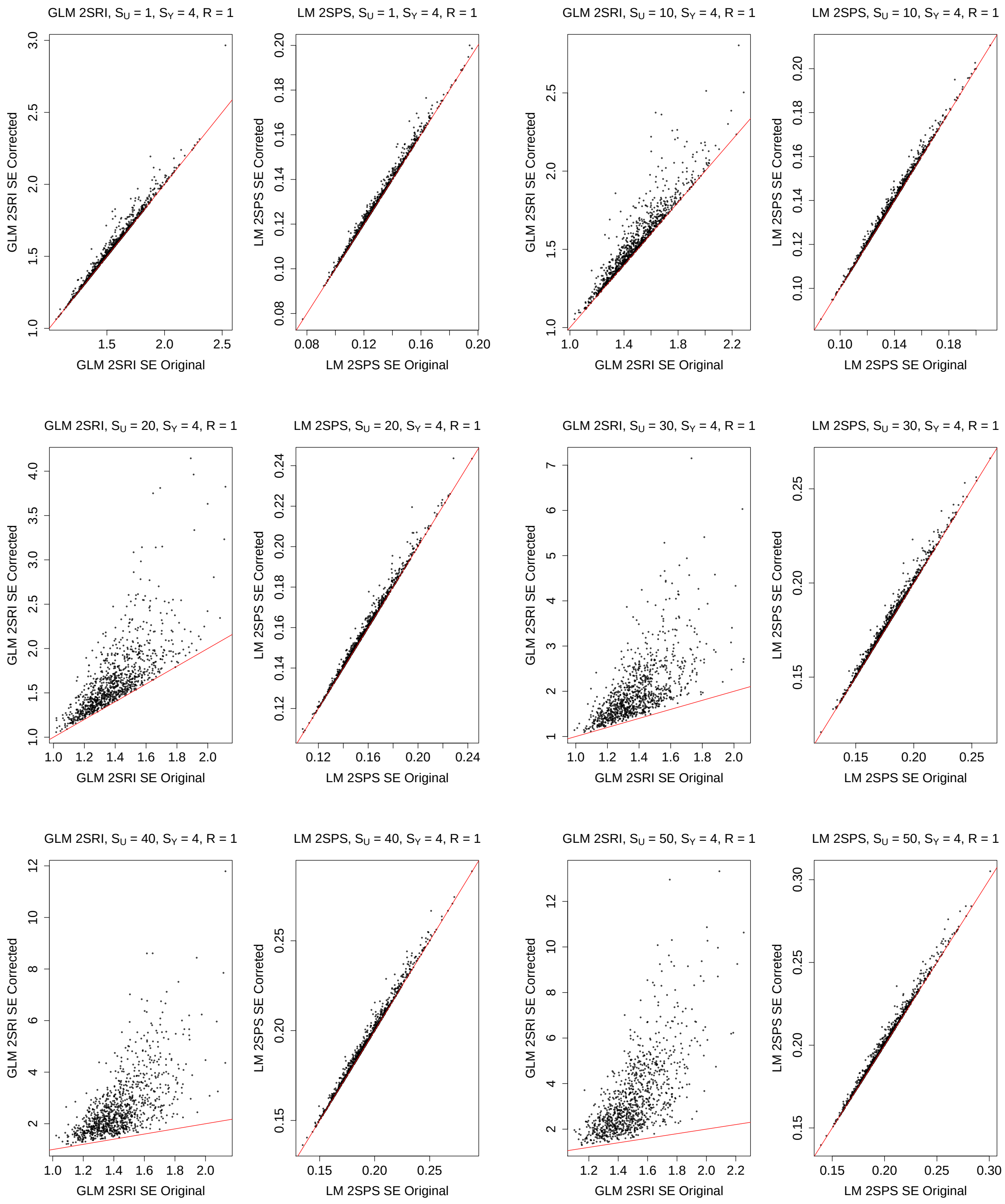

Figure S20: Simulations with the two-sample approaches: comparison of the original and corrected standard error estimates, $S_Y = 5, R = 1$.

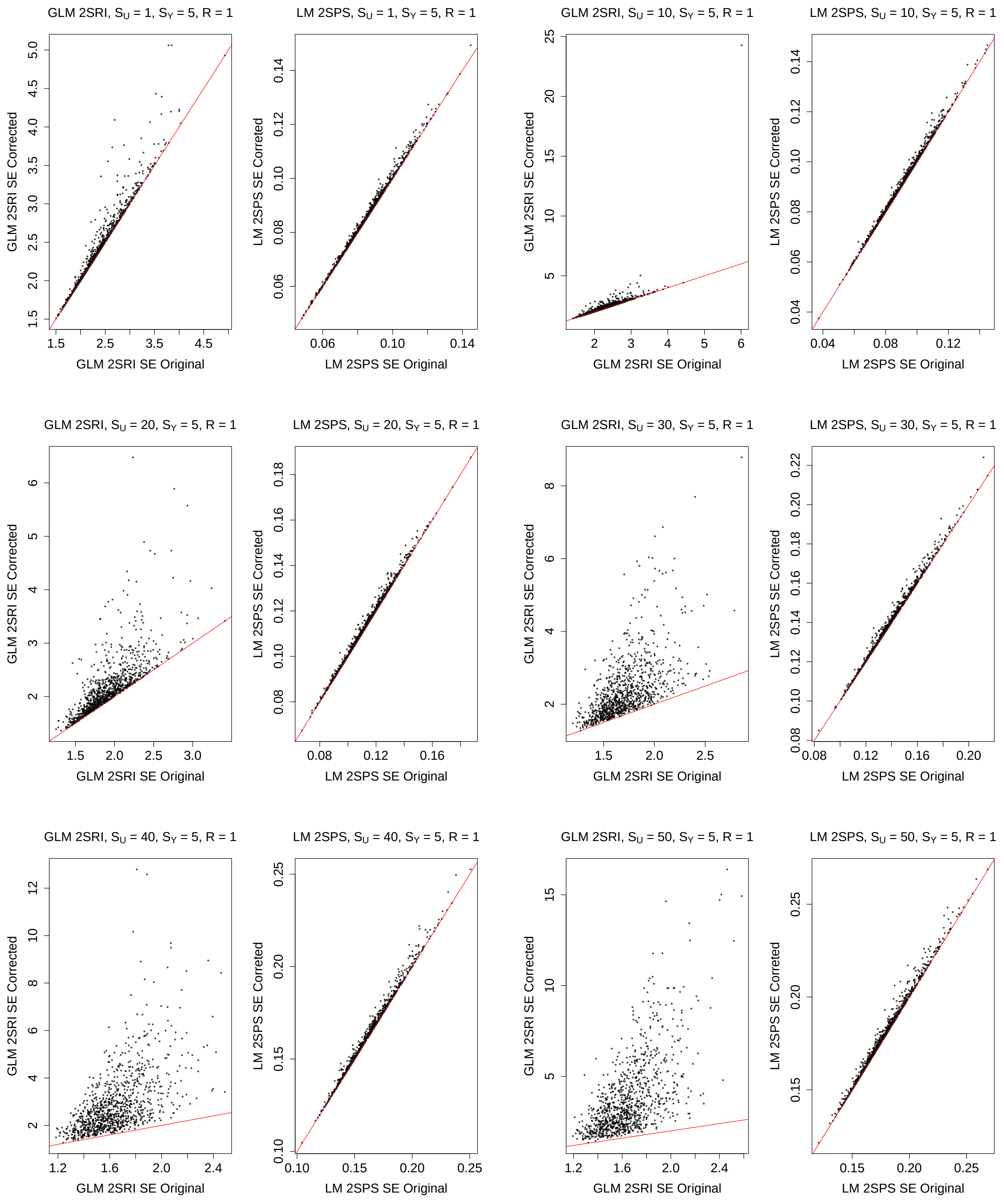

Figure S21: Simulations with the two-sample approaches: comparison of the original and corrected standard error estimates, $S_Y = 0, R = 3$.

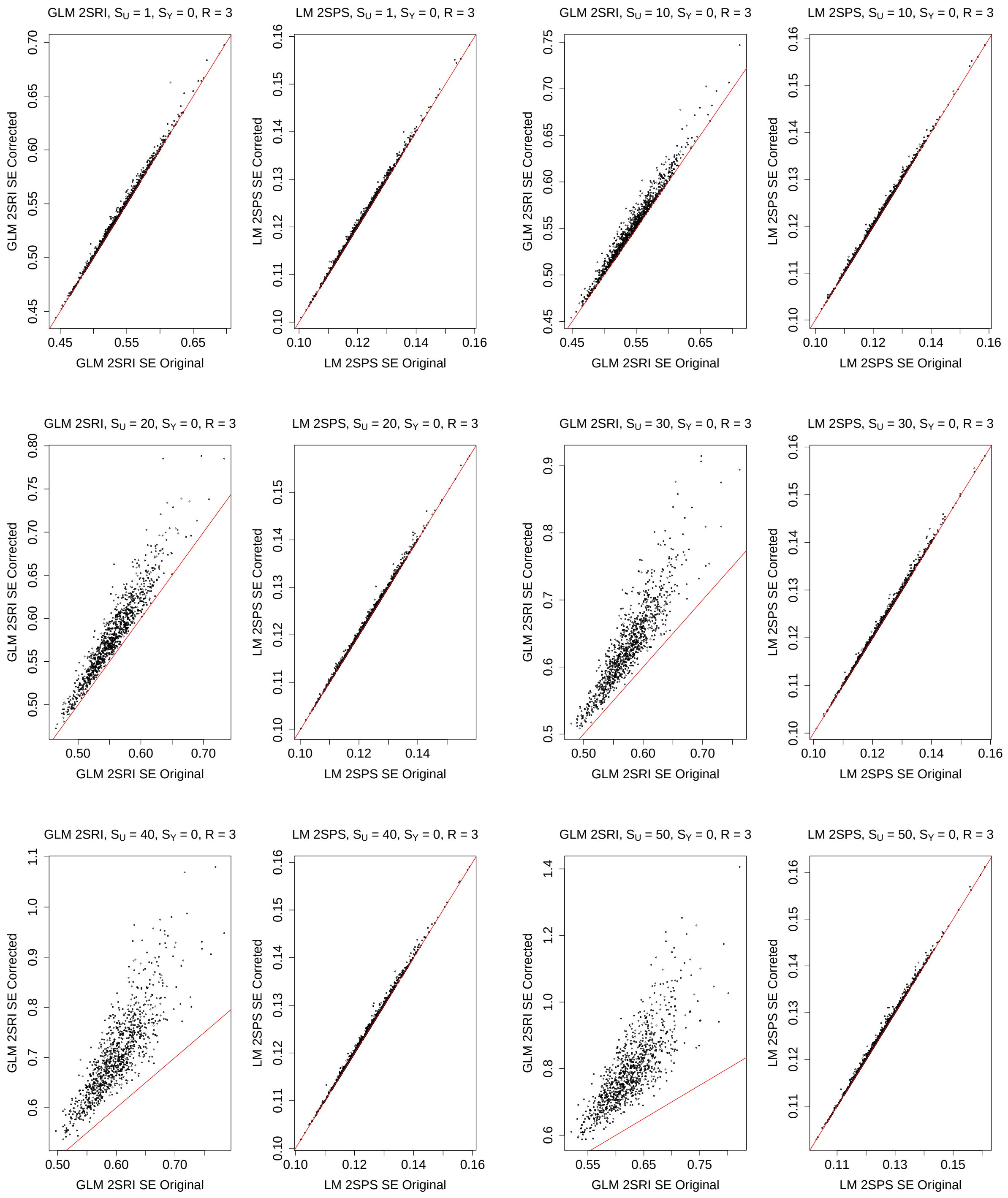

Figure S22: Simulations with the two-sample approaches: comparison of the original and corrected standard error estimates, $S_Y = 1, R = 3$.

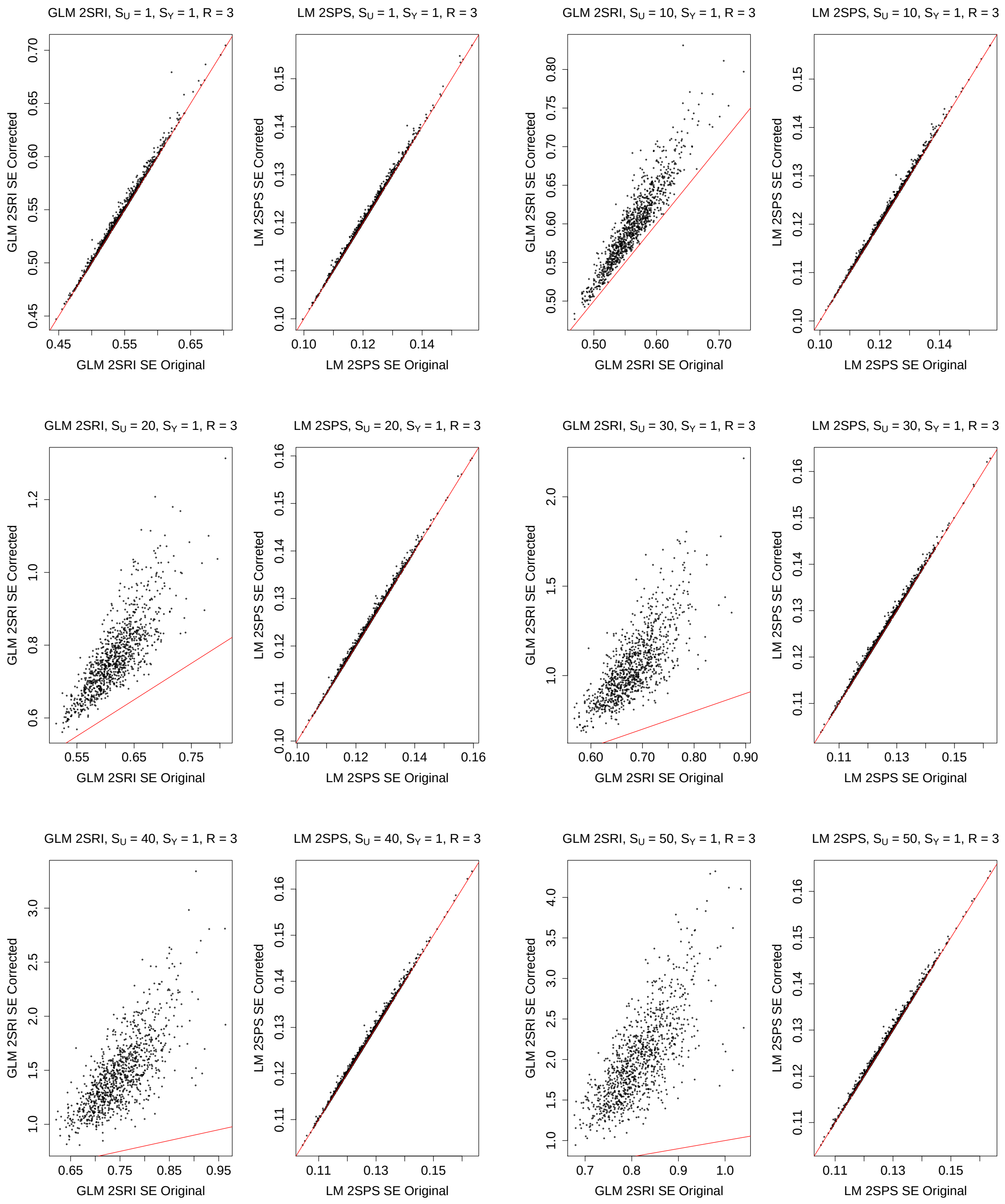

Figure S23: Simulations with the two-sample approaches: comparison of the original and corrected standard error estimates, $S_Y = 2, R = 3$.

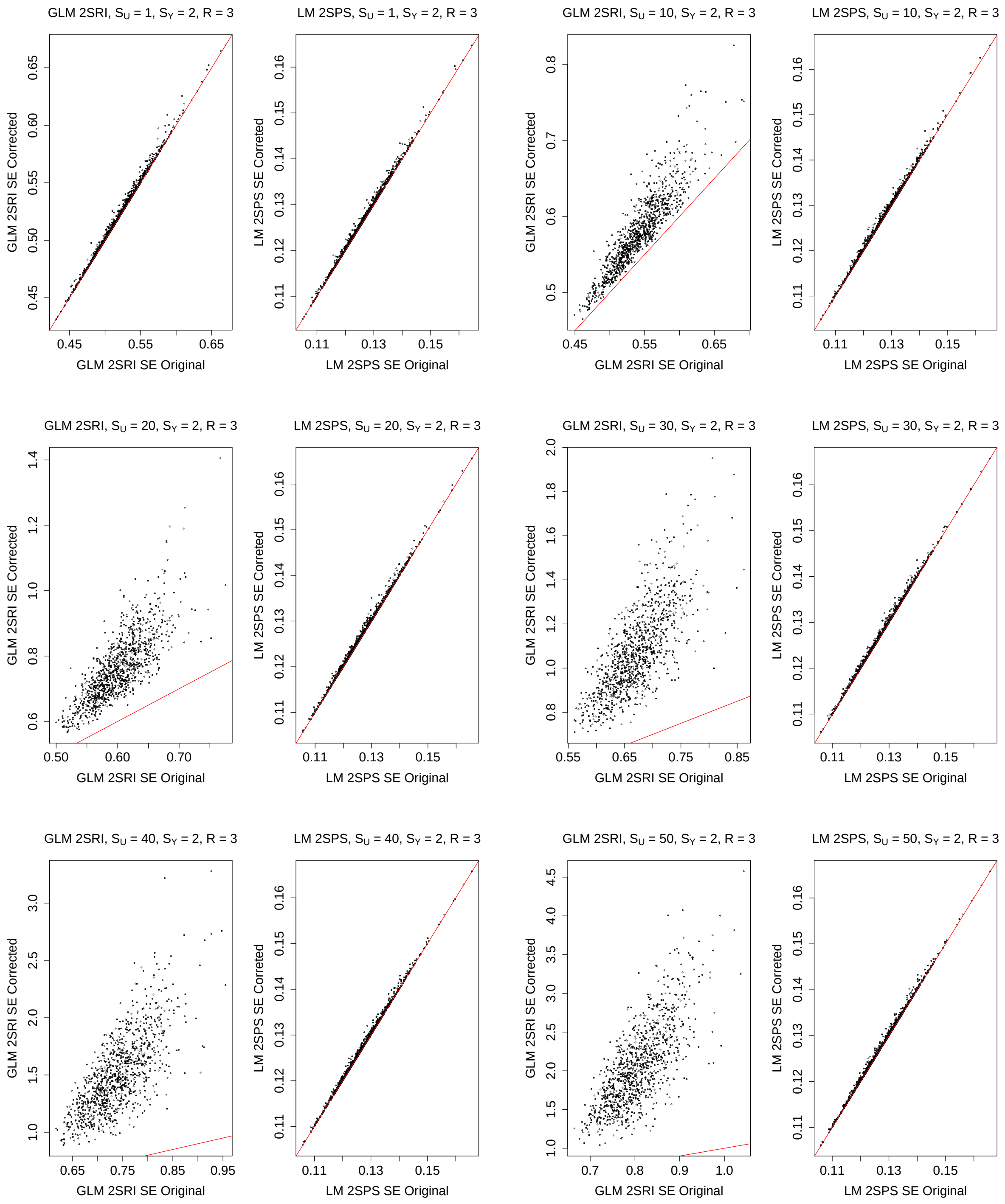

Figure S24: Simulations with the two-sample approaches: comparison of the original and corrected standard error estimates, $S_Y = 3, R = 3$.

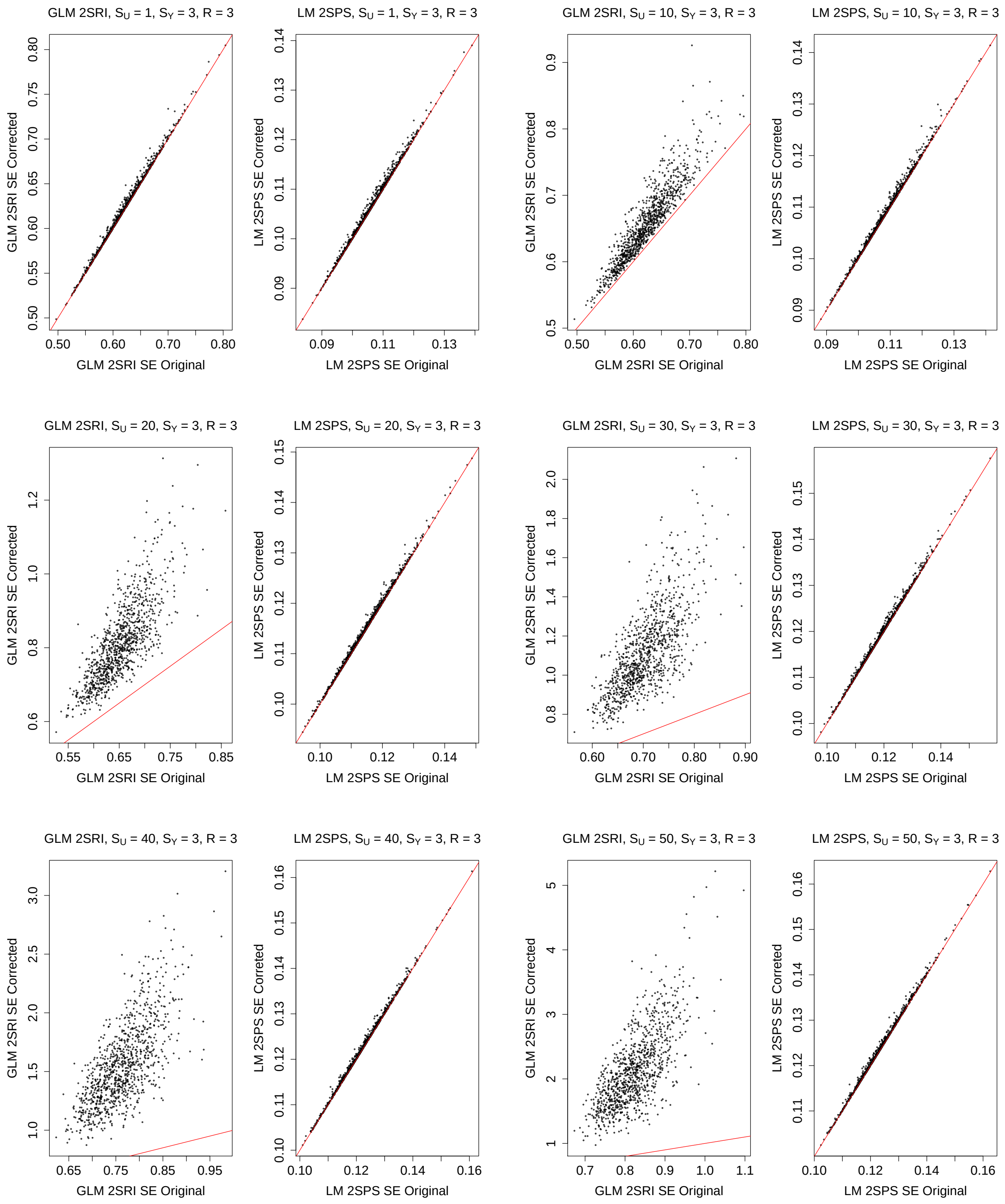

Figure S25: Simulations with the two-sample approaches: comparison of the original and corrected standard error estimates, $S_Y = 4, R = 3$.

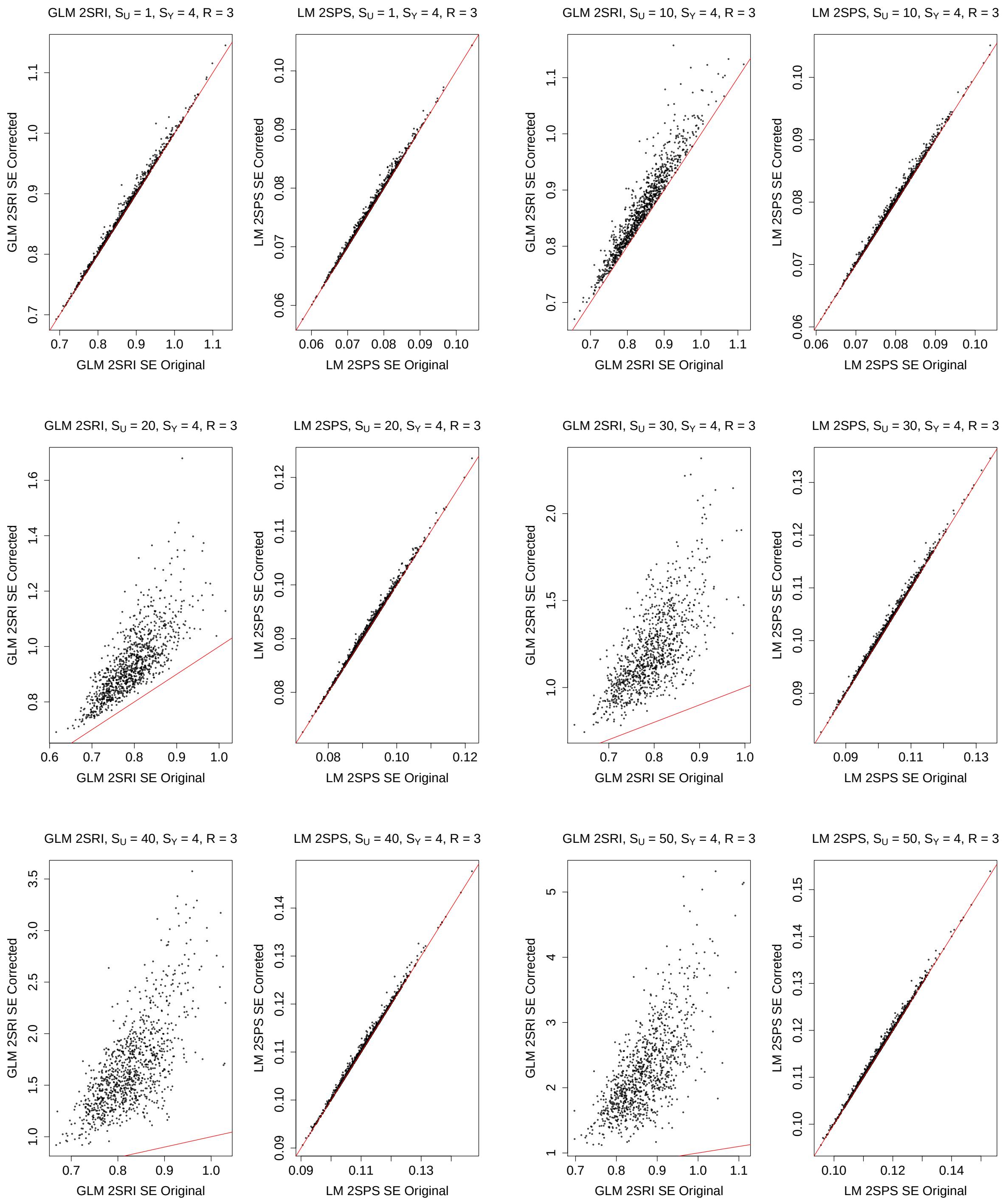

Figure S26: Simulations with the two-sample approaches: comparison of the original and corrected standard error estimates, $S_Y = 5, R = 3$.

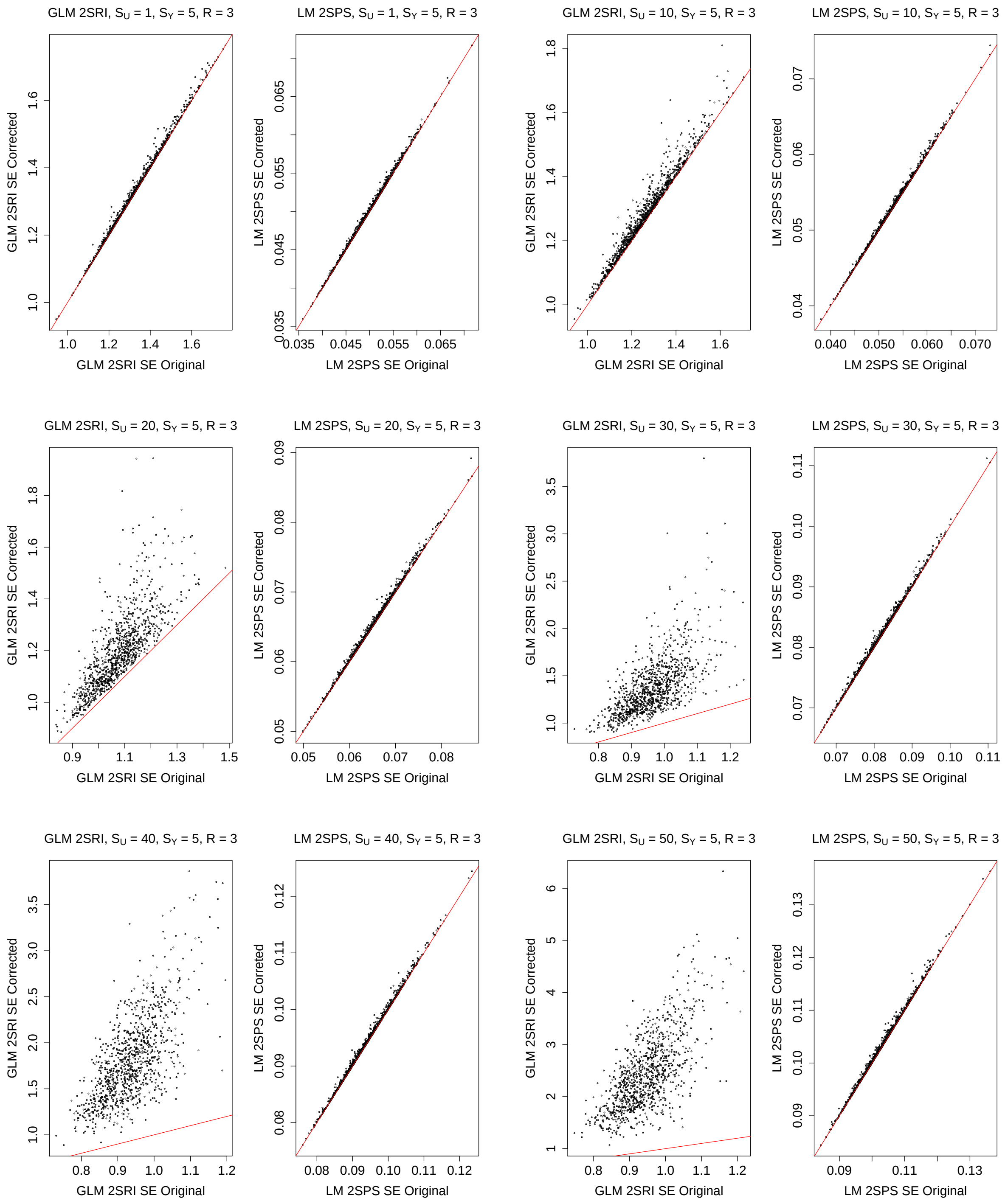

Figure S27: Simulations with the two-sample approaches: comparison of the original and corrected standard error estimates, $S_Y = 0, R = 5$.

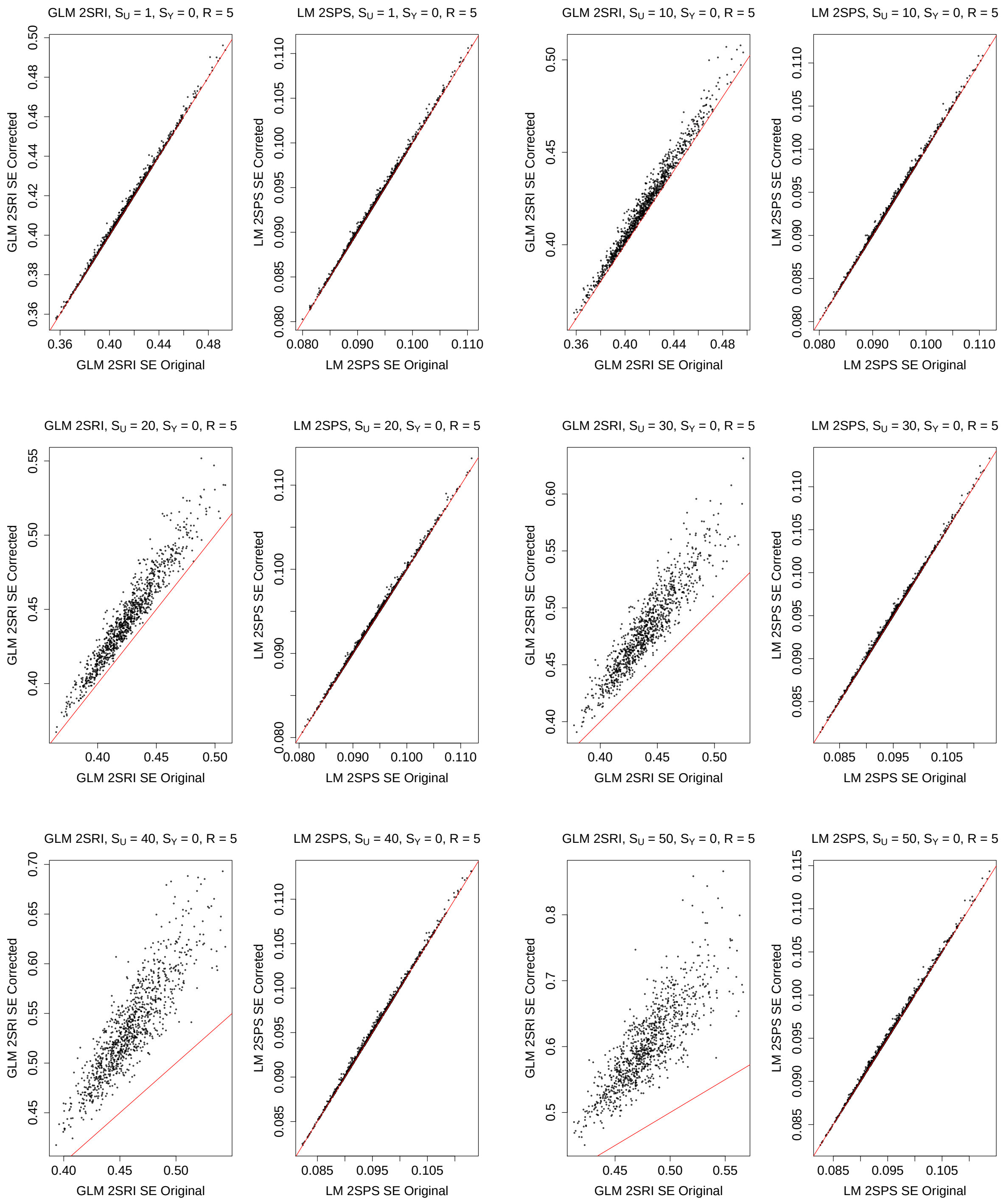

Figure S28: Simulations with the two-sample approaches: comparison of the original and corrected standard error estimates, $S_Y = 1, R = 5$.

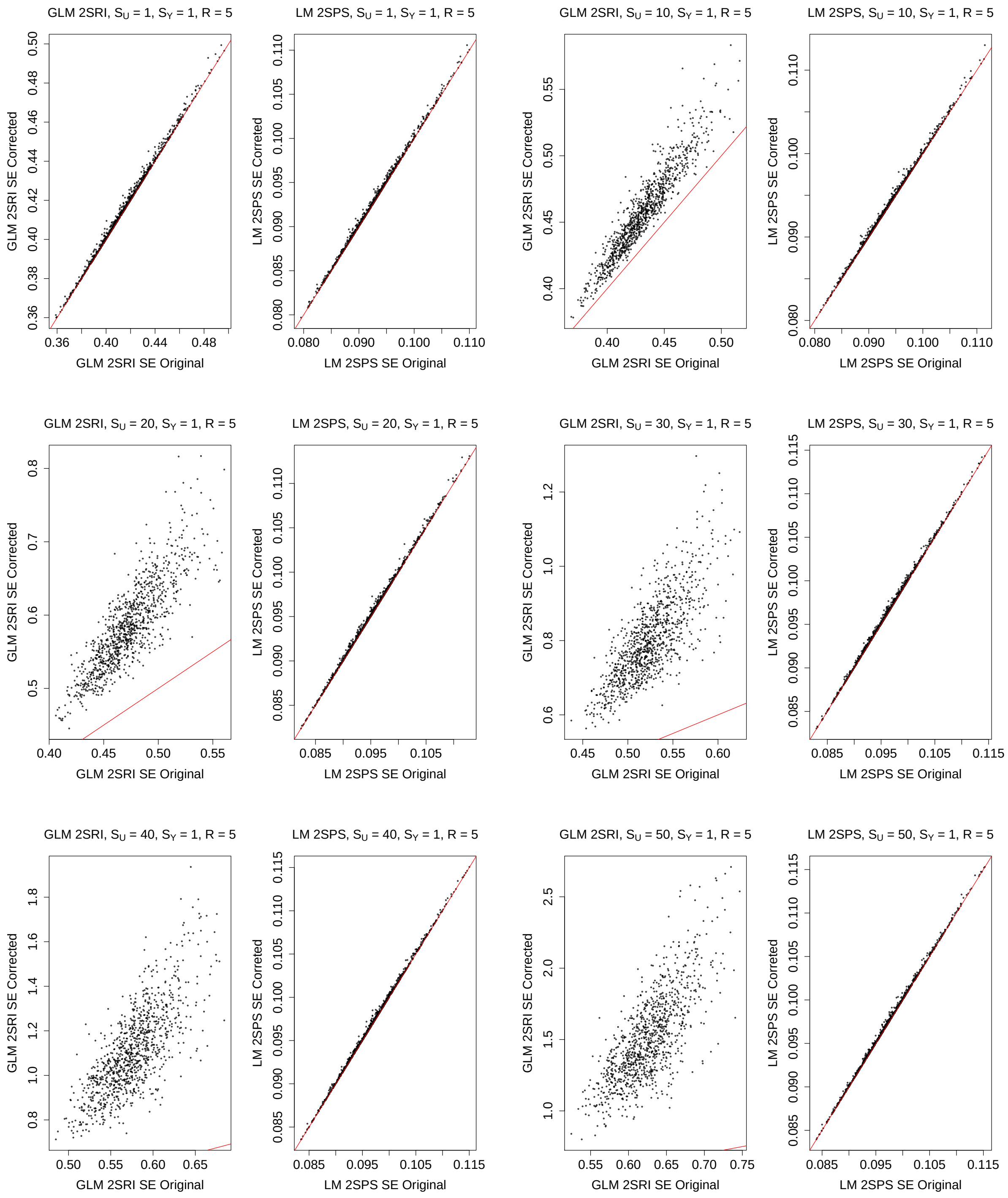

Figure S29: Simulations with the two-sample approaches: comparison of the original and corrected standard error estimates, $S_Y = 2, R = 5$.

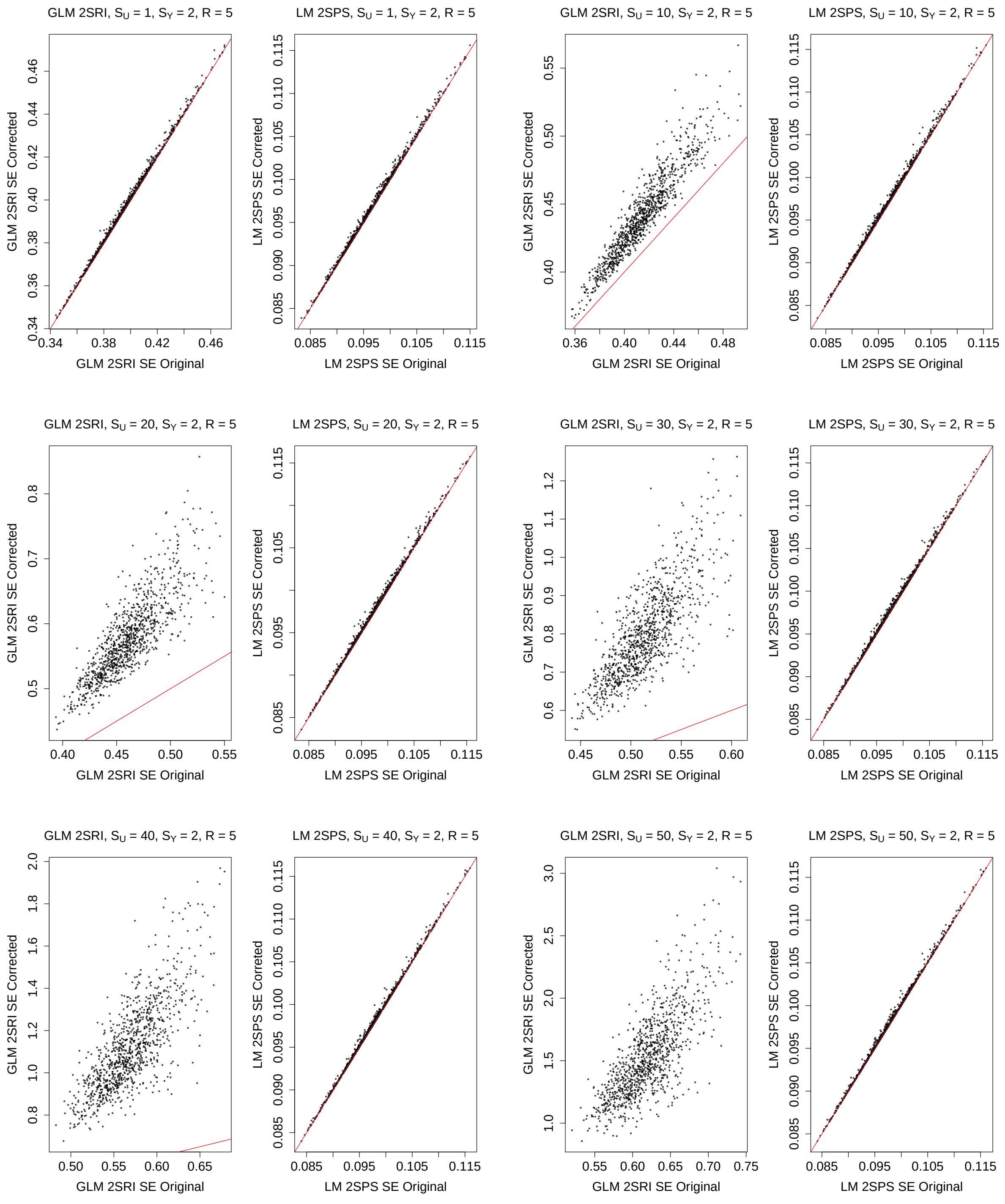

Figure S30: Simulations with the two-sample approaches: comparison of the original and corrected standard error estimates, $S_Y = 3, R = 5$.

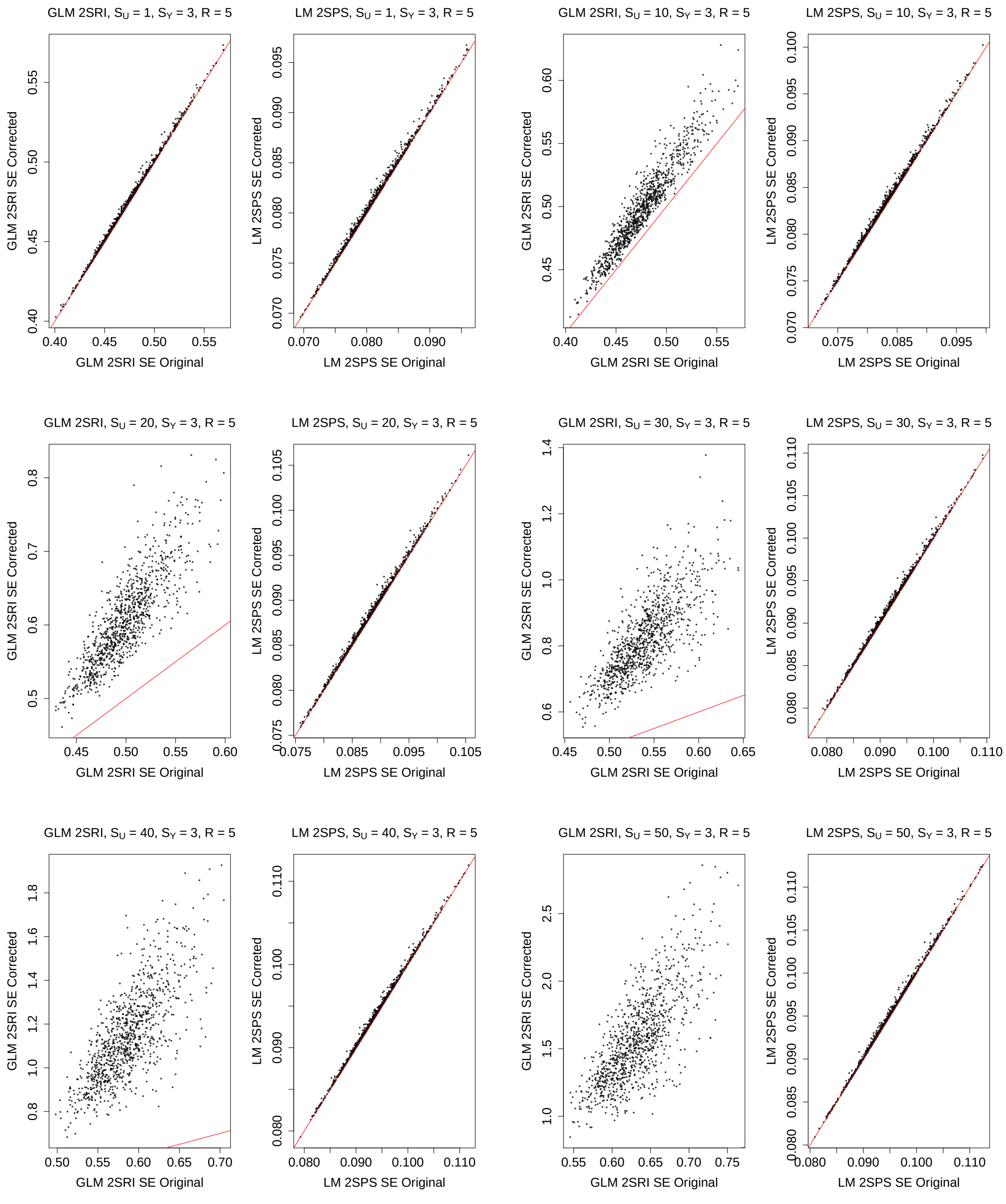

Figure S31: Simulations with the two-sample approaches: comparison of the original and corrected standard error estimates, $S_Y = 4, R = 5$.

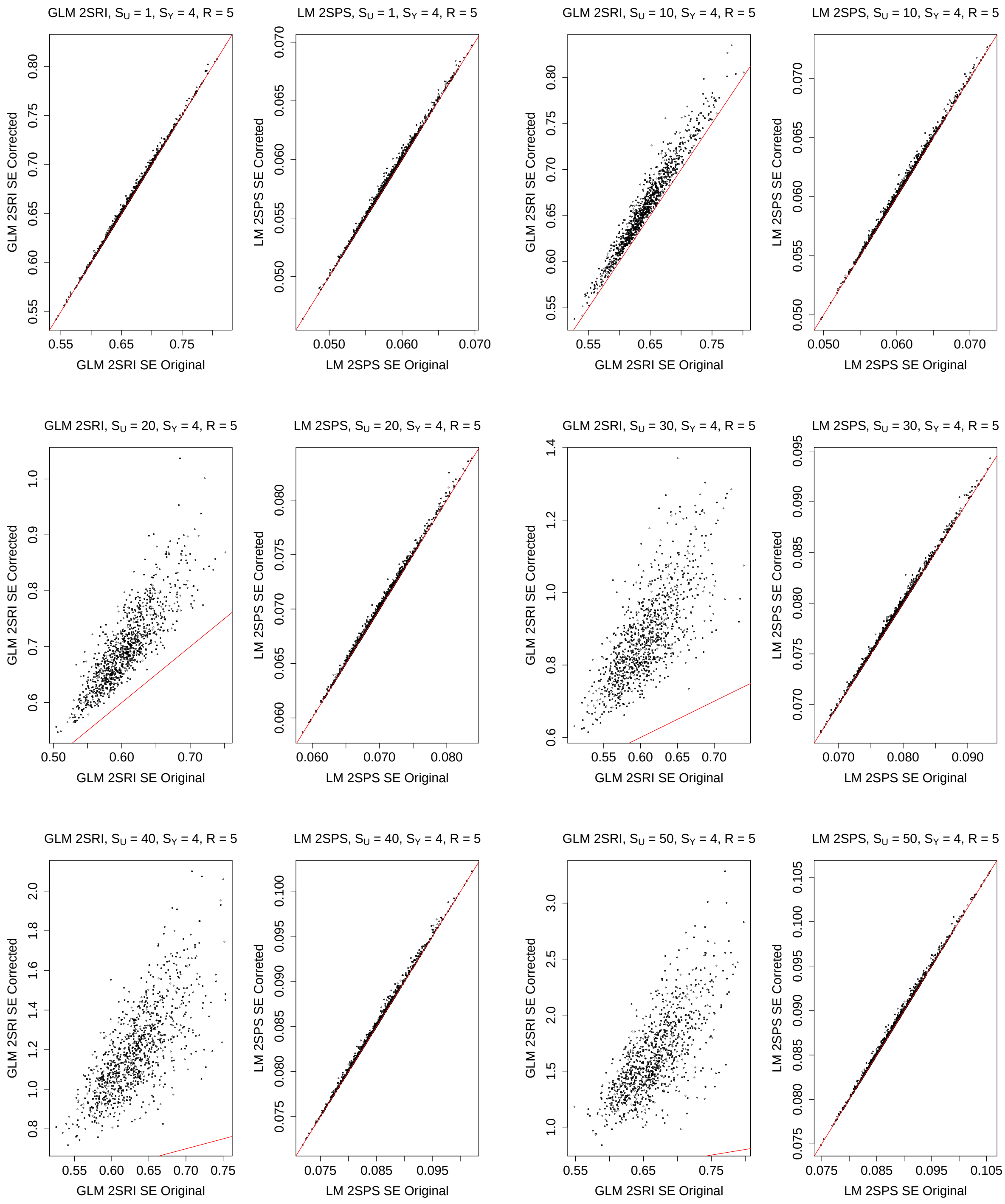

Figure S32: Simulations with the two-sample approaches: comparison of the original and corrected standard error estimates, $S_Y = 5, R = 5$.

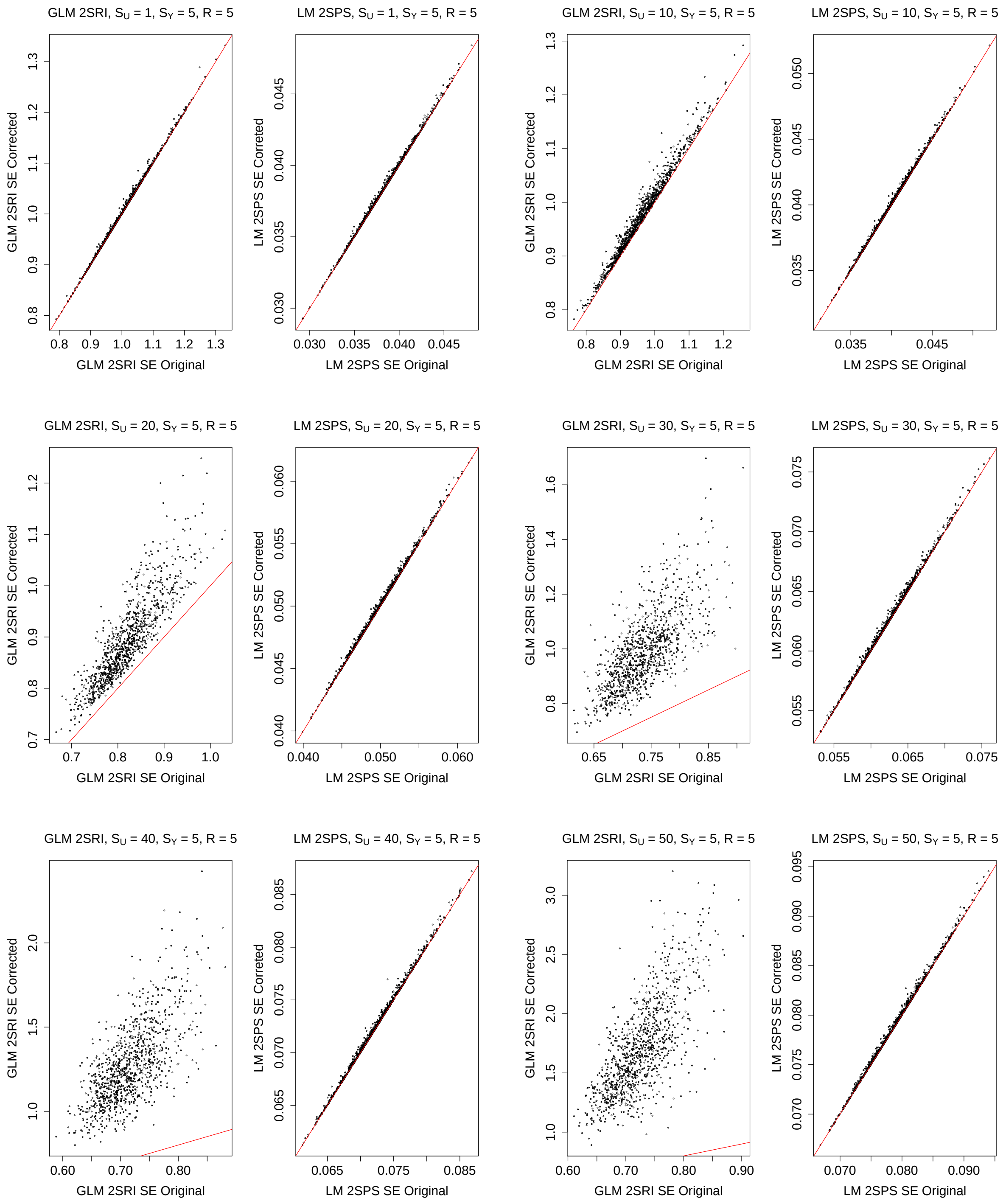

Figure S33: Simulations with the two-sample approaches: comparison of the original and corrected standard error estimates, $S_Y = 0, R = 7$.

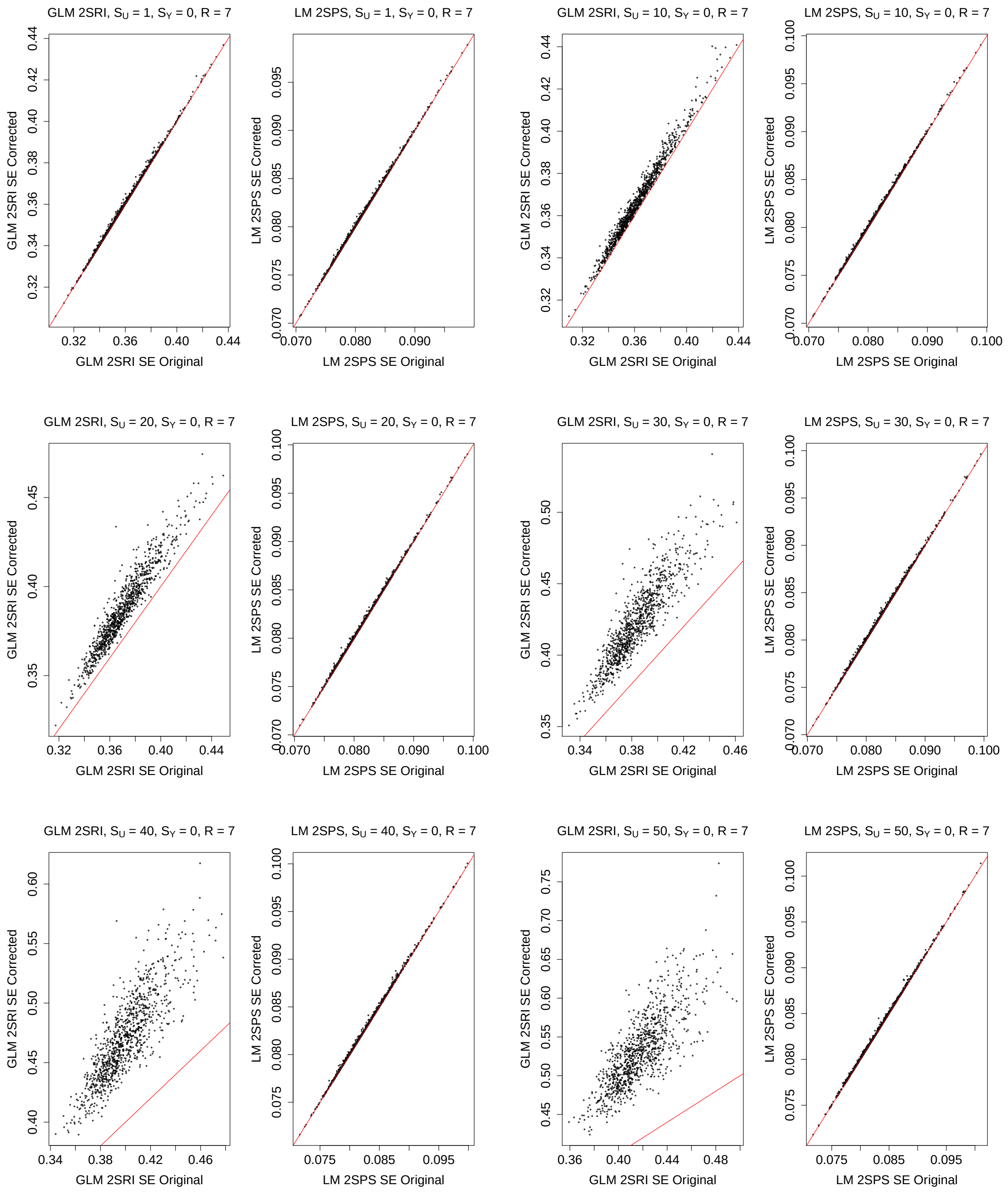

Figure S34: Simulations with the two-sample approaches: comparison of the original and corrected standard error estimates, $S_Y = 1, R = 7$.

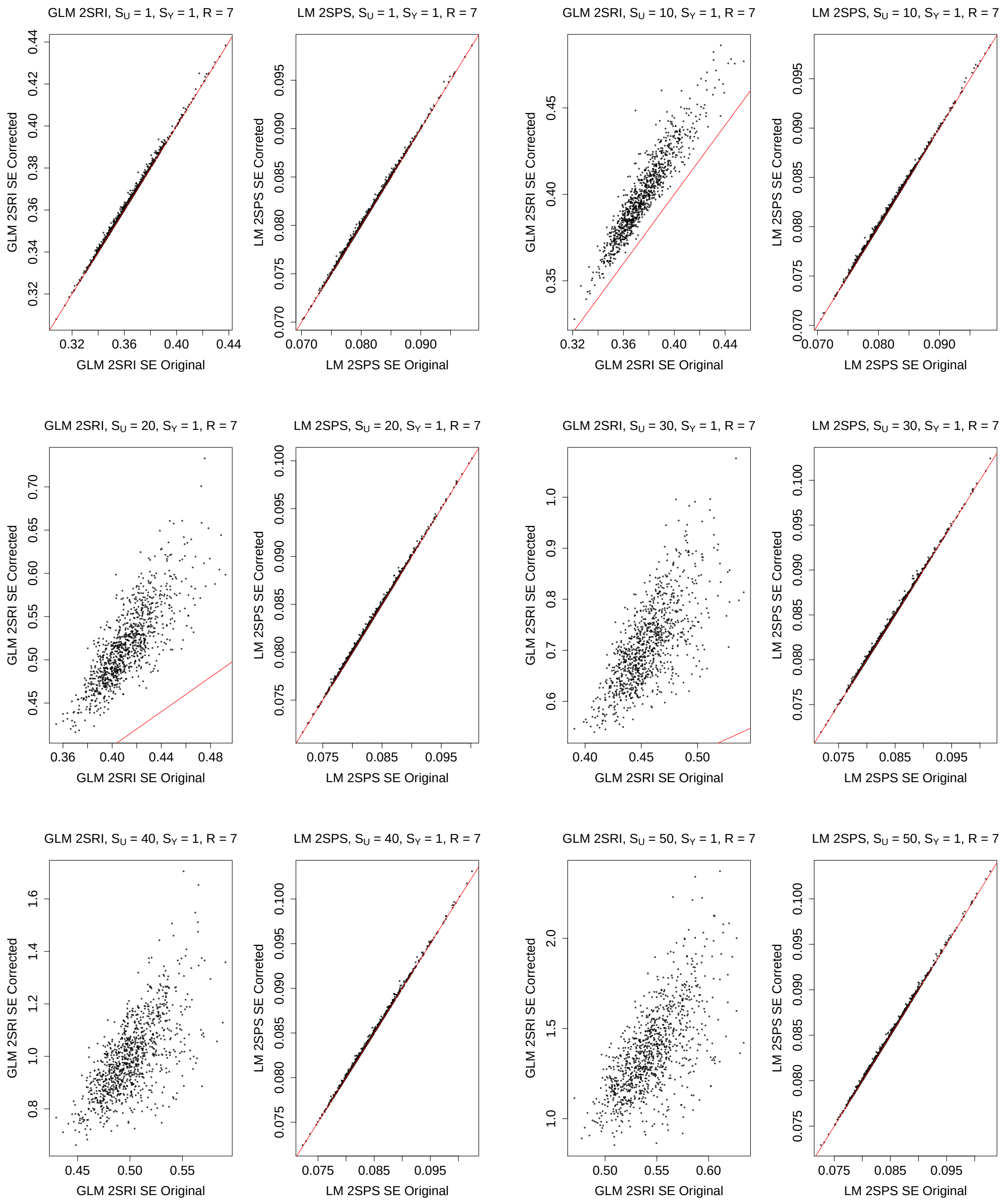

Figure S35: Simulations with the two-sample approaches: comparison of the original and corrected standard error estimates, $S_Y = 2, R = 7$.

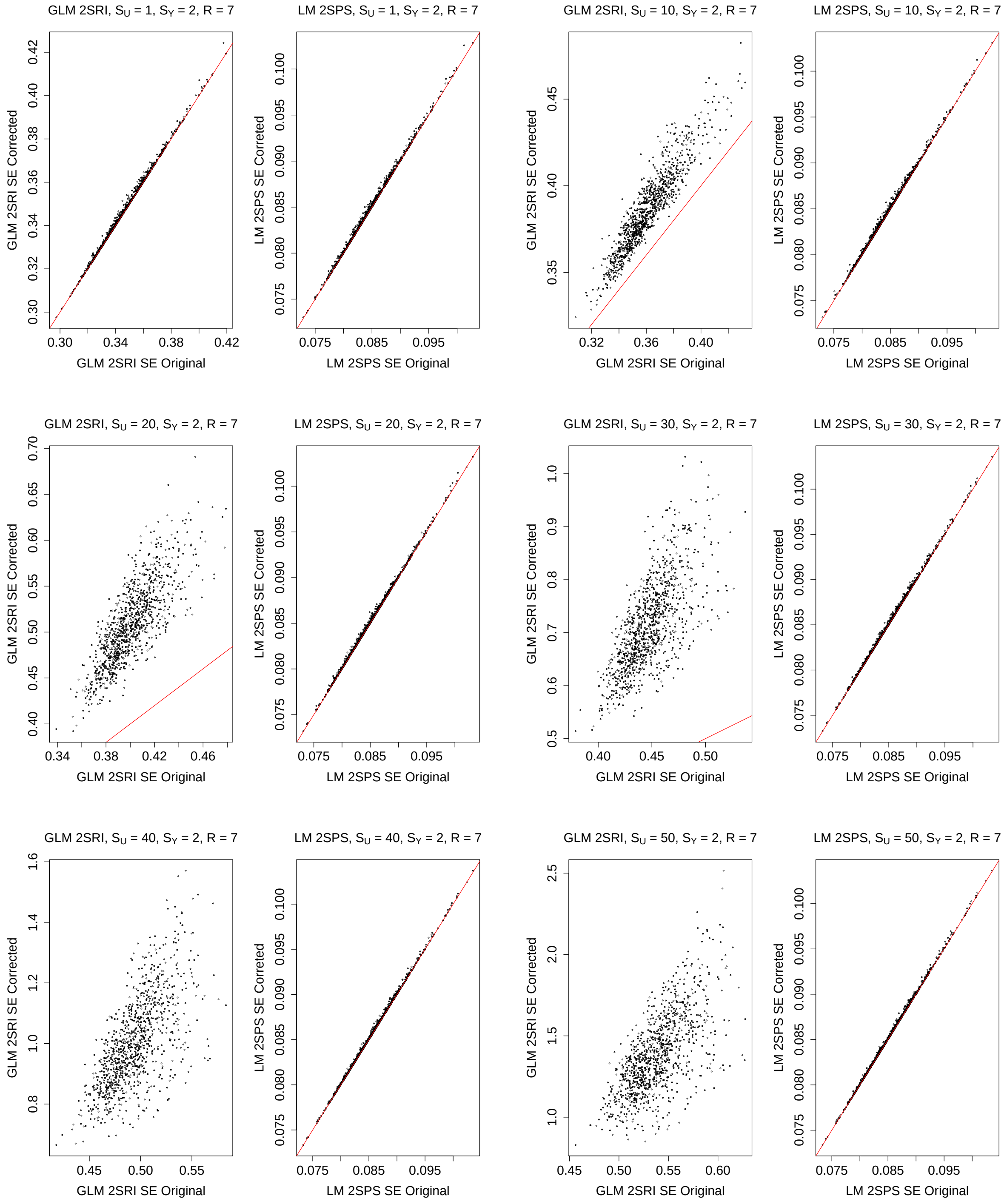

Figure S36: Simulations with the two-sample approaches: comparison of the original and corrected standard error estimates, $S_Y = 3, R = 7$.

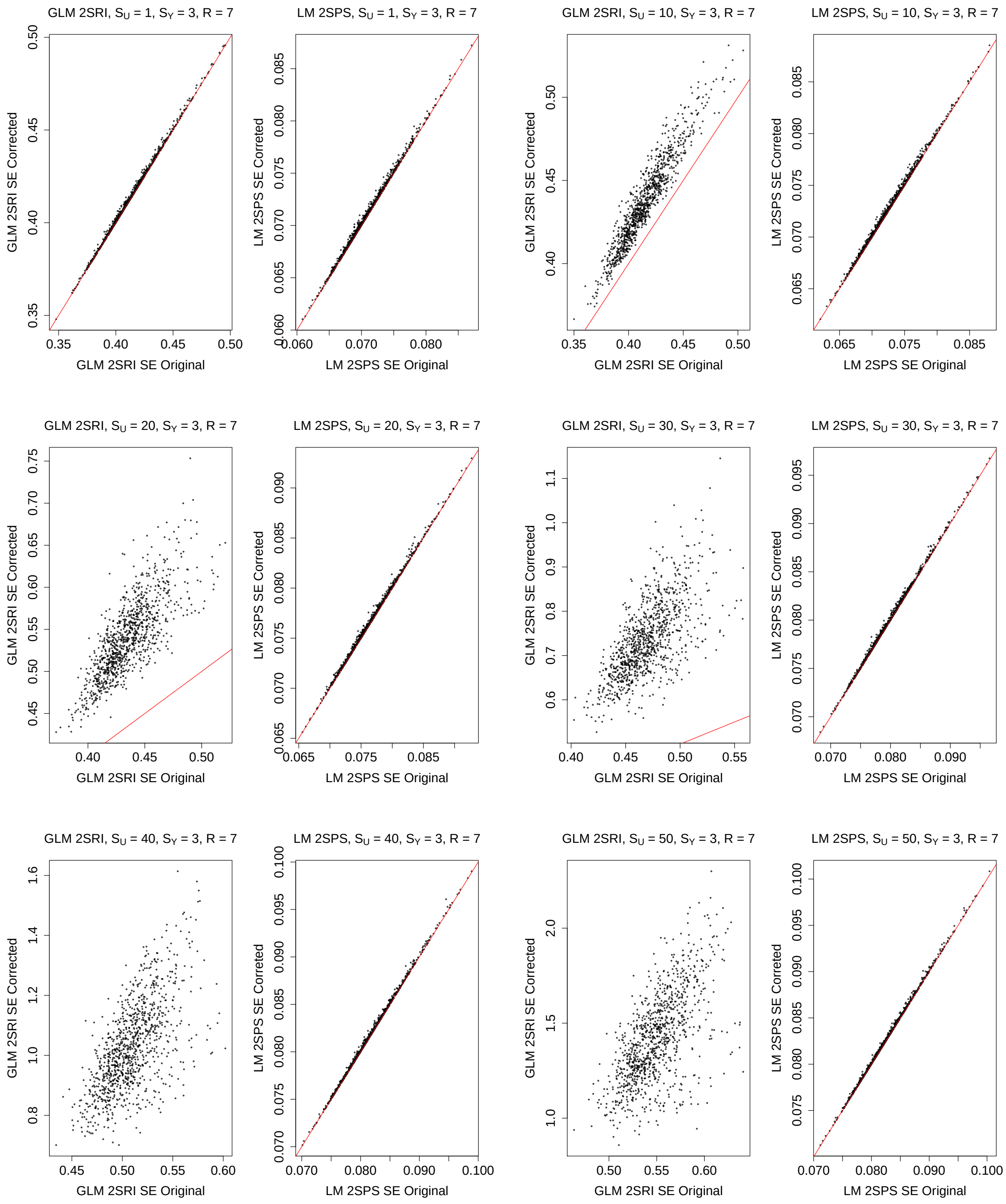

Figure S37: Simulations with the two-sample approaches: comparison of the original and corrected standard error estimates, $S_Y = 4, R = 7$.

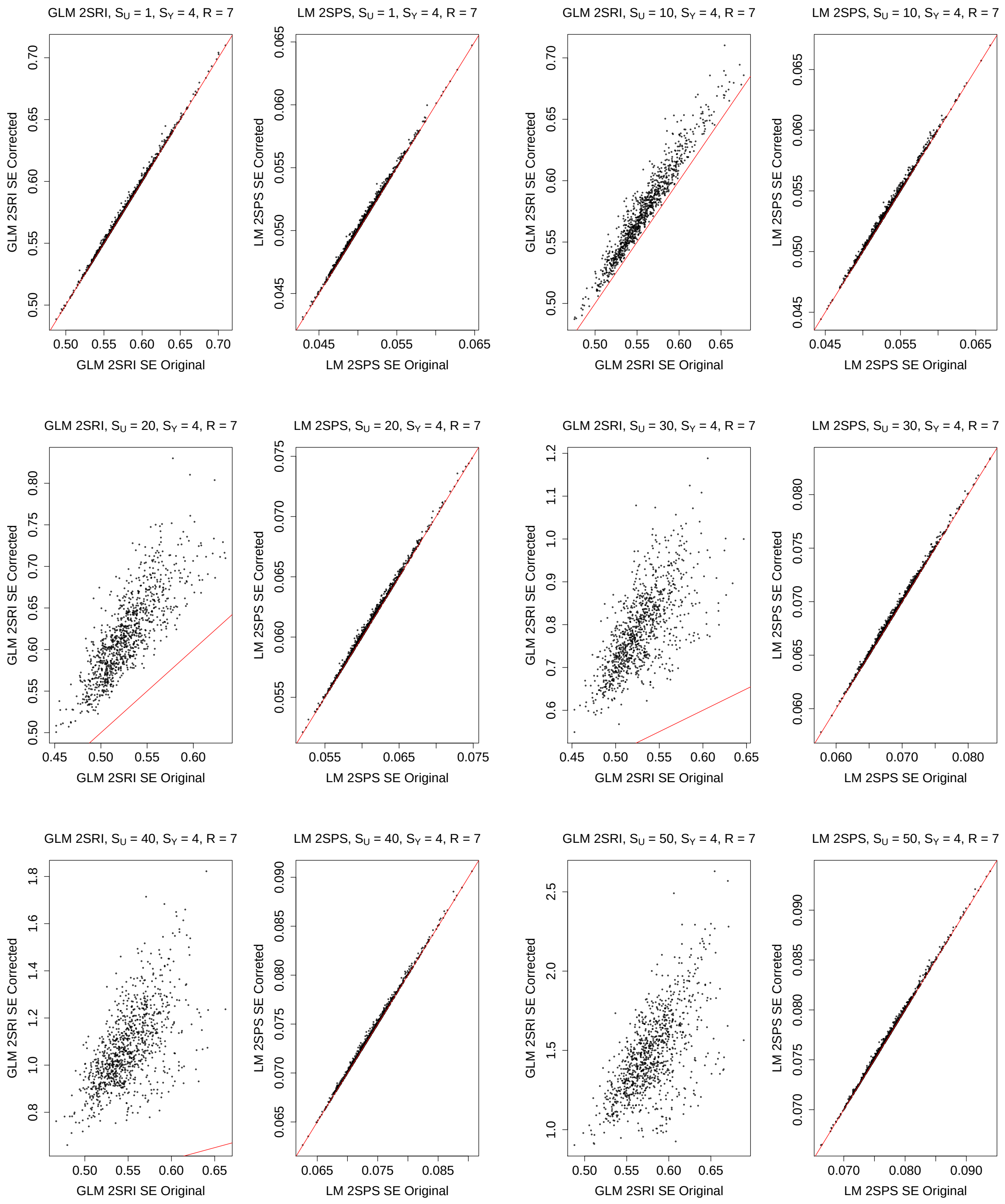

Figure S38: Simulations with the two-sample approaches: comparison of the original and corrected standard error estimates, $S_Y = 5, R = 7$.

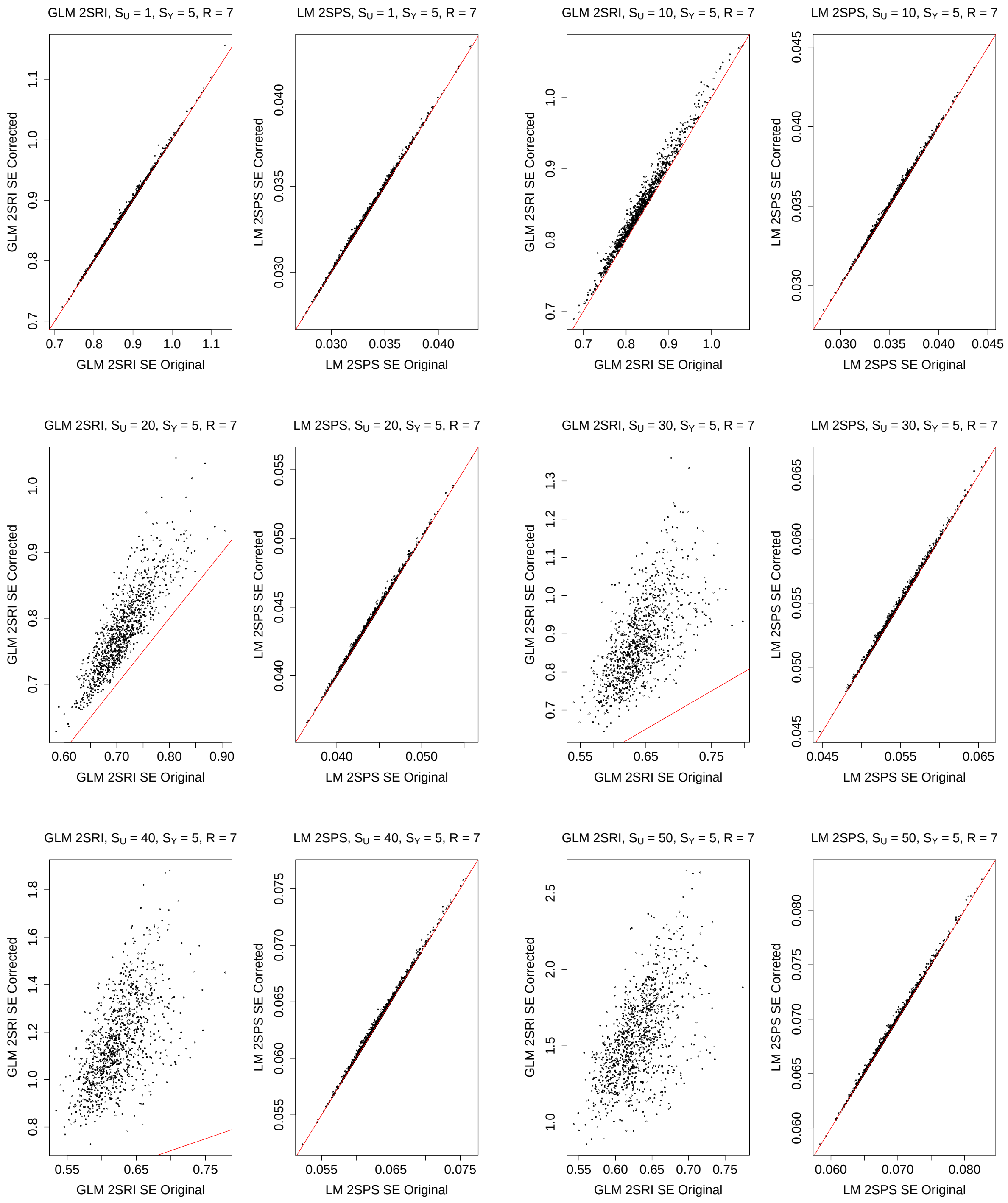

Figure S39: Simulations with the two-sample approaches: comparison of the original and corrected standard error estimates, $S_Y = 0, R = 9$.

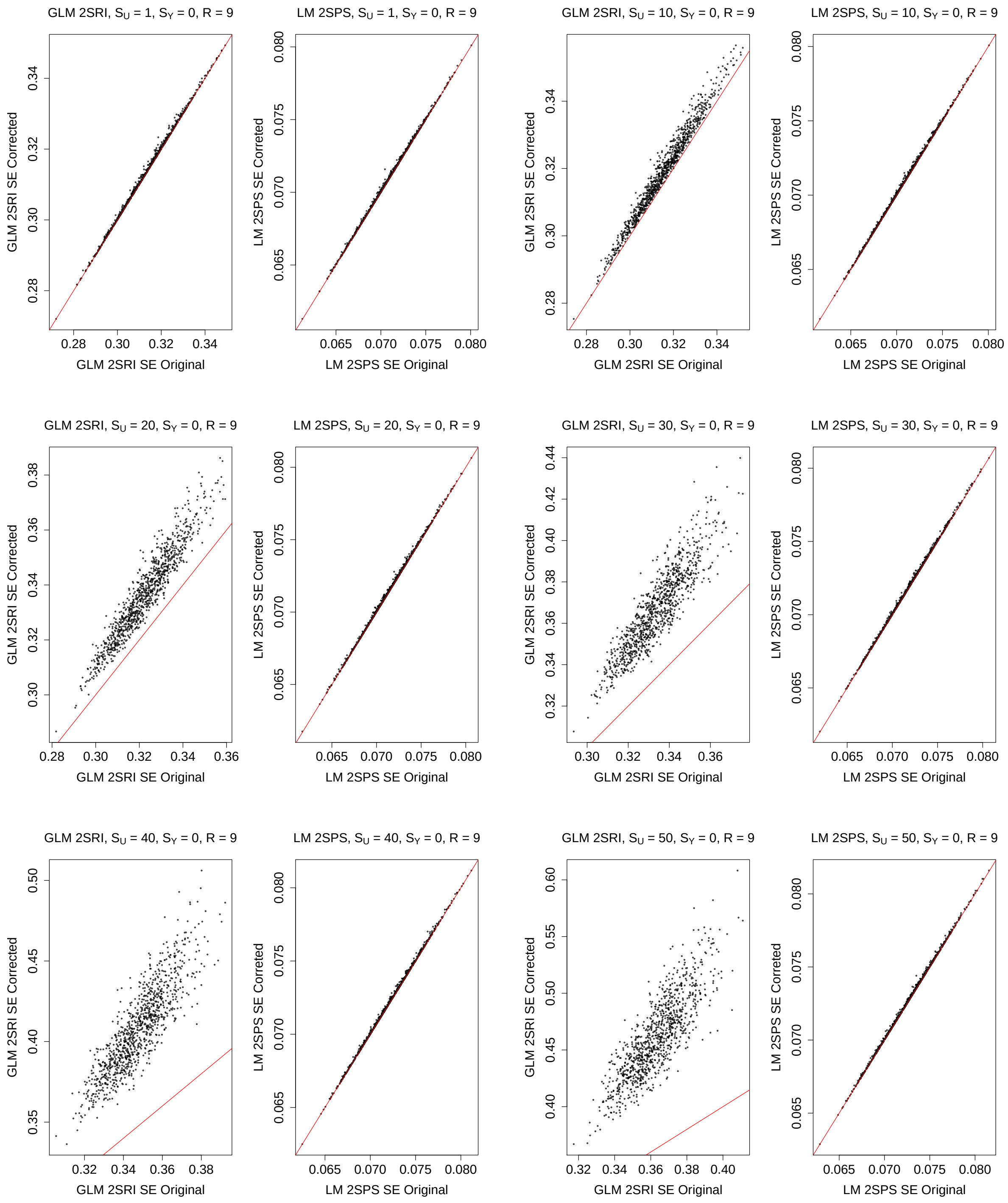

Figure S40: Simulations with the two-sample approaches: comparison of the original and corrected standard error estimates, $S_Y = 1, R = 9$.

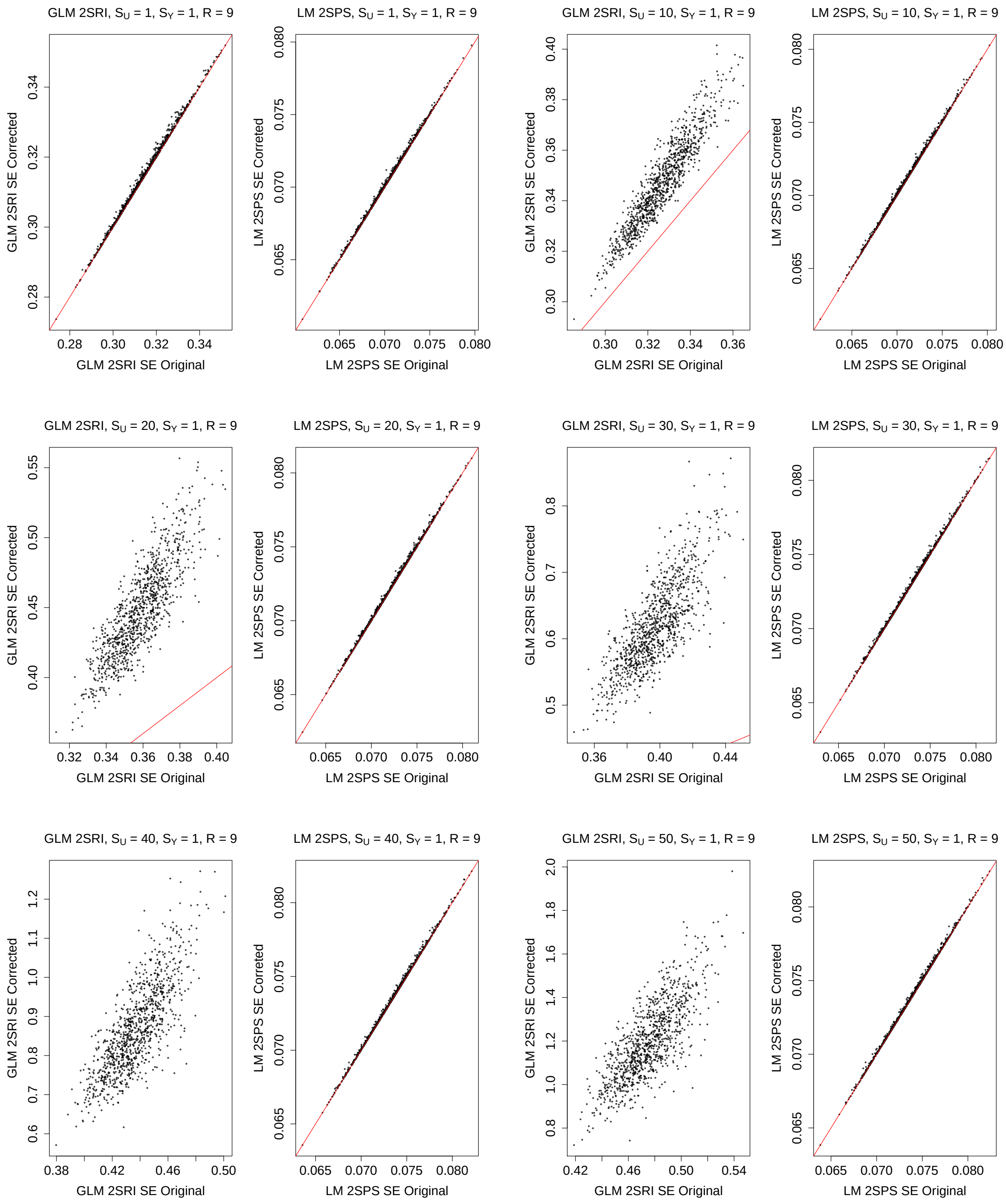

Figure S41: Simulations with the two-sample approaches: comparison of the original and corrected standard error estimates, $S_Y = 2, R = 9$.

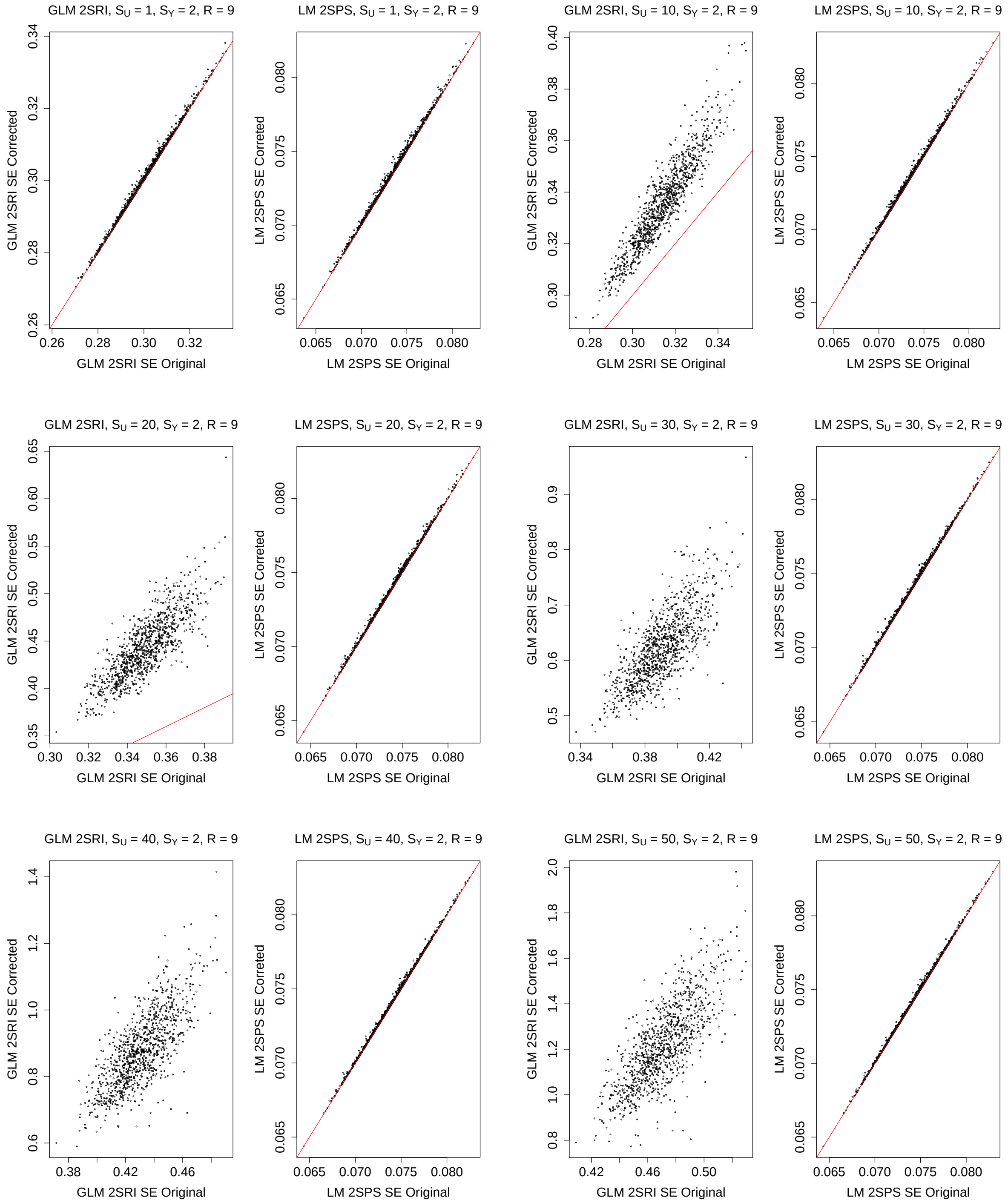

Figure S42: Simulations with the two-sample approaches: comparison of the original and corrected standard error estimates, $S_Y = 3, R = 9$.

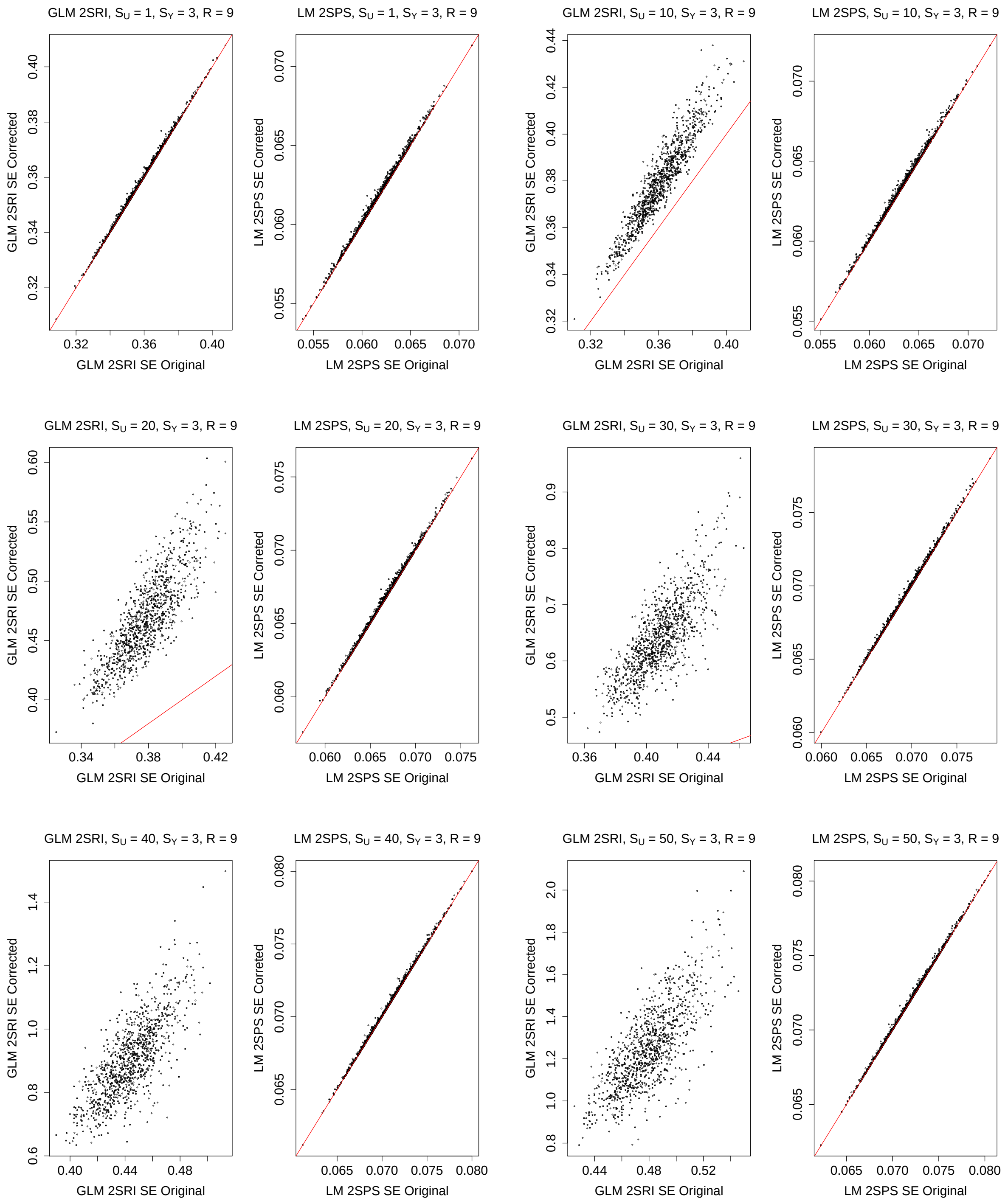

Figure S43: Simulations with the two-sample approaches: comparison of the original and corrected standard error estimates, $S_Y = 4, R = 9$.

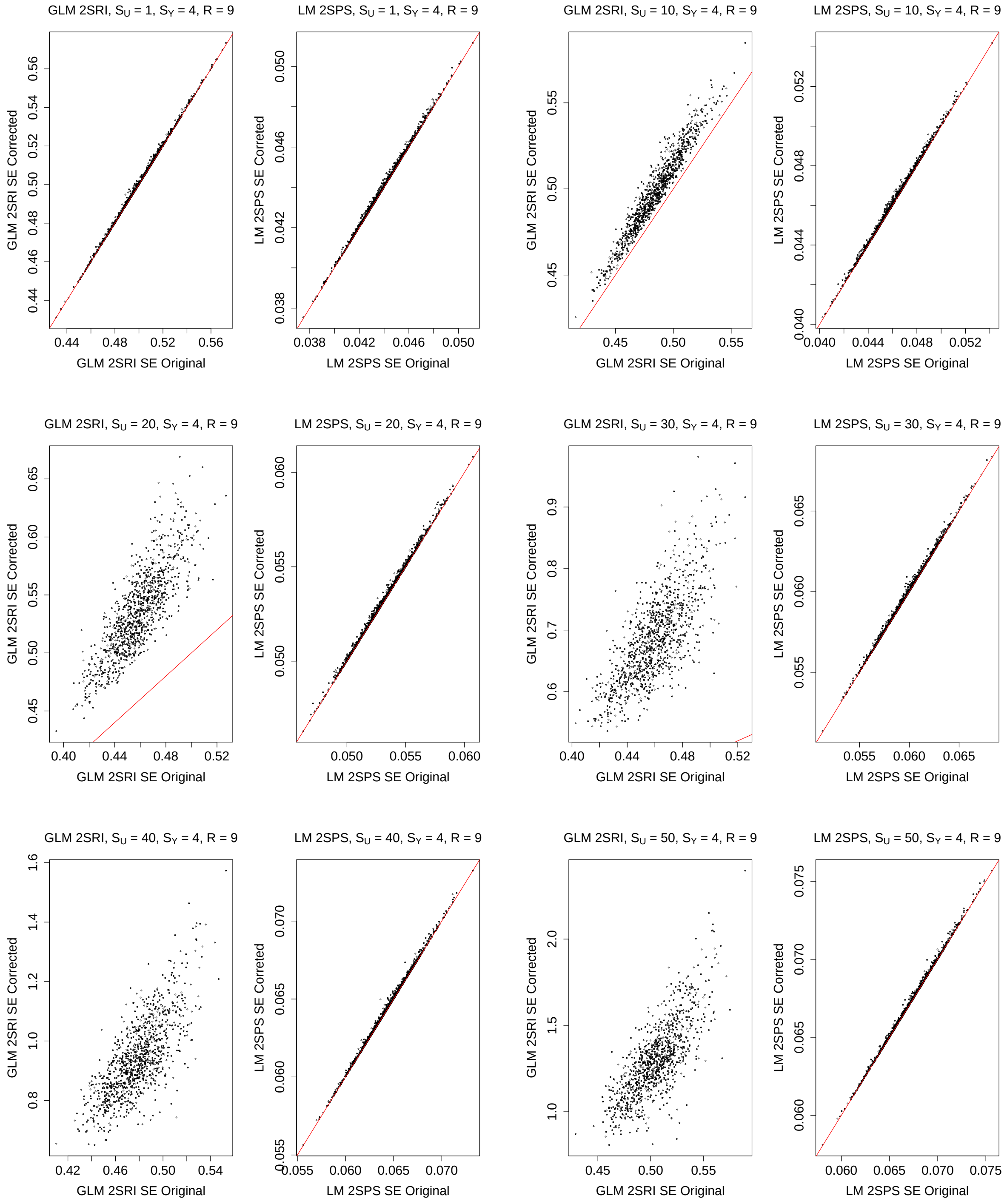

Figure S44: Simulations with the two-sample approaches: comparison of the original and corrected standard error estimates, $S_Y = 5, R = 9$.

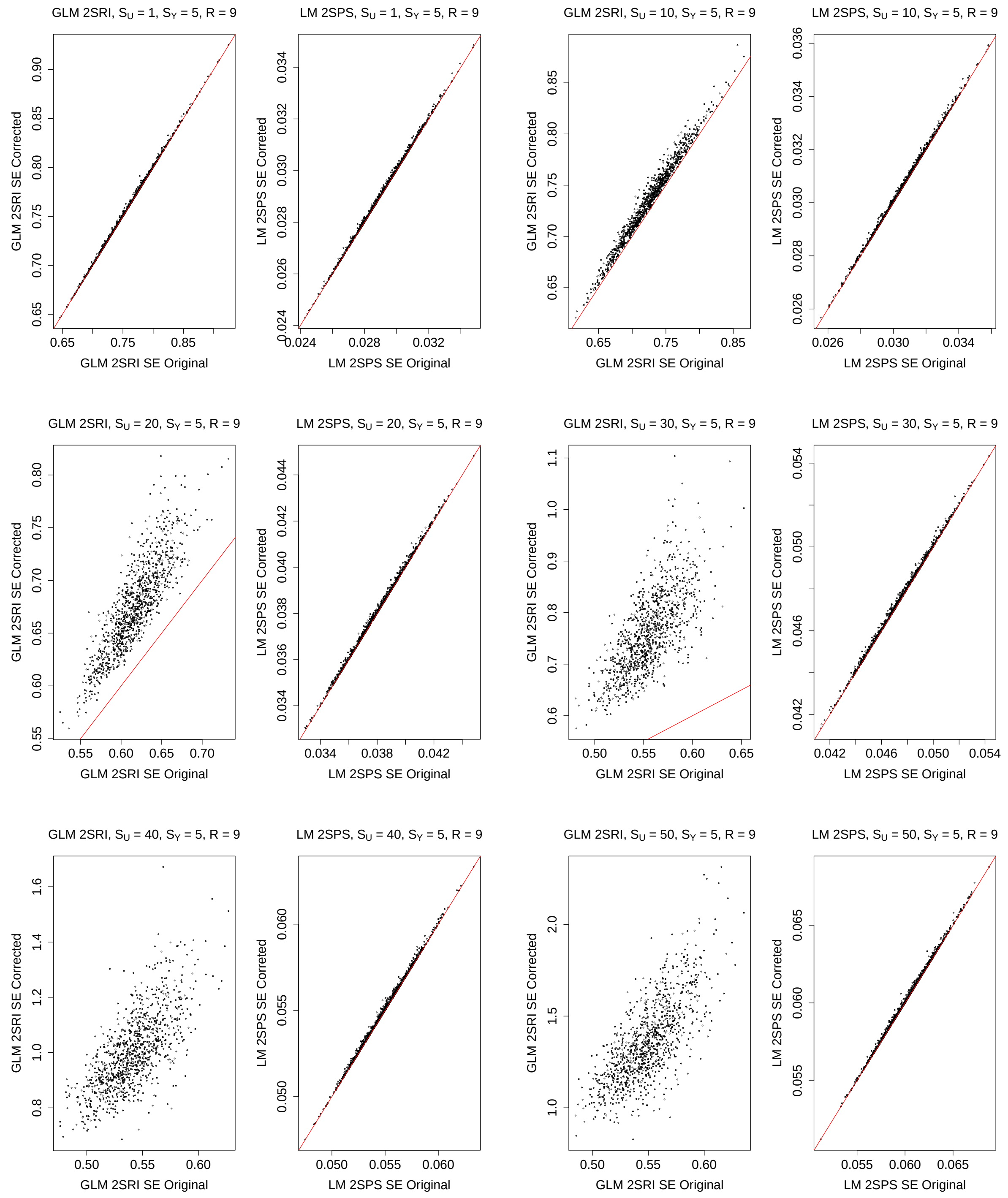

#### 2.2 One-sample Approaches

##### 2.2.1 Type I Error

Figure S45: Simulations with the one-sample approaches: empirical Type I error rates of various methods.

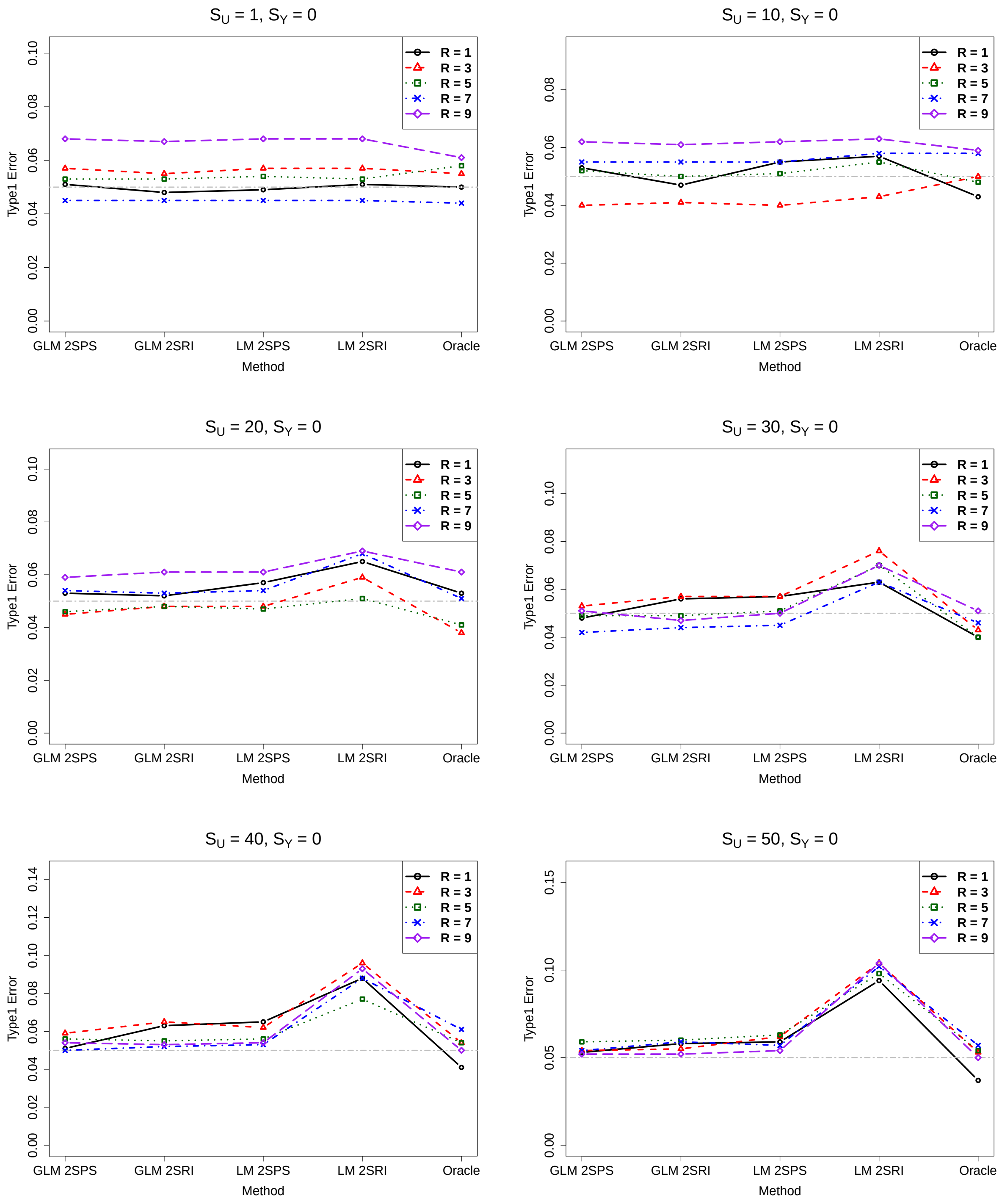

##### 2.2.2 Power

Figure S46: Simulations with the one-sample approaches: empirical power of various methods, $S_Y = 1$.

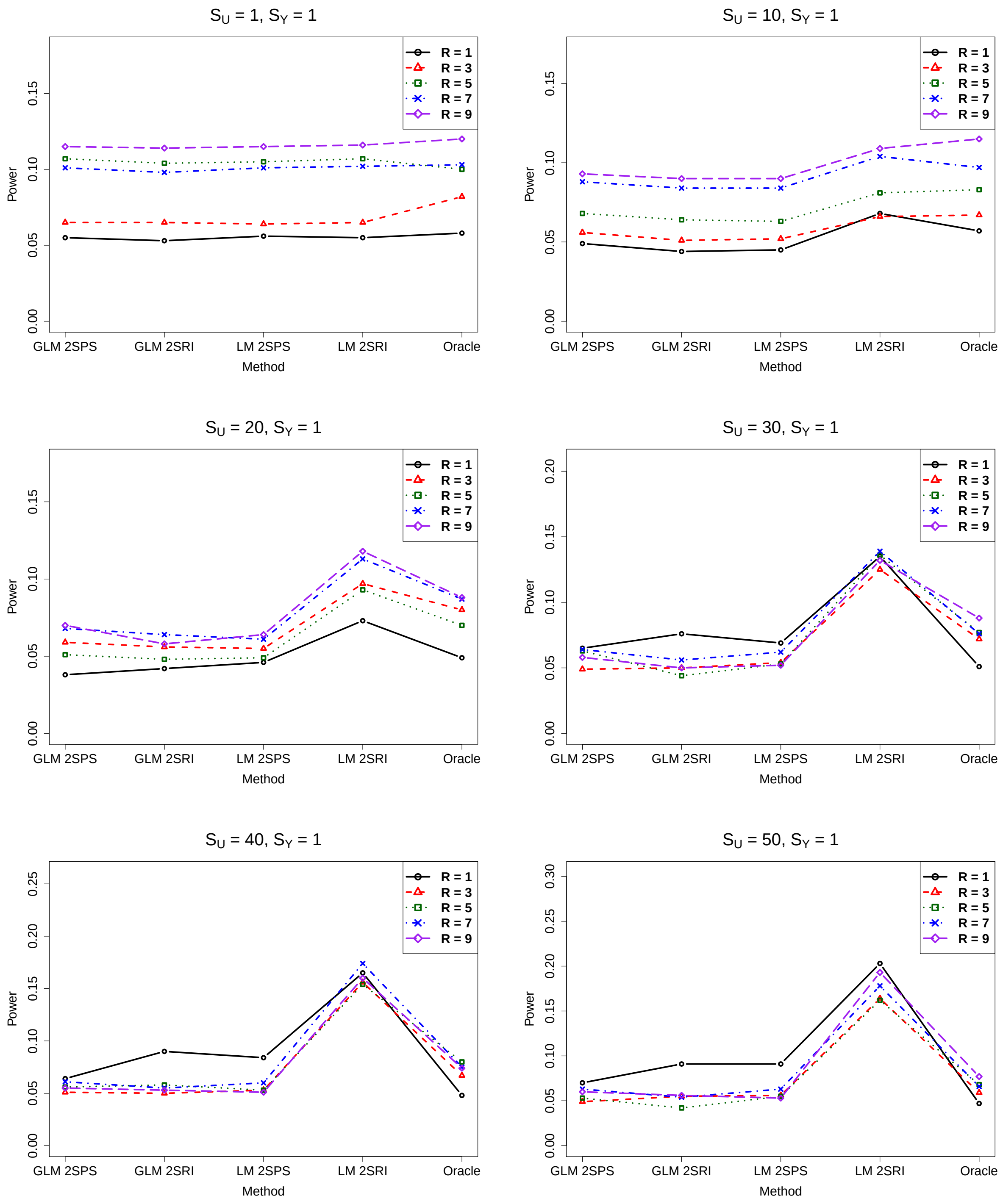

Figure S47: Simulations with the one-sample approaches: empirical power of various methods, $S_Y = 2$.

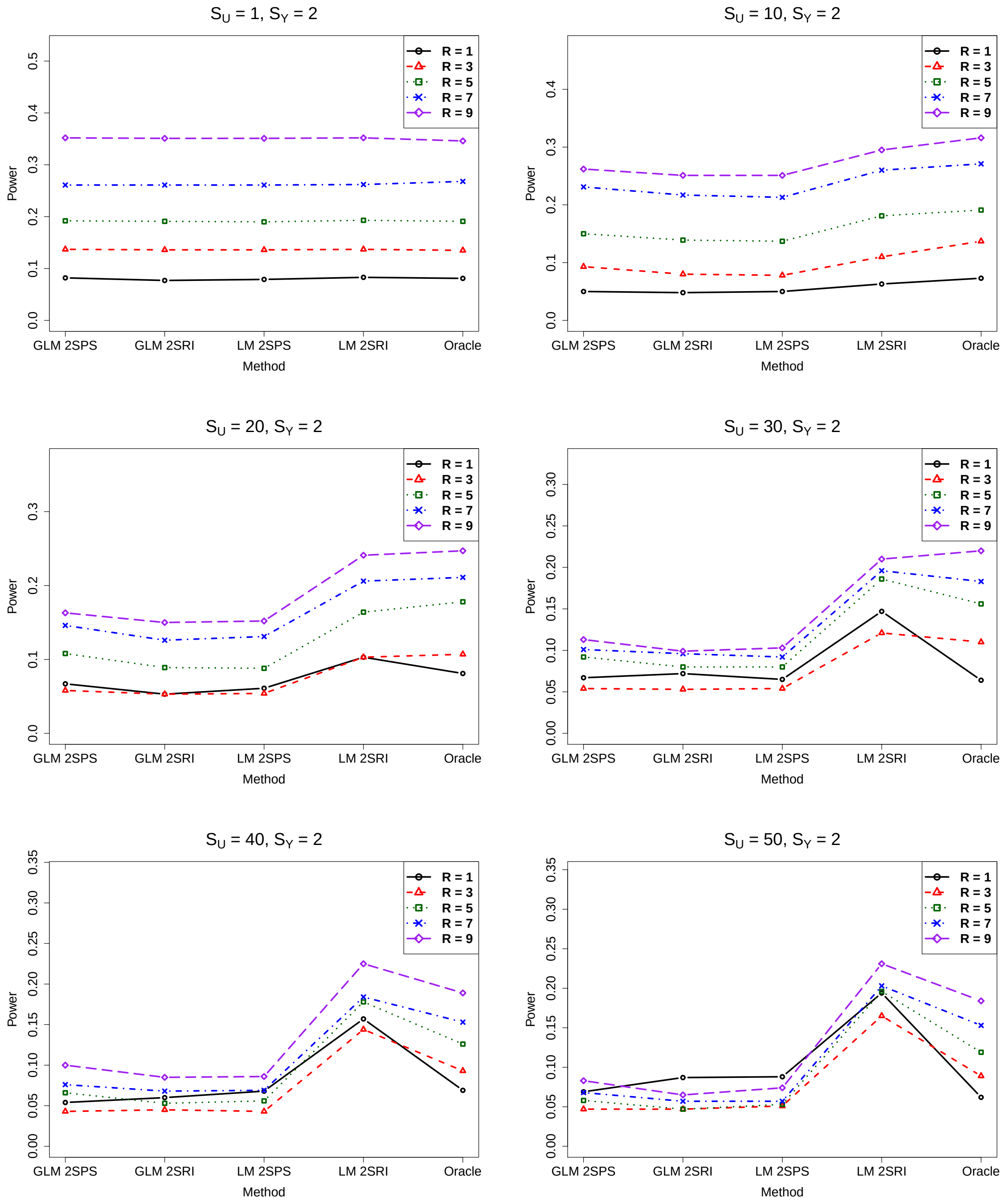

Figure S48: Simulations with the one-sample approaches: empirical power of various methods, $S_Y = 3$.

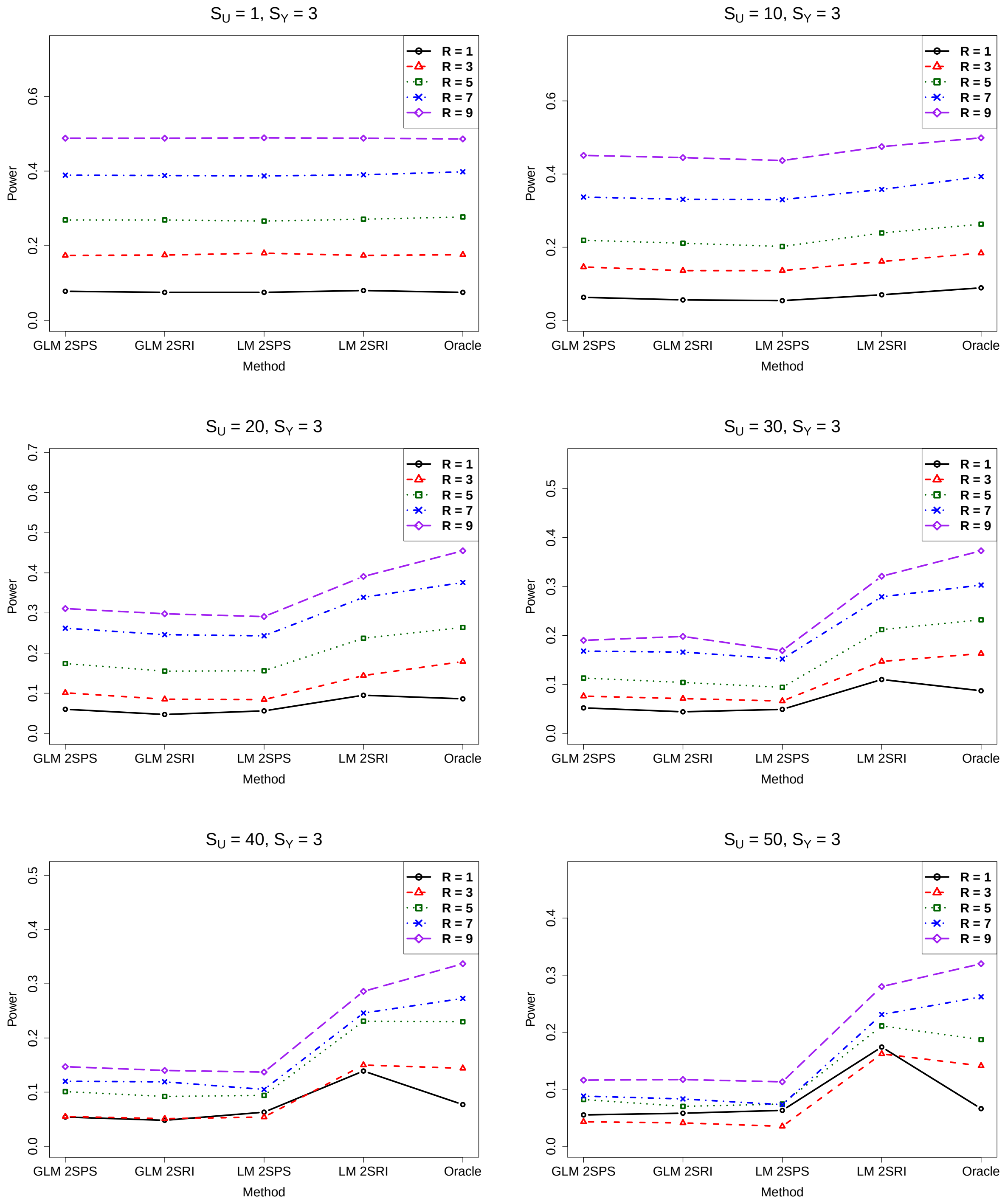

Figure S49: Simulations with the one-sample approaches: empirical power of various methods, $S_Y = 4$.

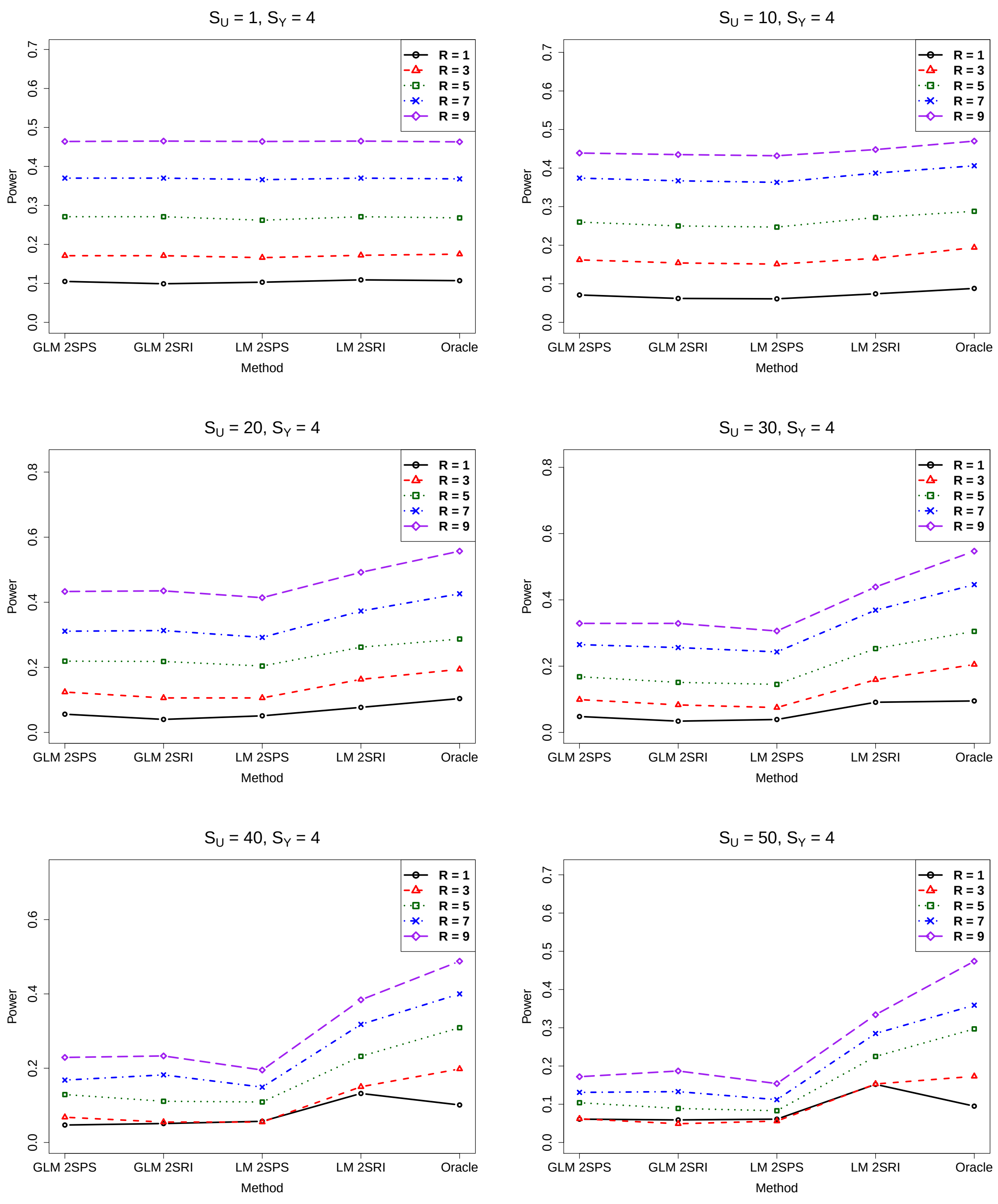

Figure S50: Simulations with the one-sample approaches: empirical power of various methods, $S_Y = 5$.

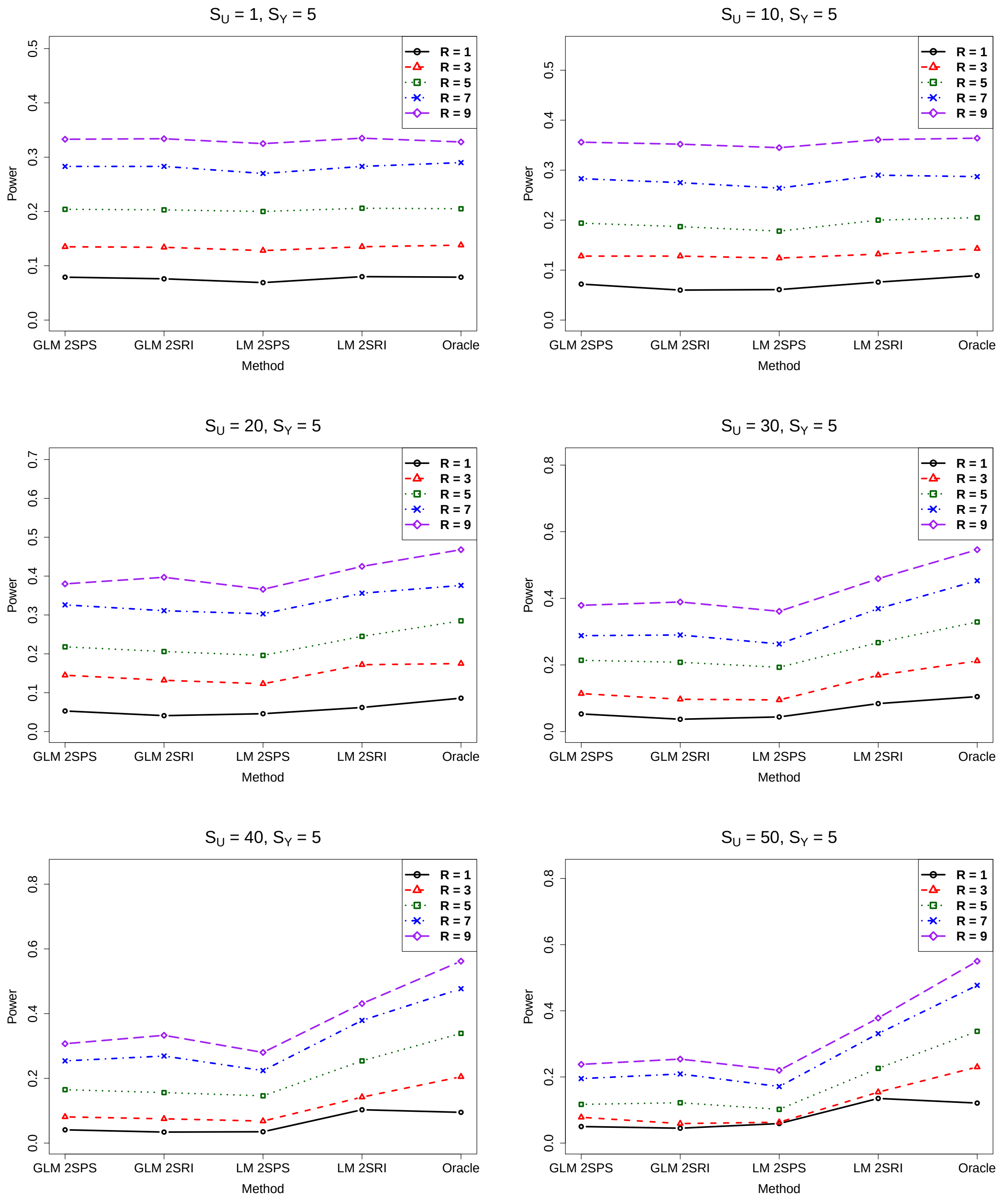

##### 2.2.3 Bias

Figure S51: Simulations with the one-sample approaches: biases of various methods, $S_Y = 0$

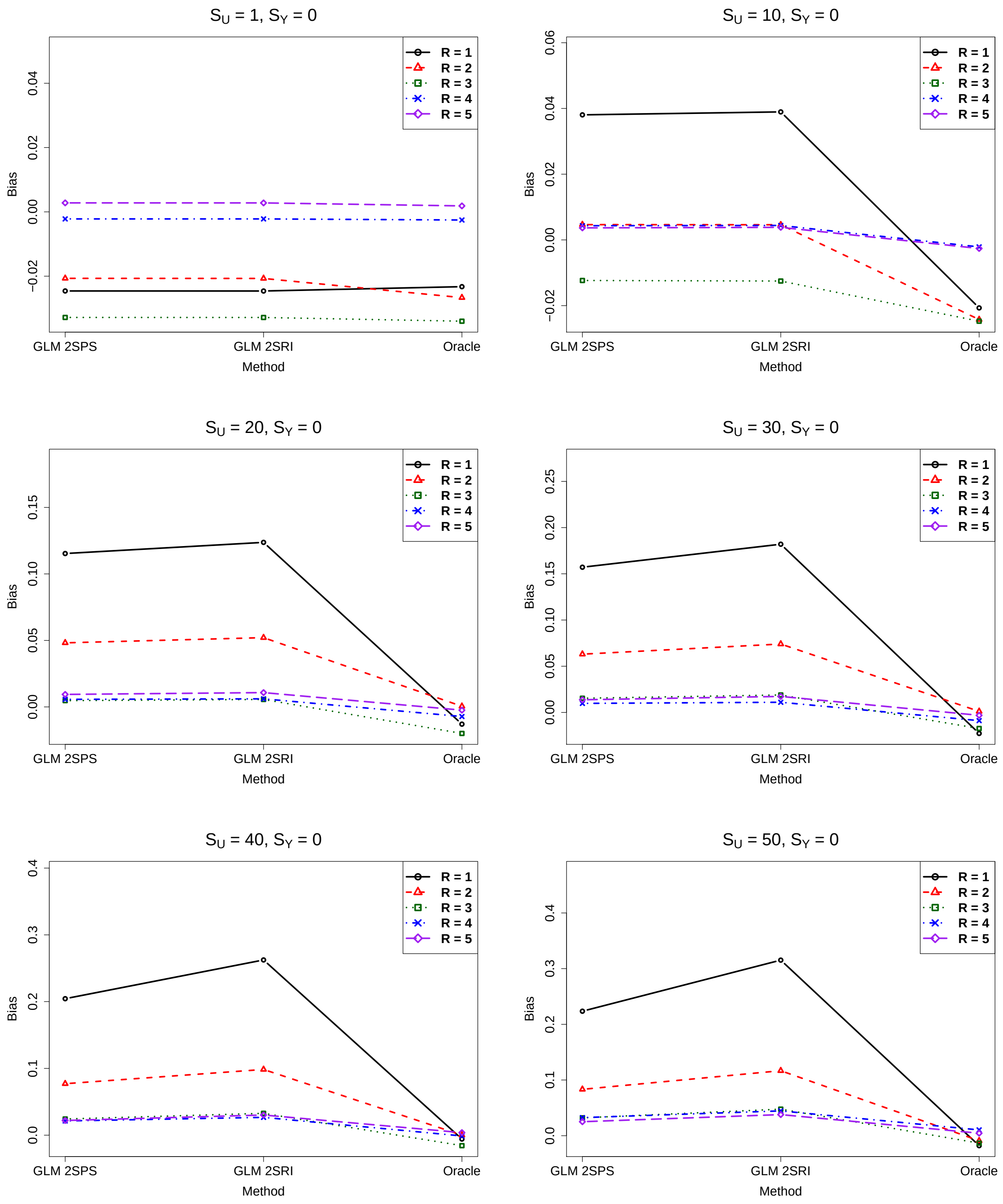

Figure S52: Simulations with the one-sample approaches: biases of various methods, $S_Y = 1$

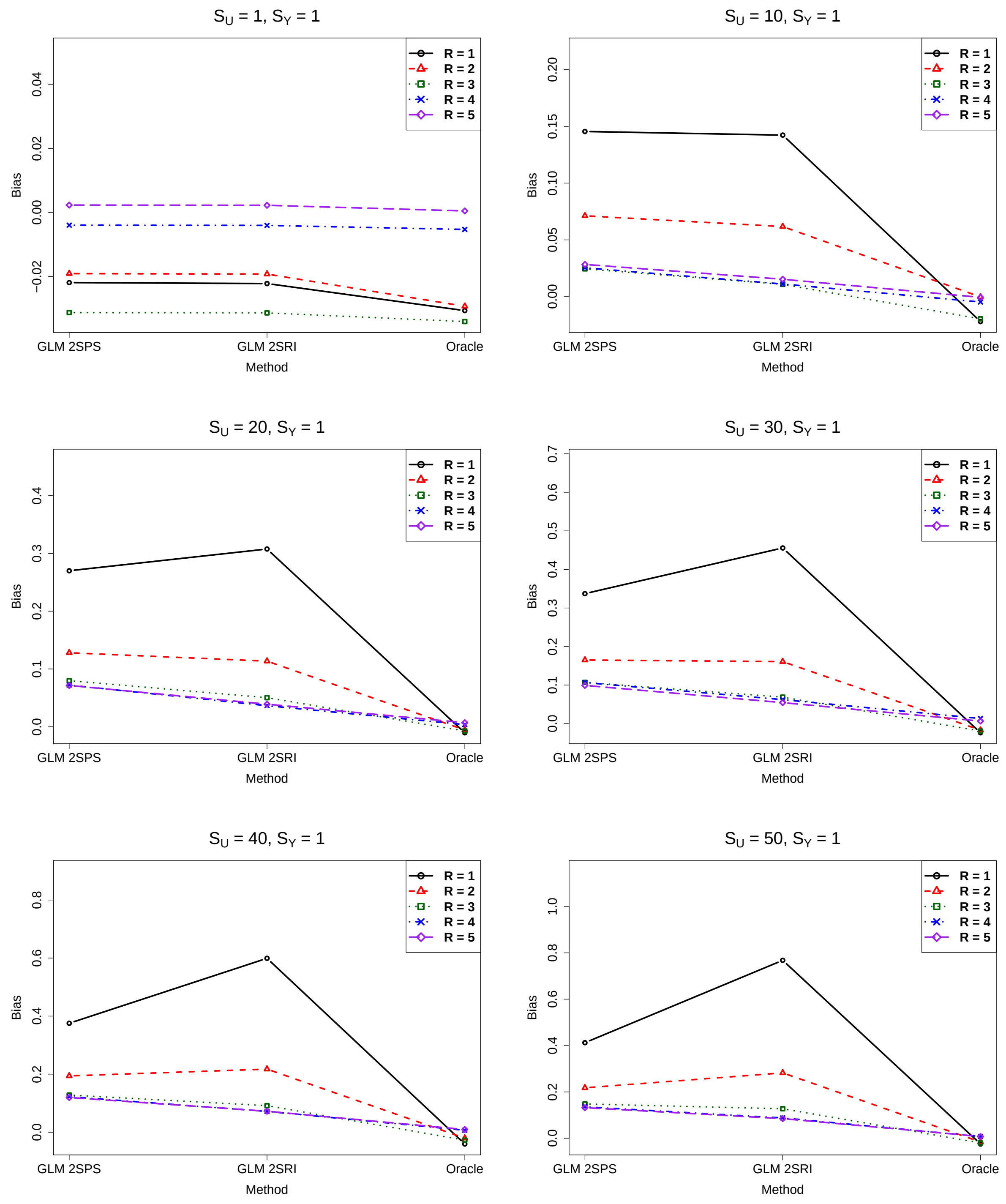

Figure S53: Simulations with the one-sample approaches: biases of various methods, $S_Y = 2$

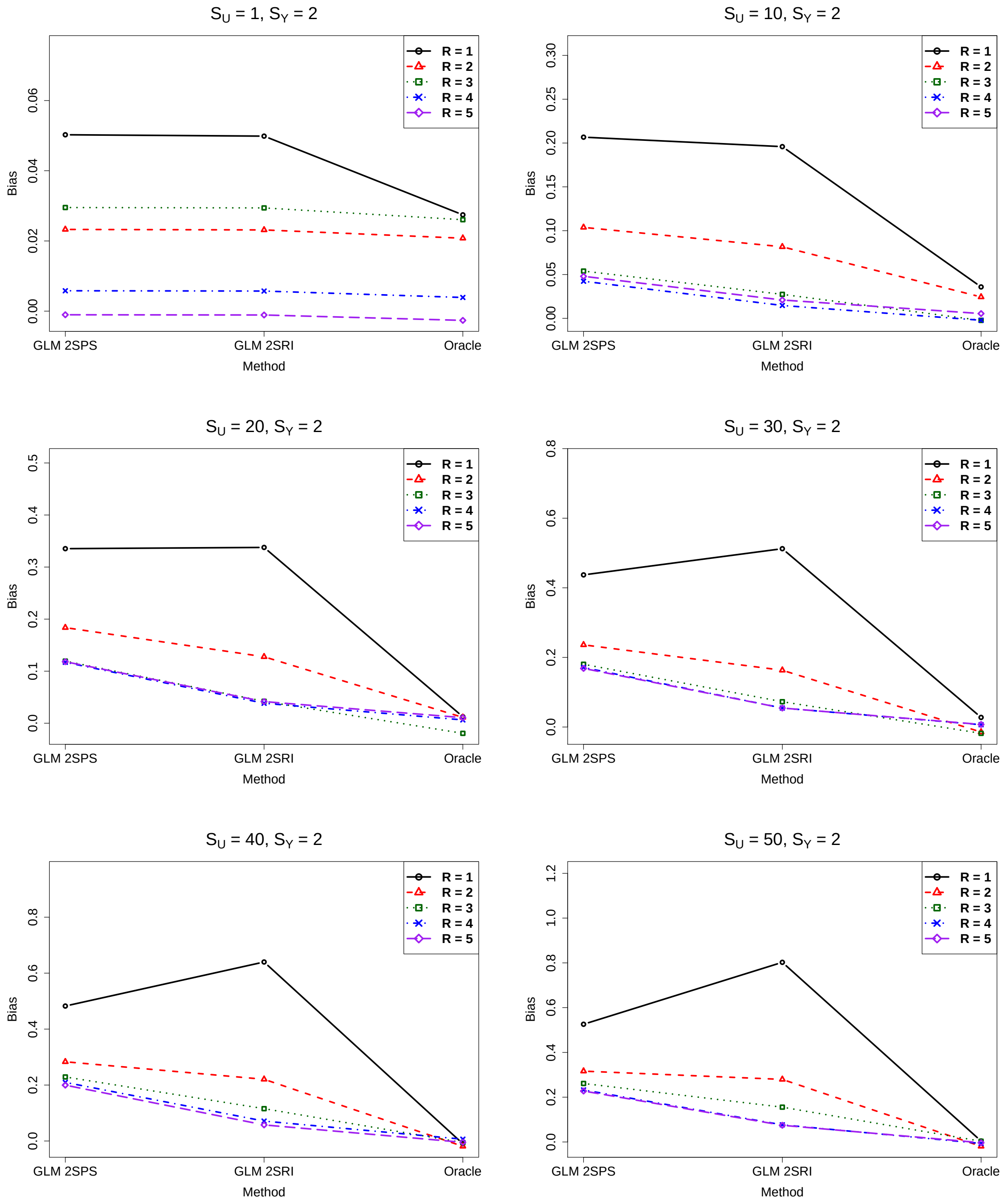

Figure S54: Simulations with the one-sample approaches: biases of various methods, $S_Y = 3$

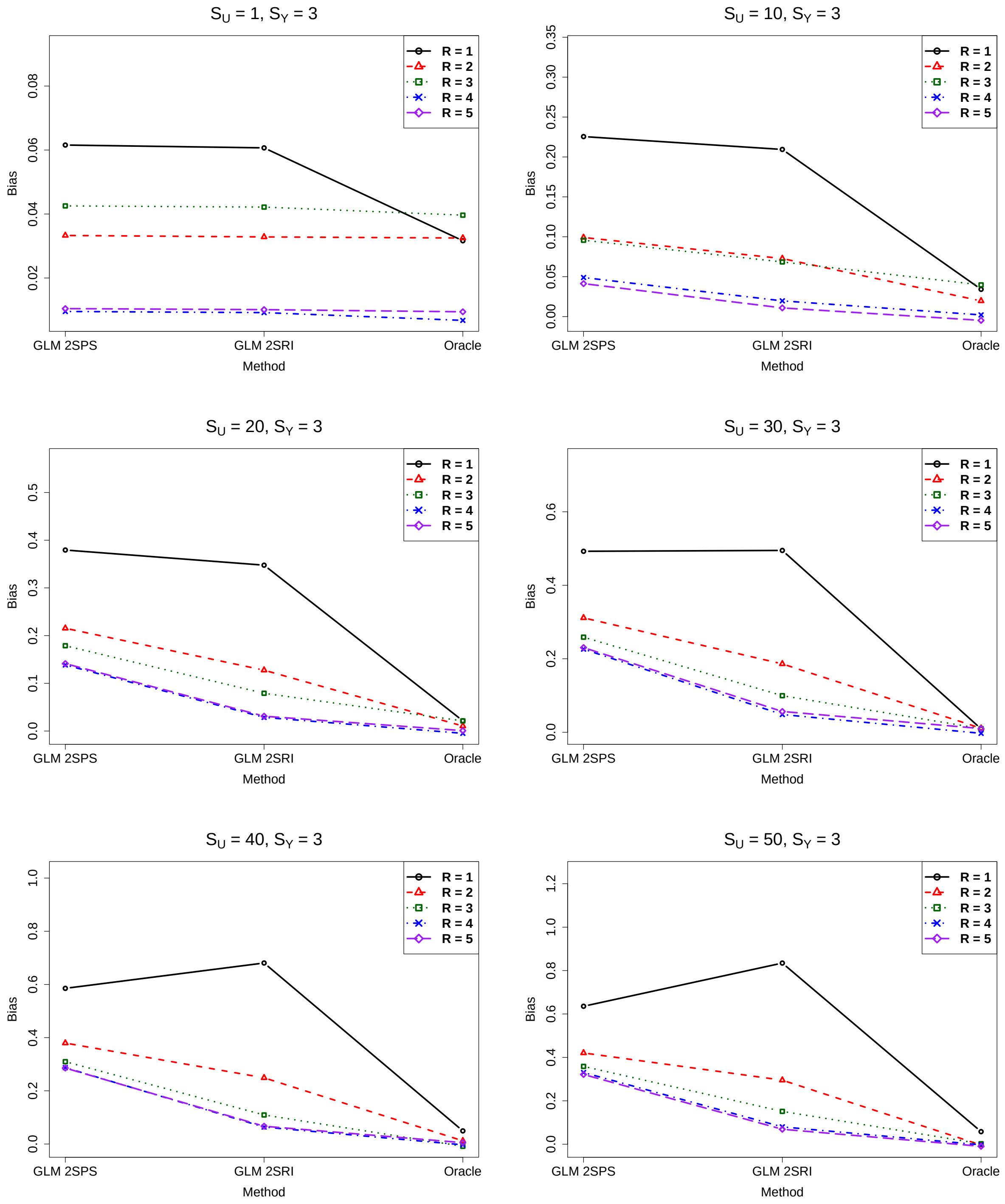

Figure S55: Simulations with the one-sample approaches: biases of various methods, $S_Y = 4$

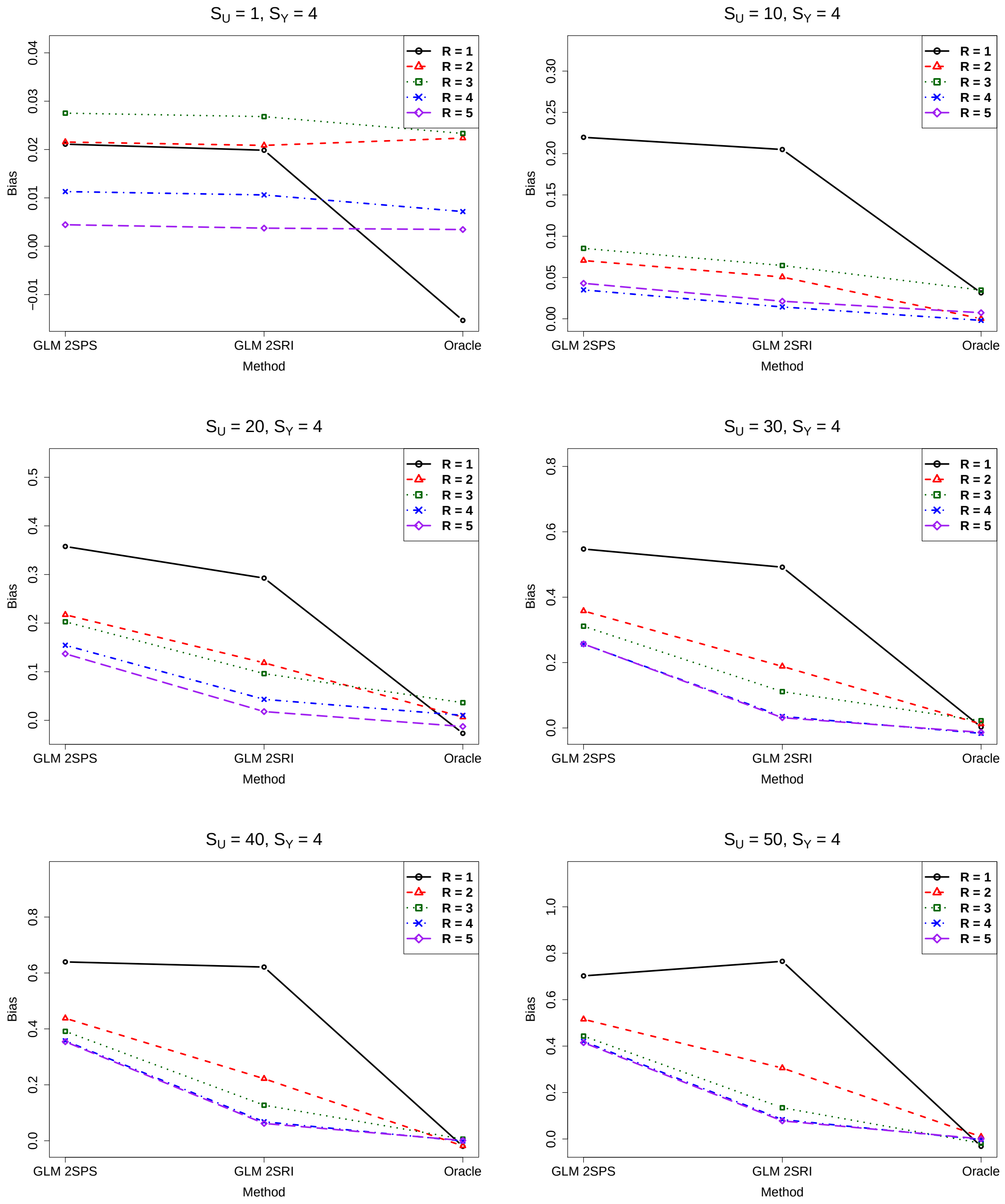

Figure S56: Simulations with the one-sample approaches: biases of various methods, $S_Y = 5$

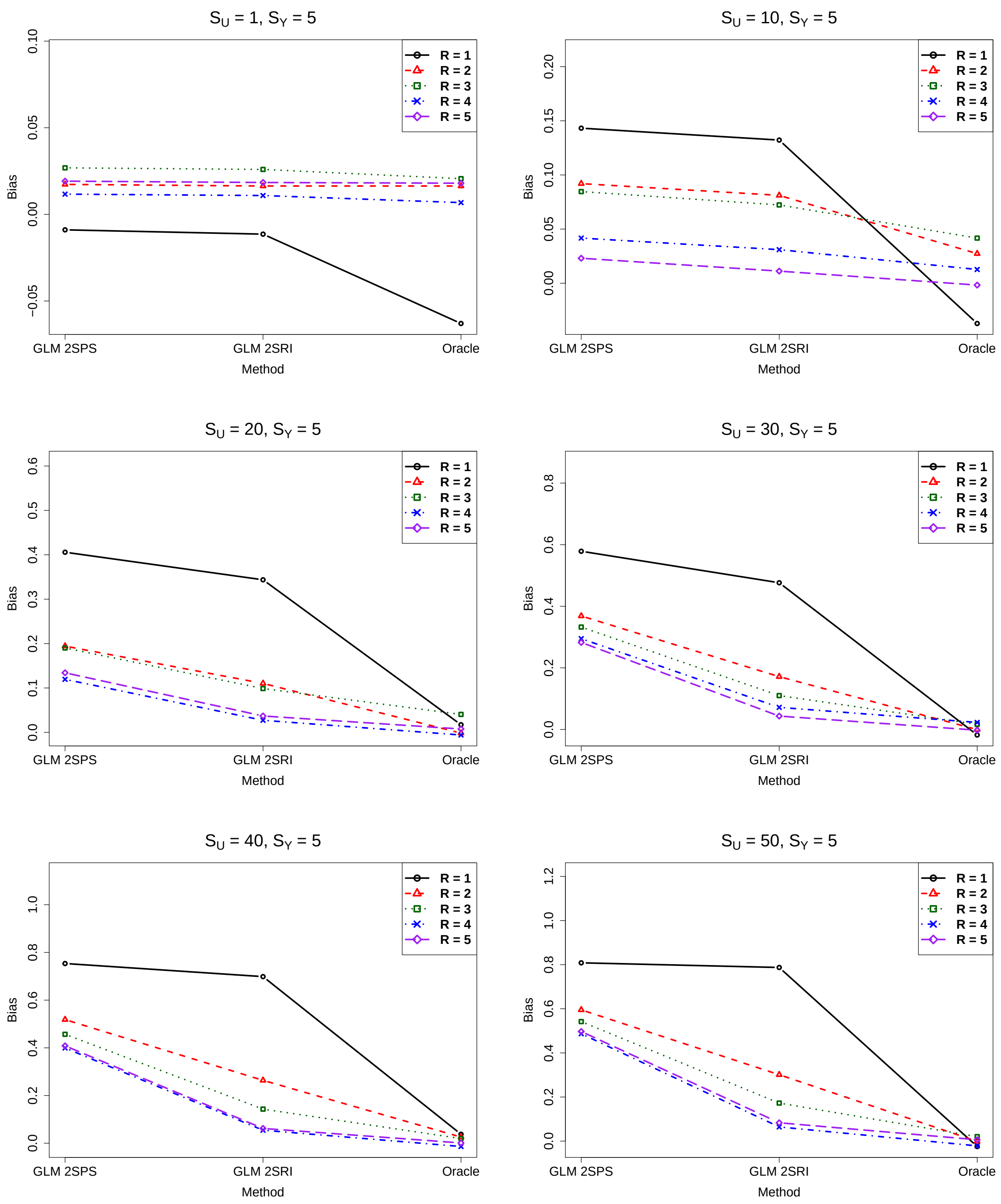

##### 2.2.4 Comparison of the original and corrected standard error estimates

Figure S57: Simulations with the one-sample approaches: comparison of the original and corrected standard error estimates, $S_Y = 0, R = 1$.

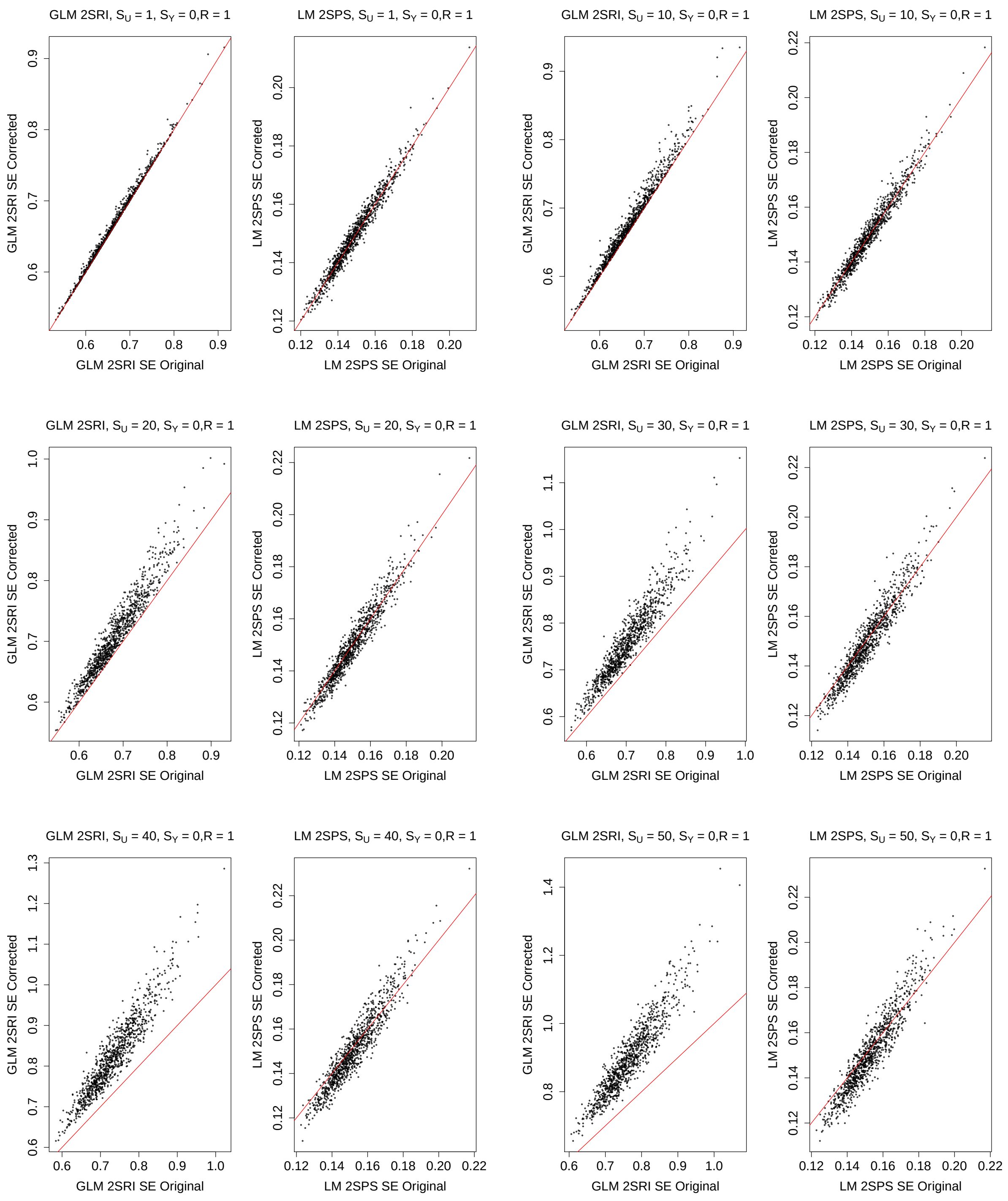

Figure S58: Simulations with the one-sample approaches: comparison of the original and corrected standard error estimates, $S_Y = 1, R = 1$.

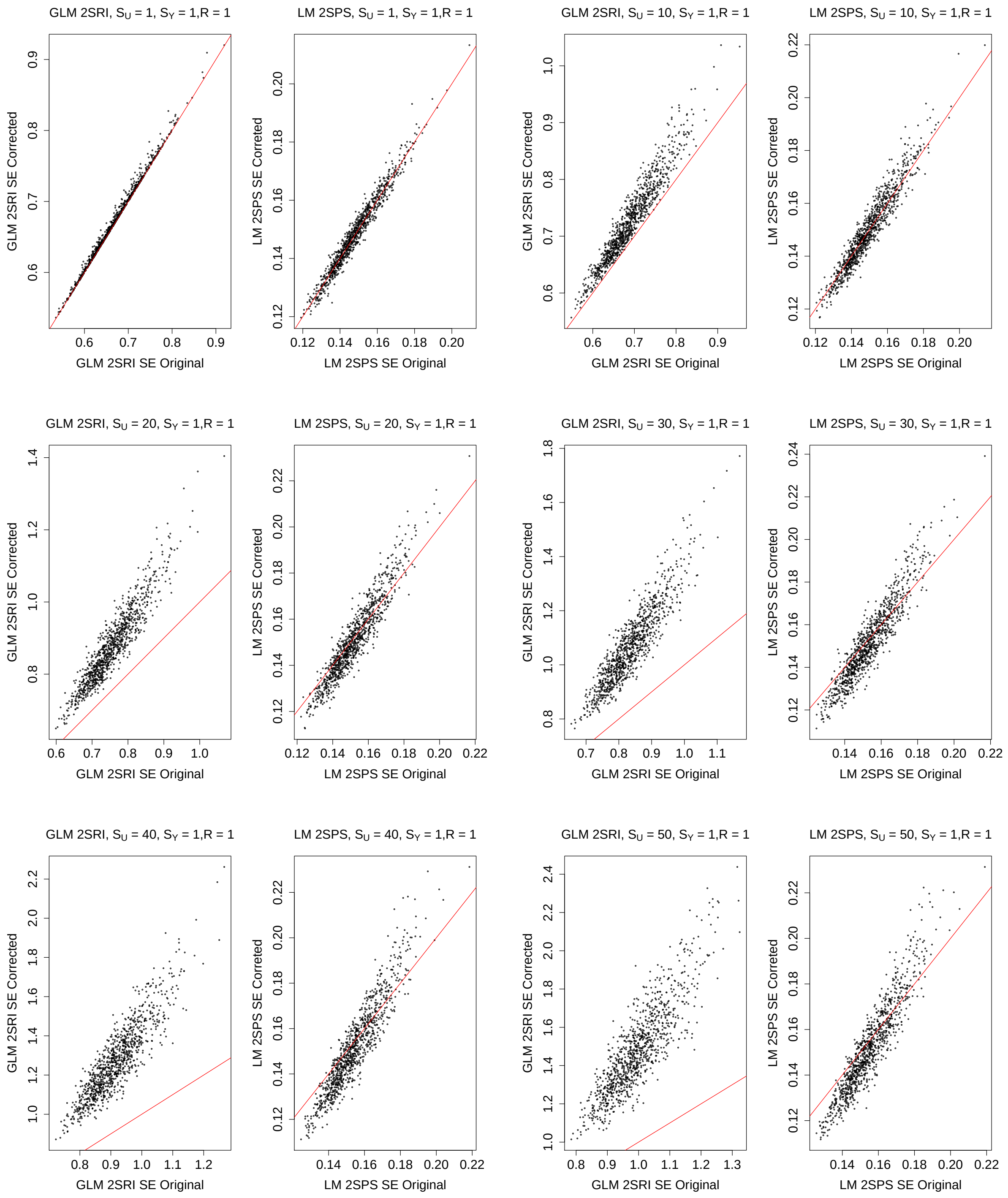

Figure S59: Simulations with the one-sample approaches: comparison of the original and corrected standard error estimates, $S_Y = 2, R = 1$.

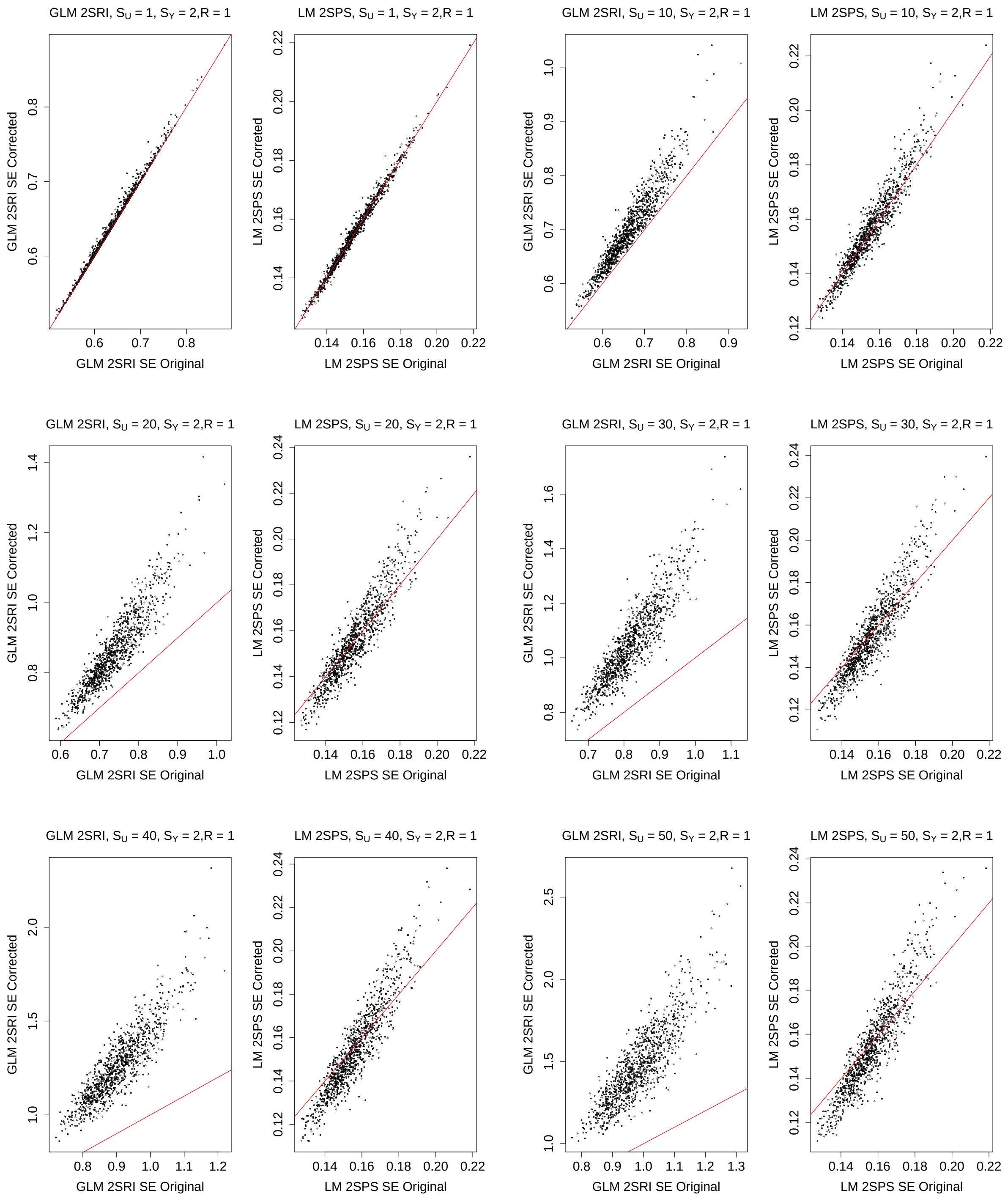

Figure S60: Simulations with the one-sample approaches: comparison of the original and corrected standard error estimates, $S_Y = 3, R = 1$.

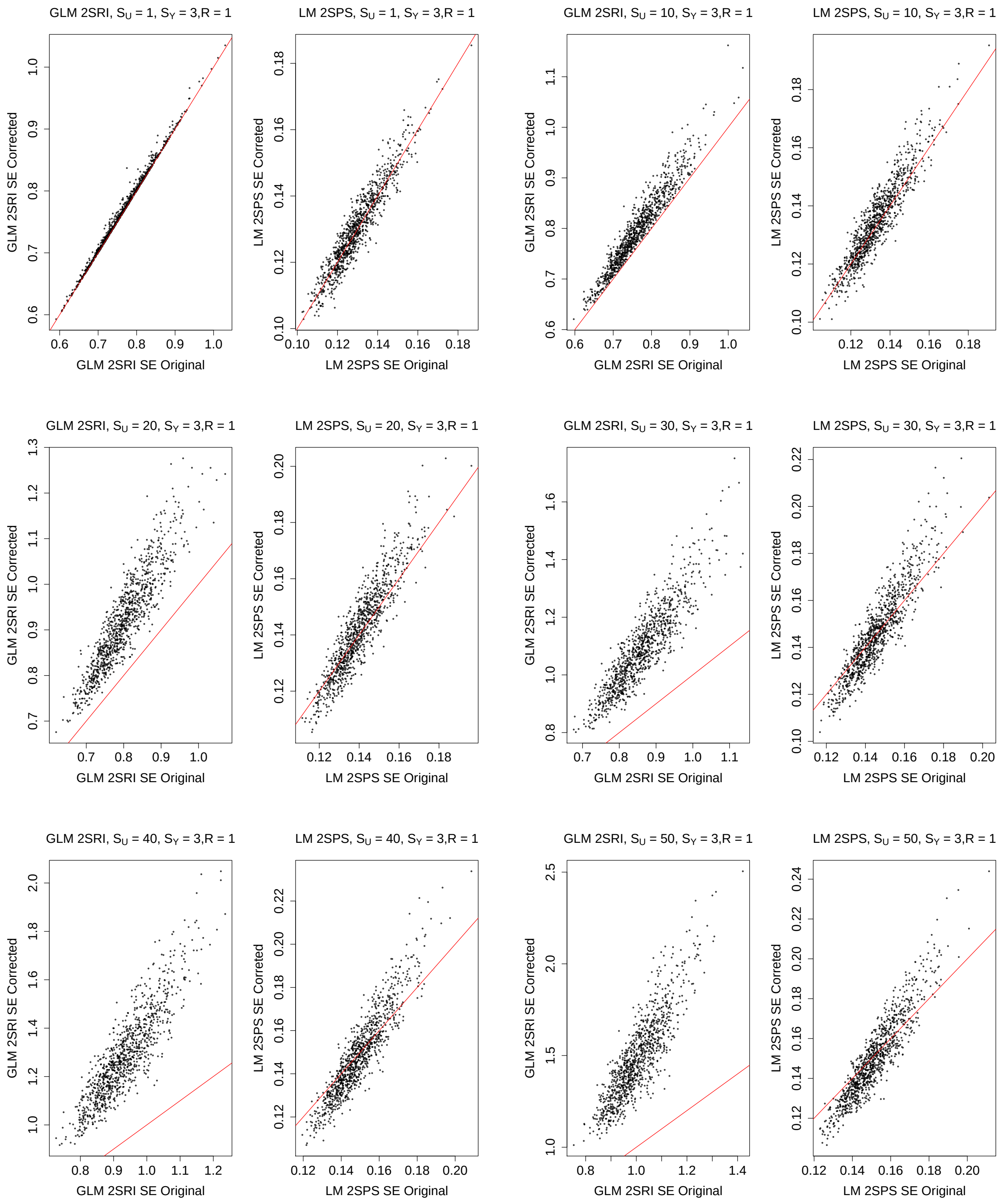

Figure S61: Simulations with the one-sample approaches: comparison of the original and corrected standard error estimates, $S_Y = 4, R = 1$.

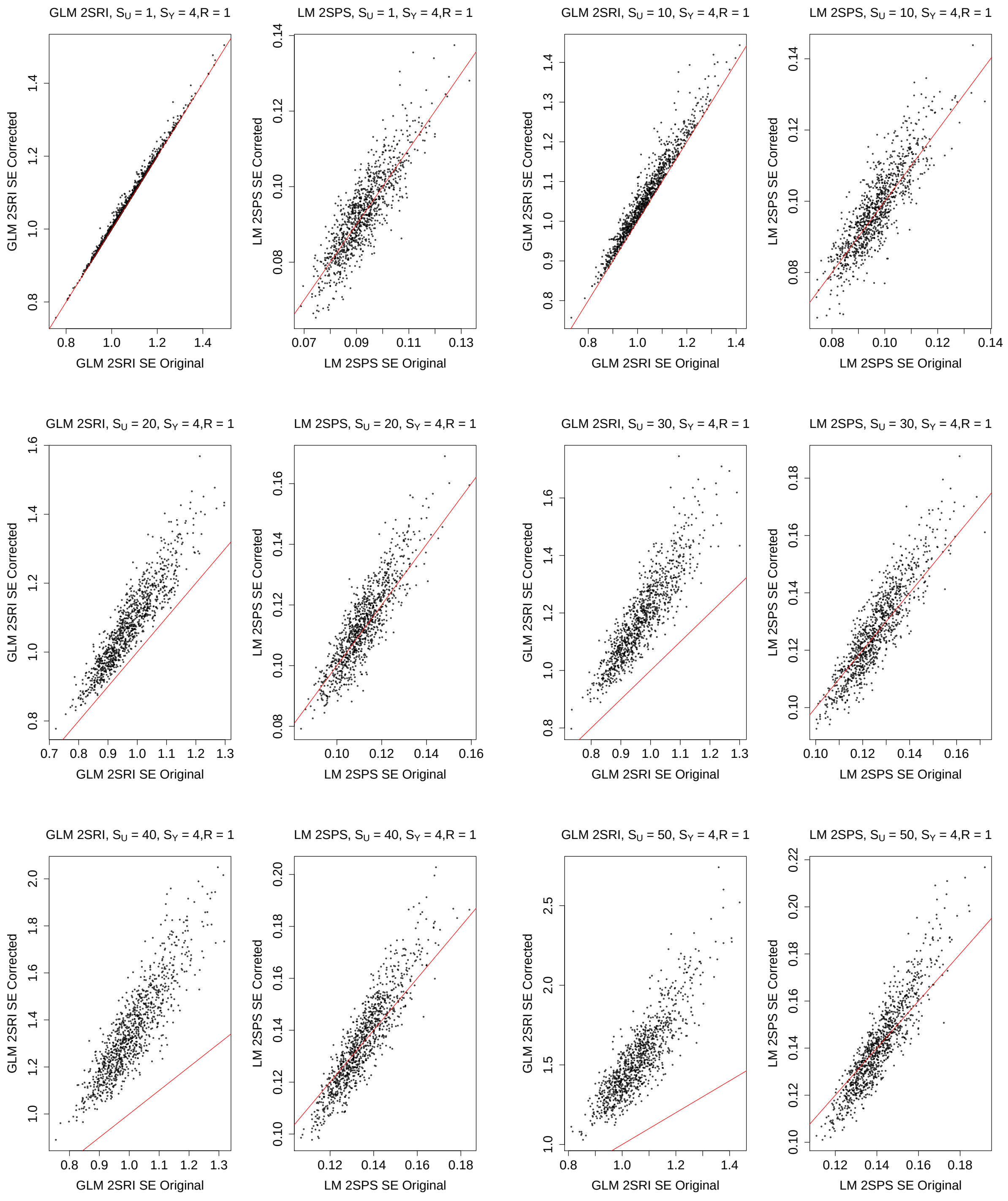

Figure S62: Simulations with the one-sample approaches: comparison of the original and corrected standard error estimates, $S_Y = 5, R = 1$.

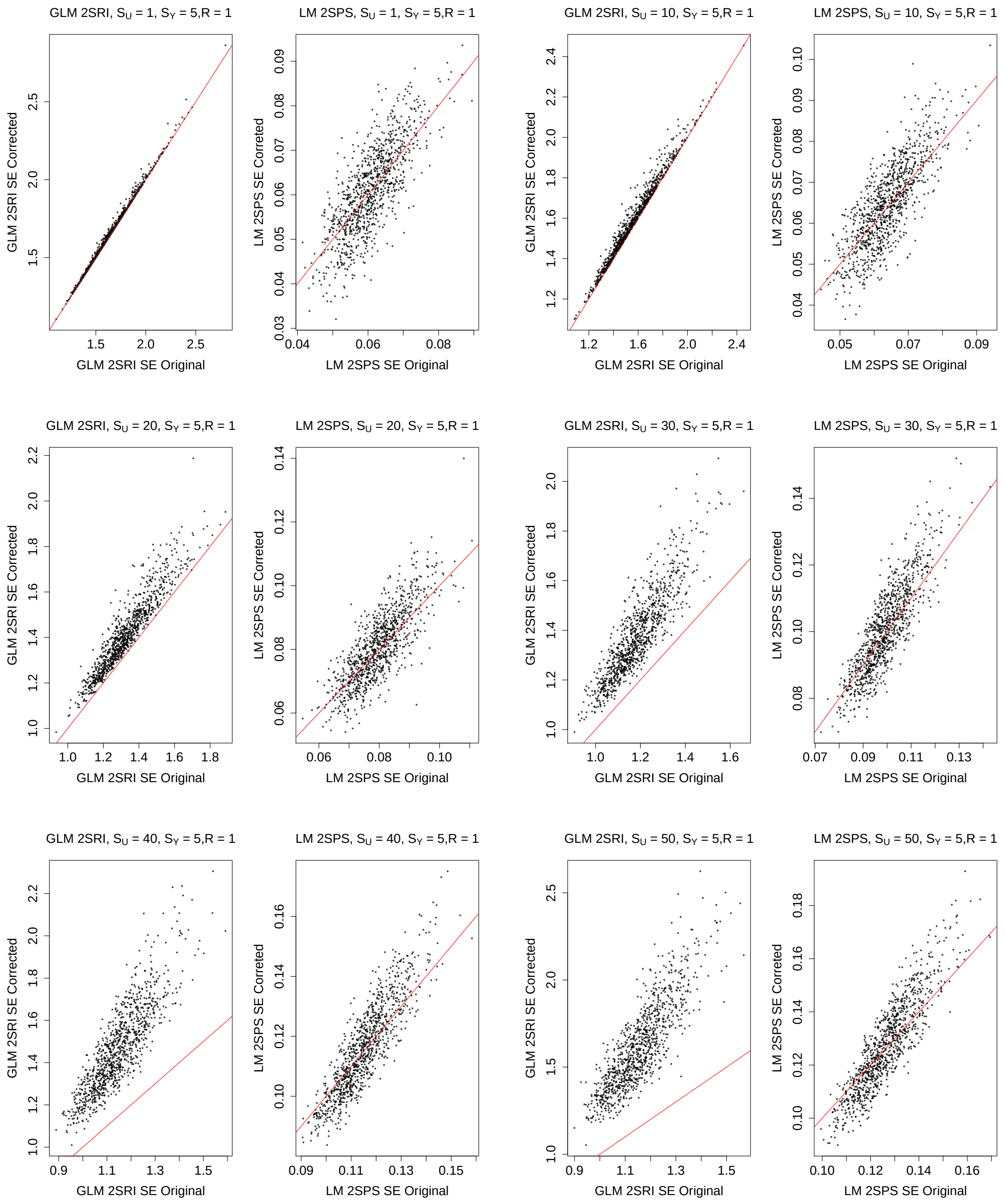

Figure S63: Simulations with the one-sample approaches: comparison of the original and corrected standard error estimates, $S_Y = 0, R = 3$.

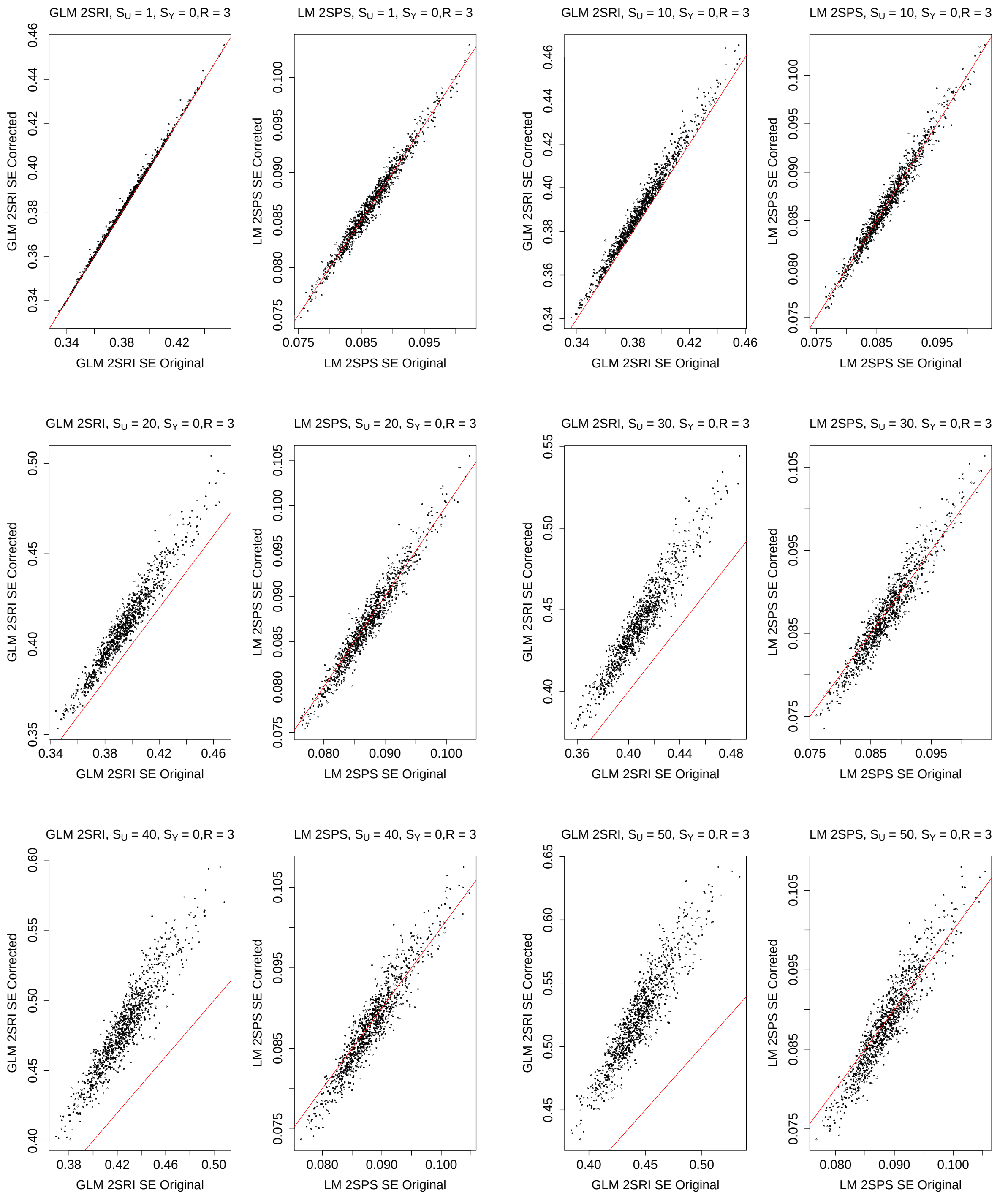

Figure S64: Simulations with the one-sample approaches: comparison of the original and corrected standard error estimates, $S_Y = 1, R = 3$.

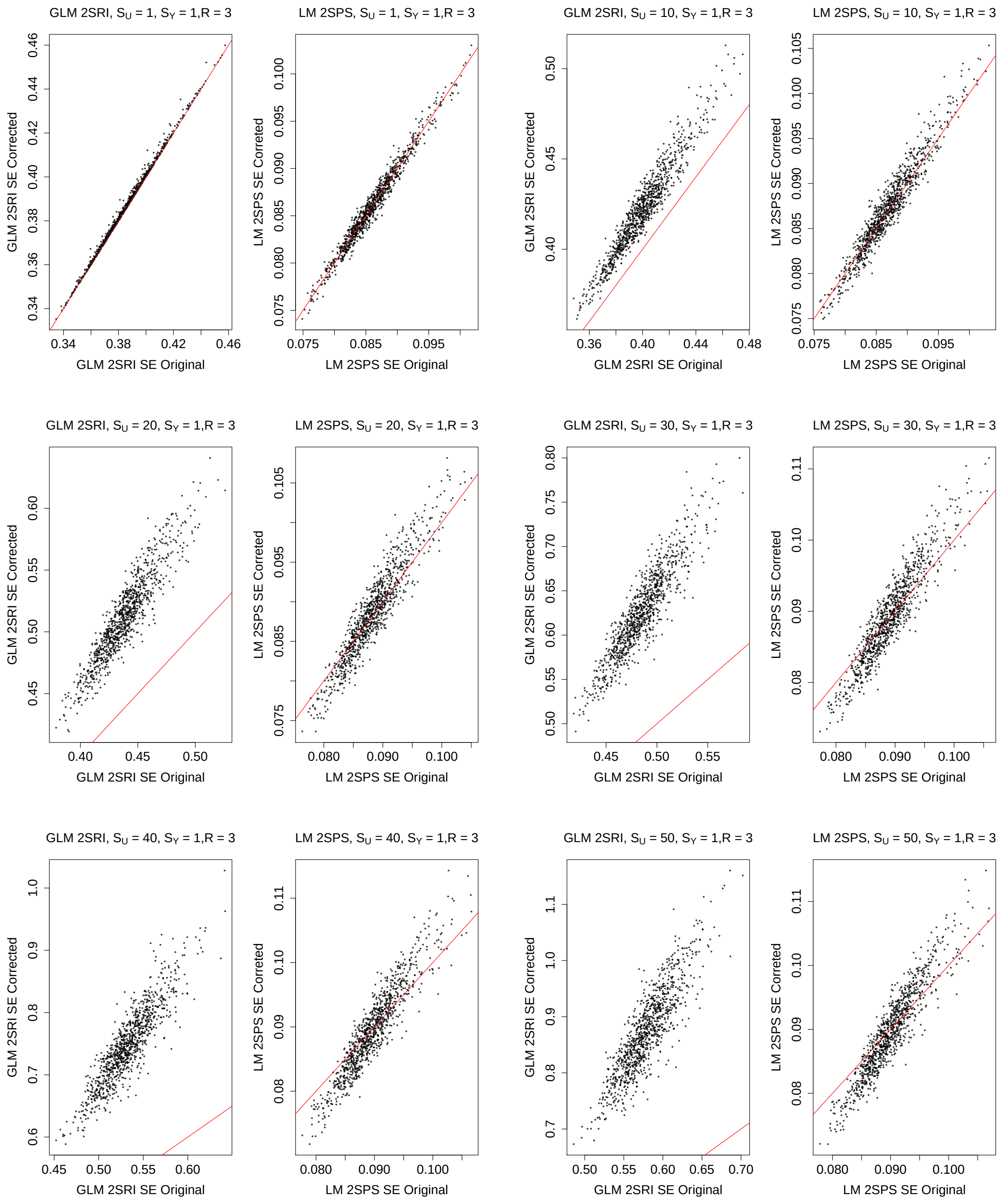

Figure S65: Simulations with the one-sample approaches: comparison of the original and corrected standard error estimates, $S_Y = 2, R = 3$.

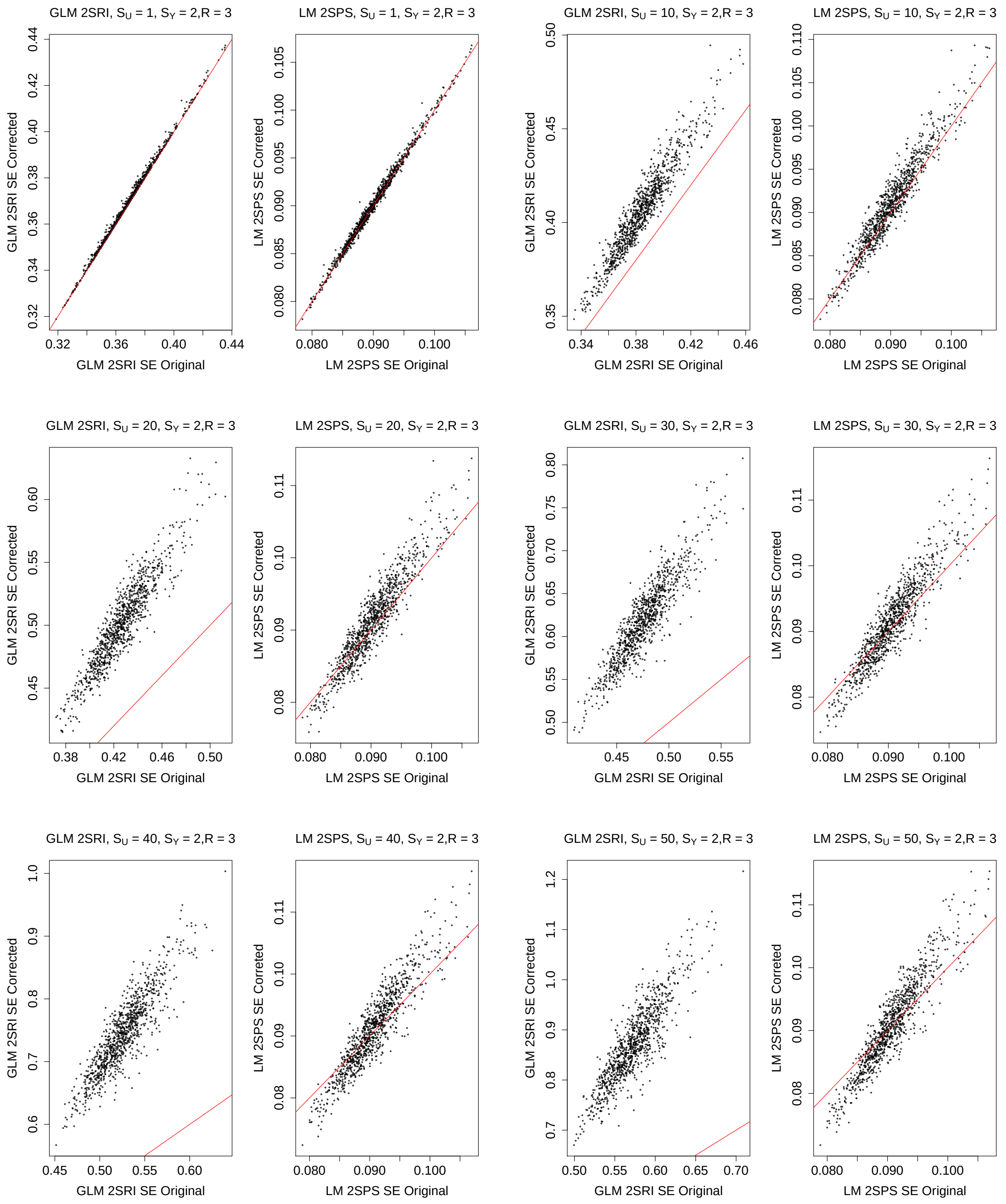

Figure S66: Simulations with the one-sample approaches: comparison of the original and corrected standard error estimates, $S_Y = 3, R = 3$.

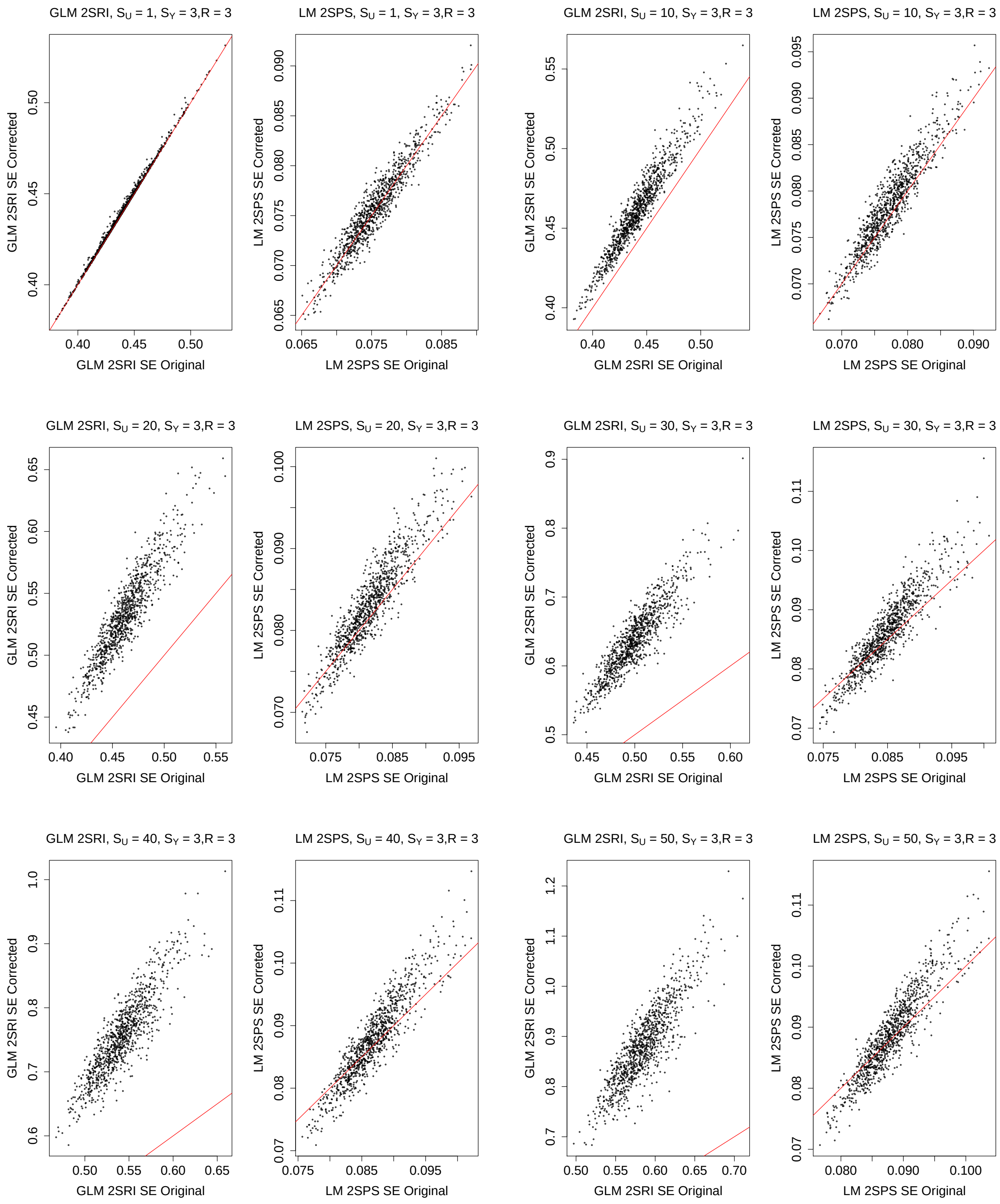

Figure S67: Simulations with the one-sample approaches: comparison of the original and corrected standard error estimates, $S_Y = 4, R = 3$.

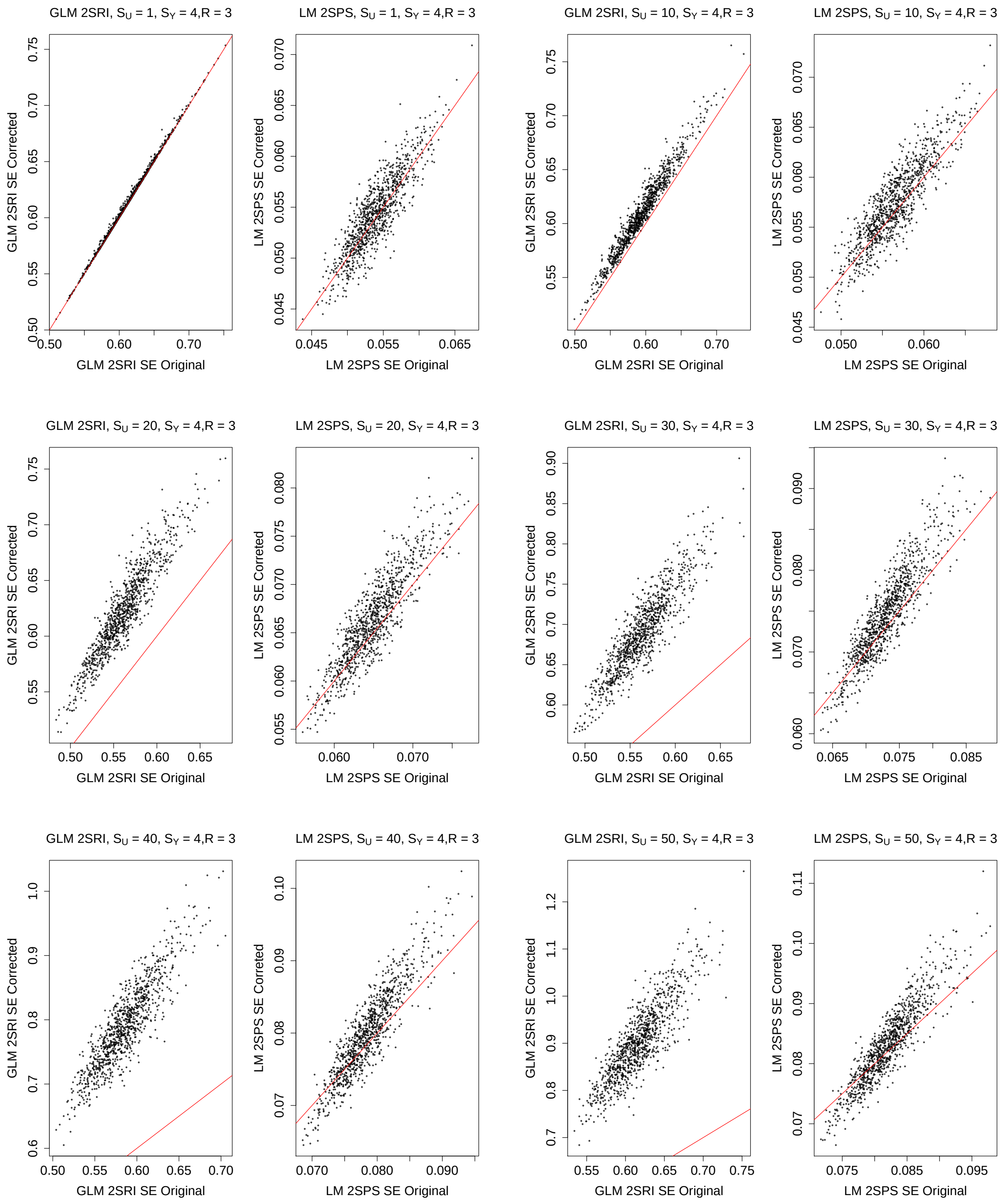

Figure S68: Simulations with the one-sample approaches: comparison of the original and corrected standard error estimates, $S_Y = 5, R = 3$.

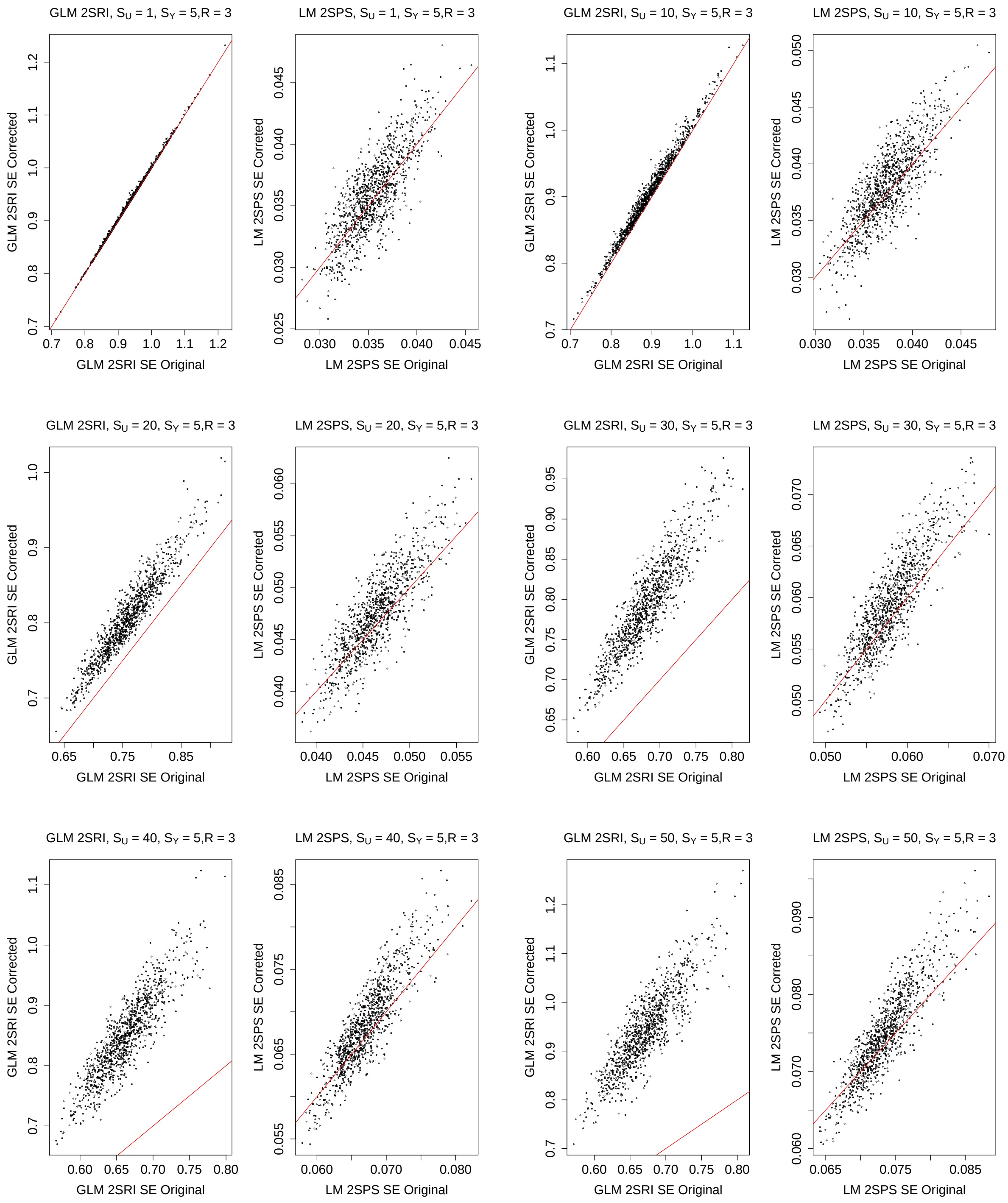

Figure S69: Simulations with the one-sample approaches: comparison of the original and corrected standard error estimates, $S_Y = 0, R = 5$.

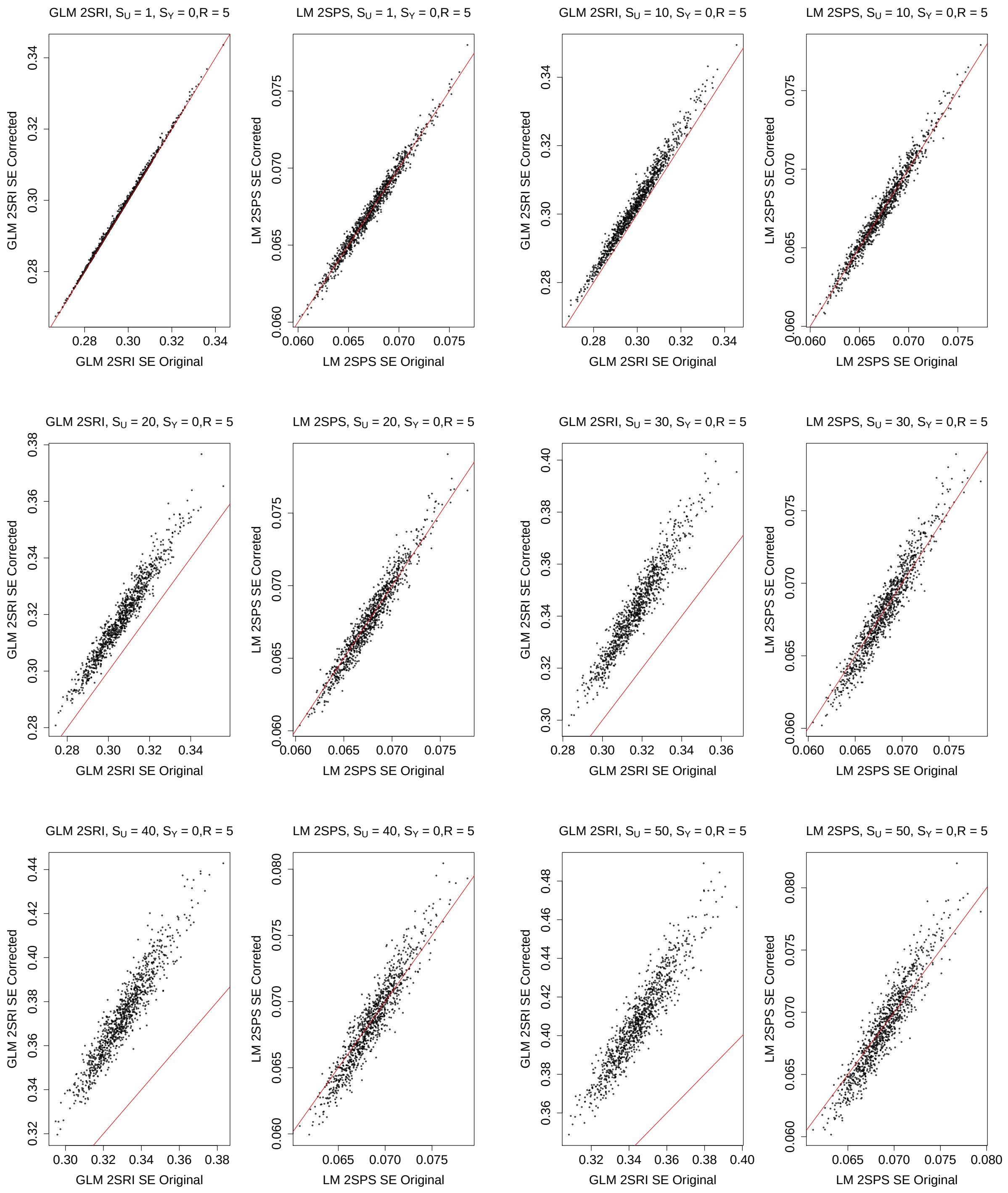

Figure S70: Simulations with the one-sample approaches: comparison of the original and corrected standard error estimates, $S_Y = 1, R = 5$.

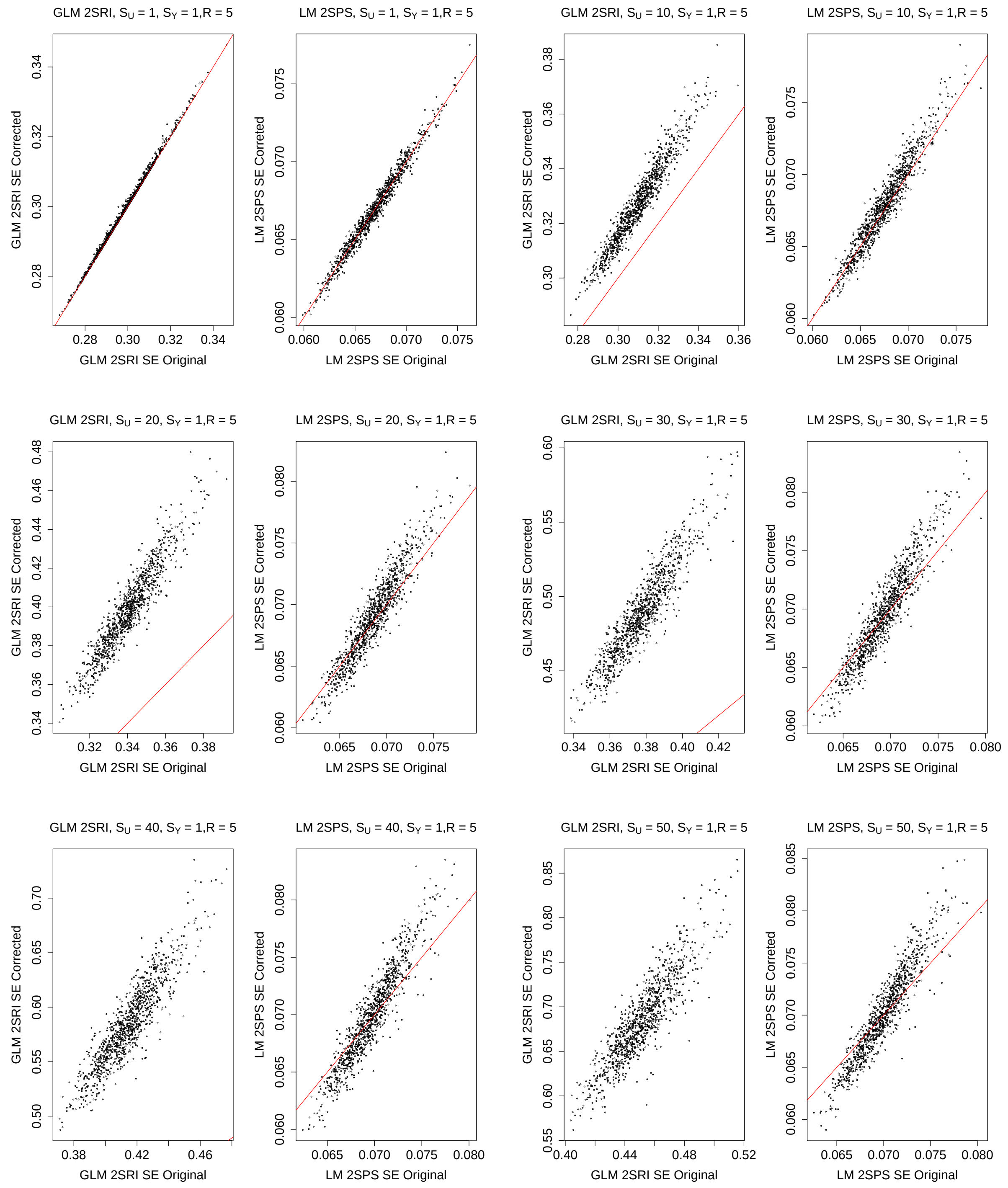

Figure S71: Simulations with the one-sample approaches: comparison of the original and corrected standard error estimates, $S_Y = 2, R = 5$.

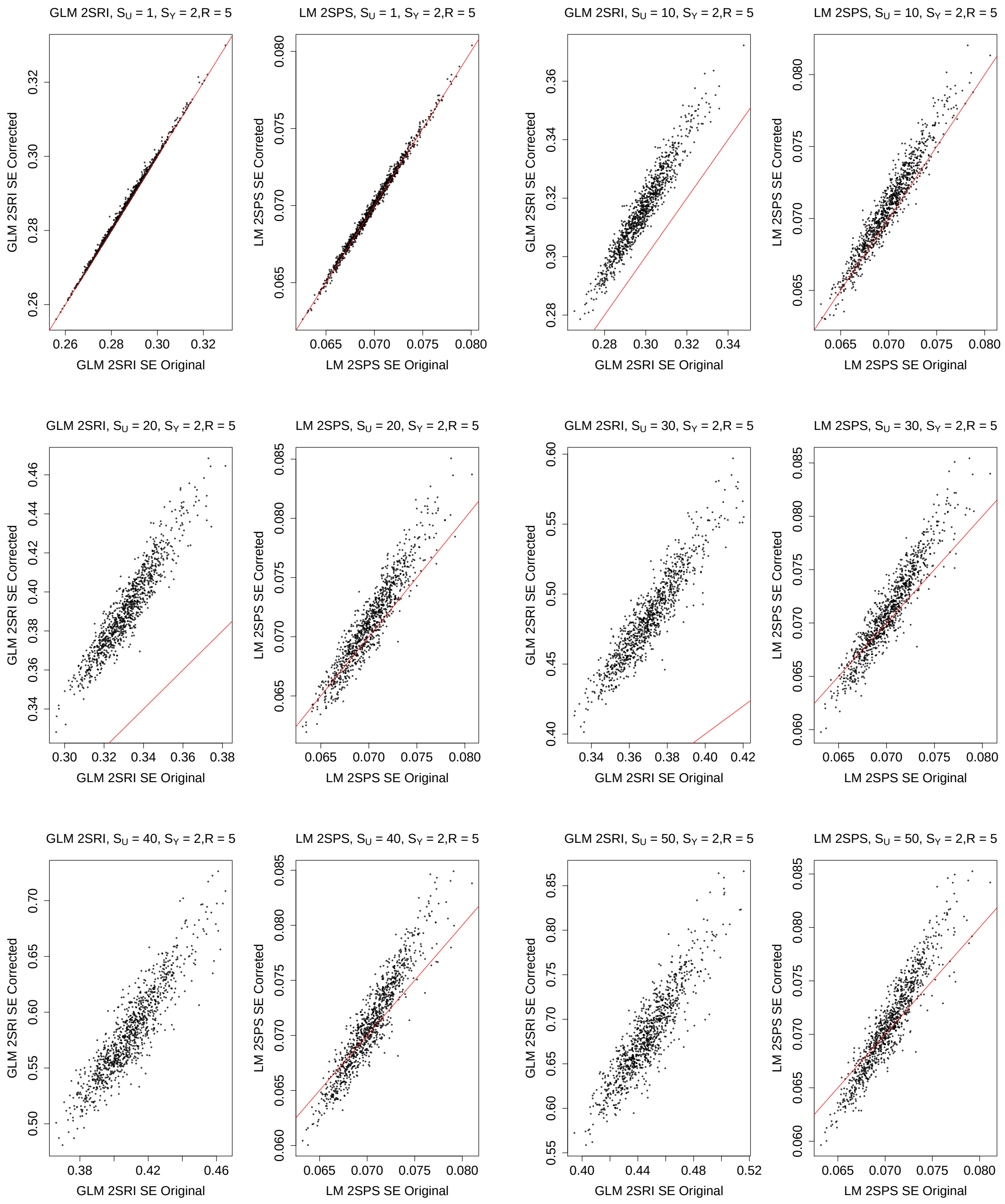

Figure S72: Simulations with the one-sample approaches: comparison of the original and corrected standard error estimates, $S_Y = 3, R = 5$.

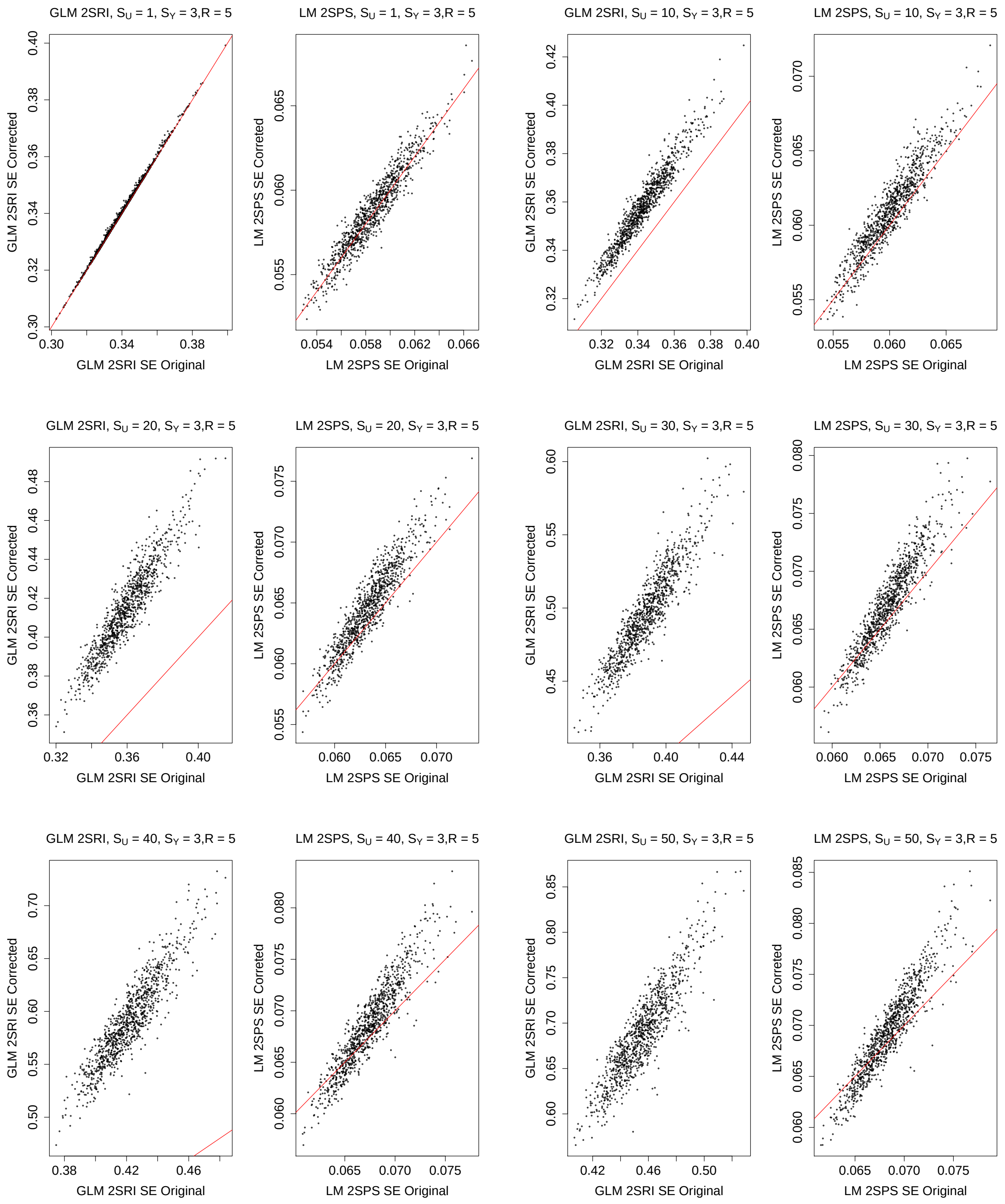

Figure S73: Simulations with the one-sample approaches: comparison of the original and corrected standard error estimates, $S_Y = 4, R = 5$.

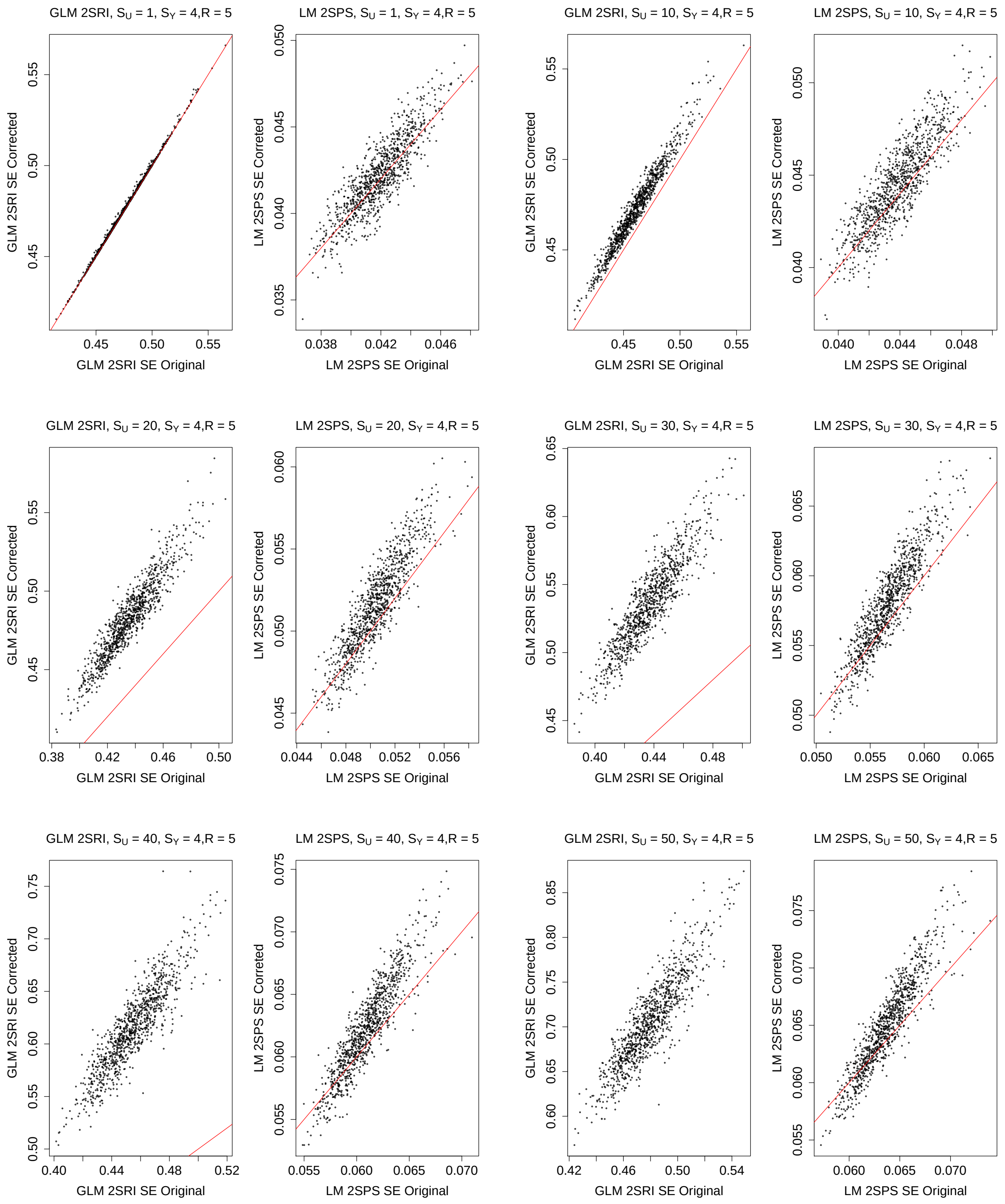

Figure S74: Simulations with the one-sample approaches: comparison of the original and corrected standard error estimates, $S_Y = 5, R = 5$.

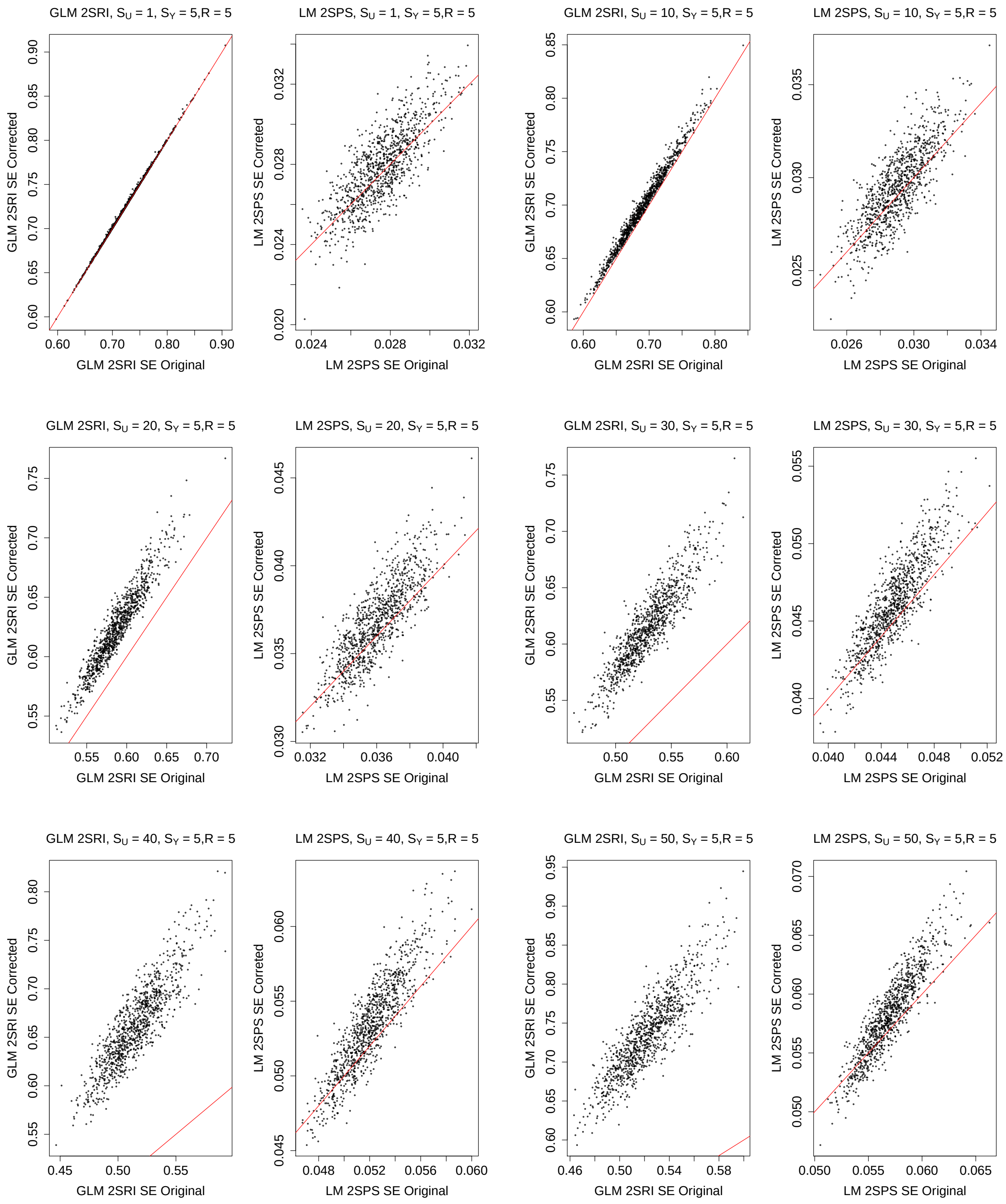

Figure S75: Simulations with the one-sample approaches: comparison of the original and corrected standard error estimates, $S_Y = 0, R = 7$.

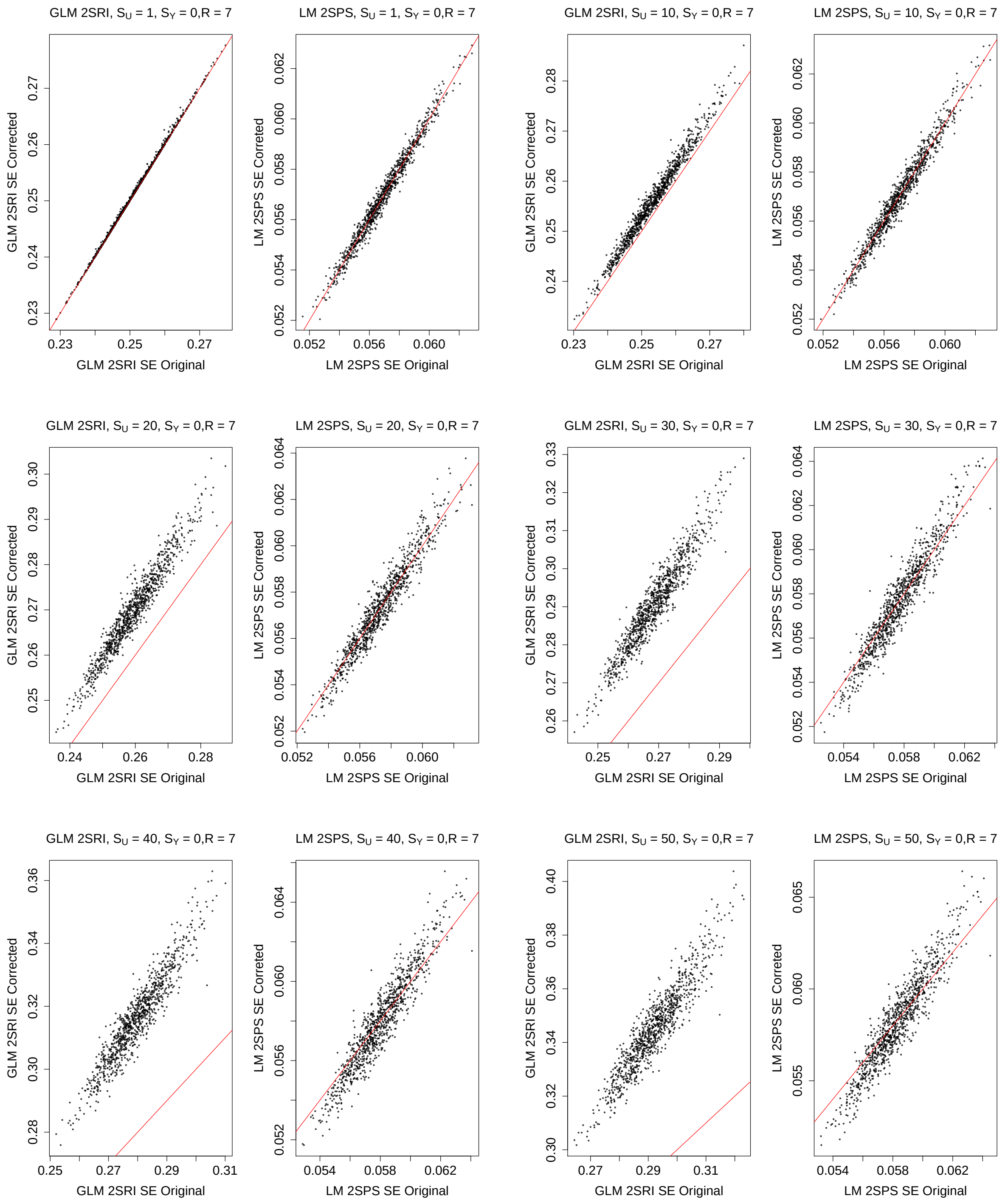

Figure S76: Simulations with the one-sample approaches: comparison of the original and corrected standard error estimates, $S_Y = 1, R = 7$.

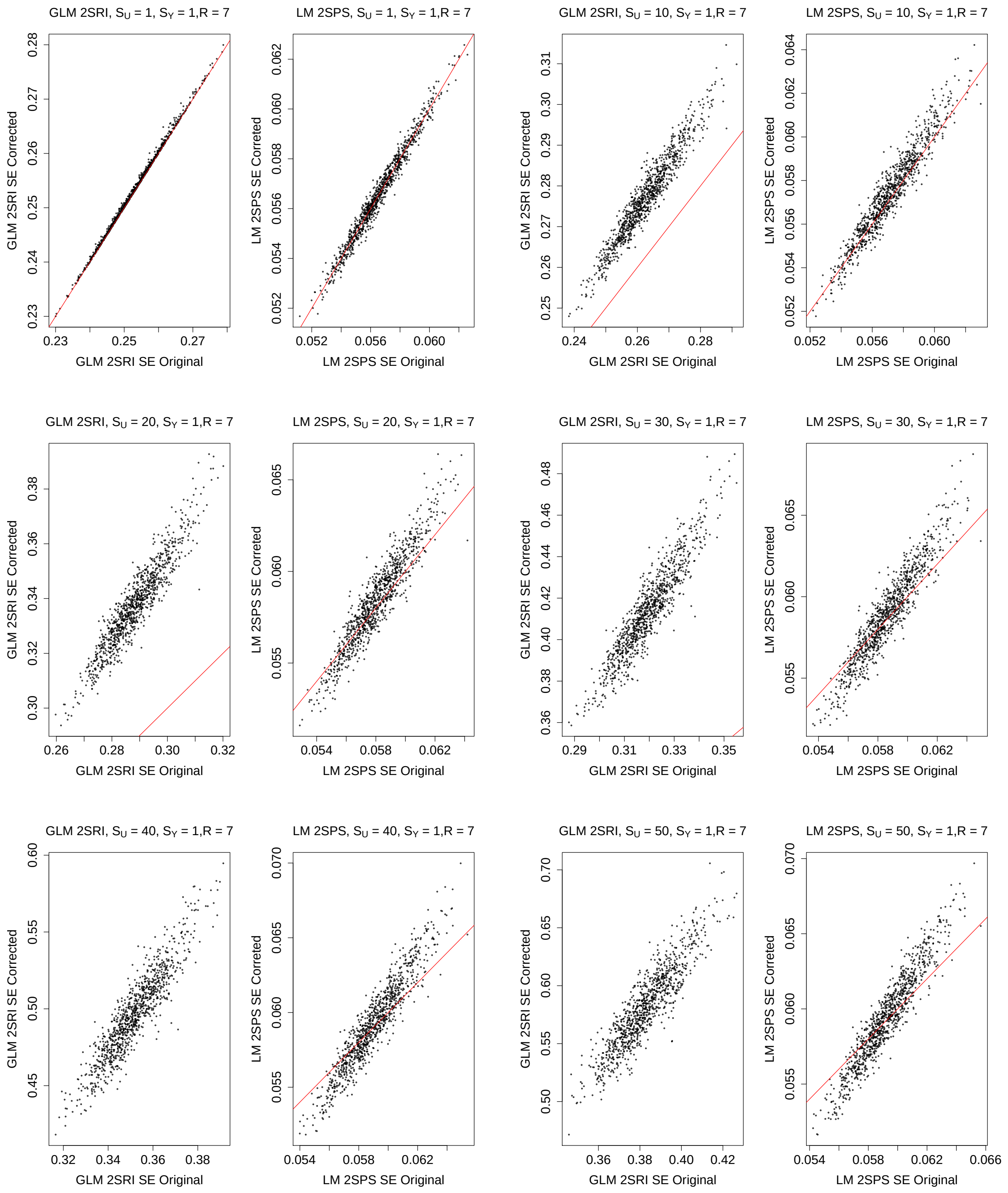

Figure S77: Simulations with the one-sample approaches: comparison of the original and corrected standard error estimates, $S_Y = 2, R = 7$.

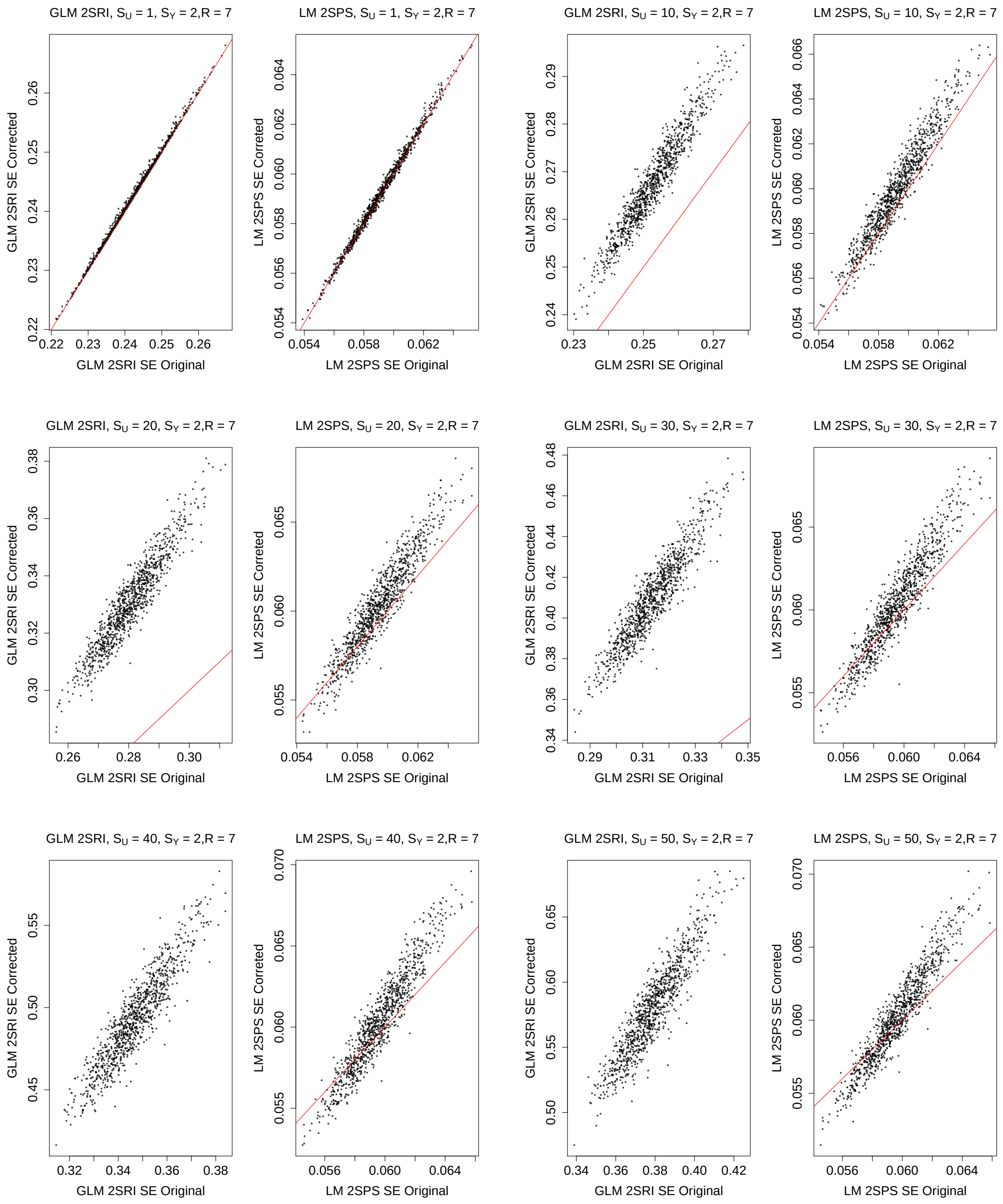

Figure S78: Simulations with the one-sample approaches: comparison of the original and corrected standard error estimates, $S_Y = 3, R = 7$.

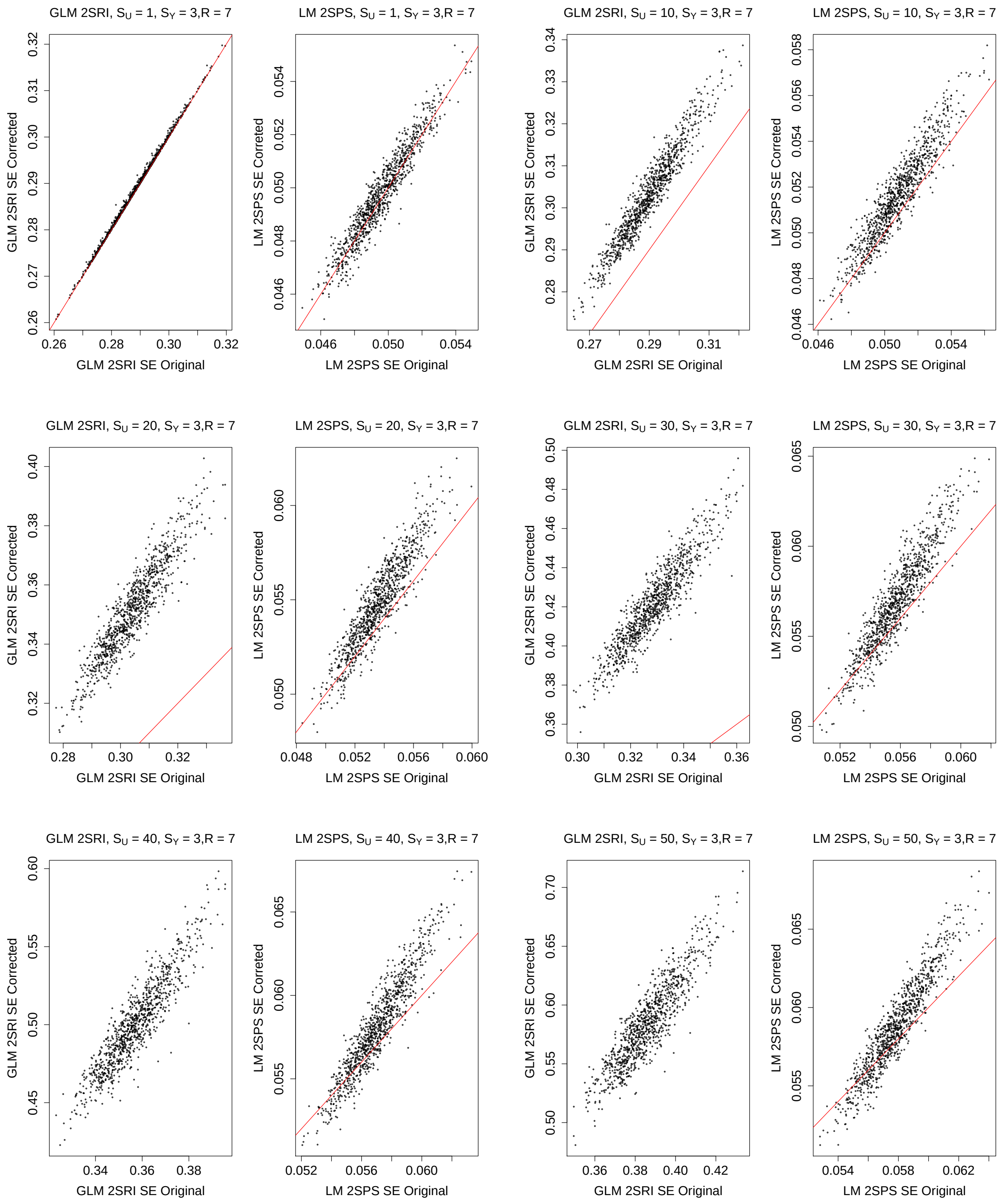

Figure S79: Simulations with the one-sample approaches: comparison of the original and corrected standard error estimates, $S_Y = 4, R = 7$.

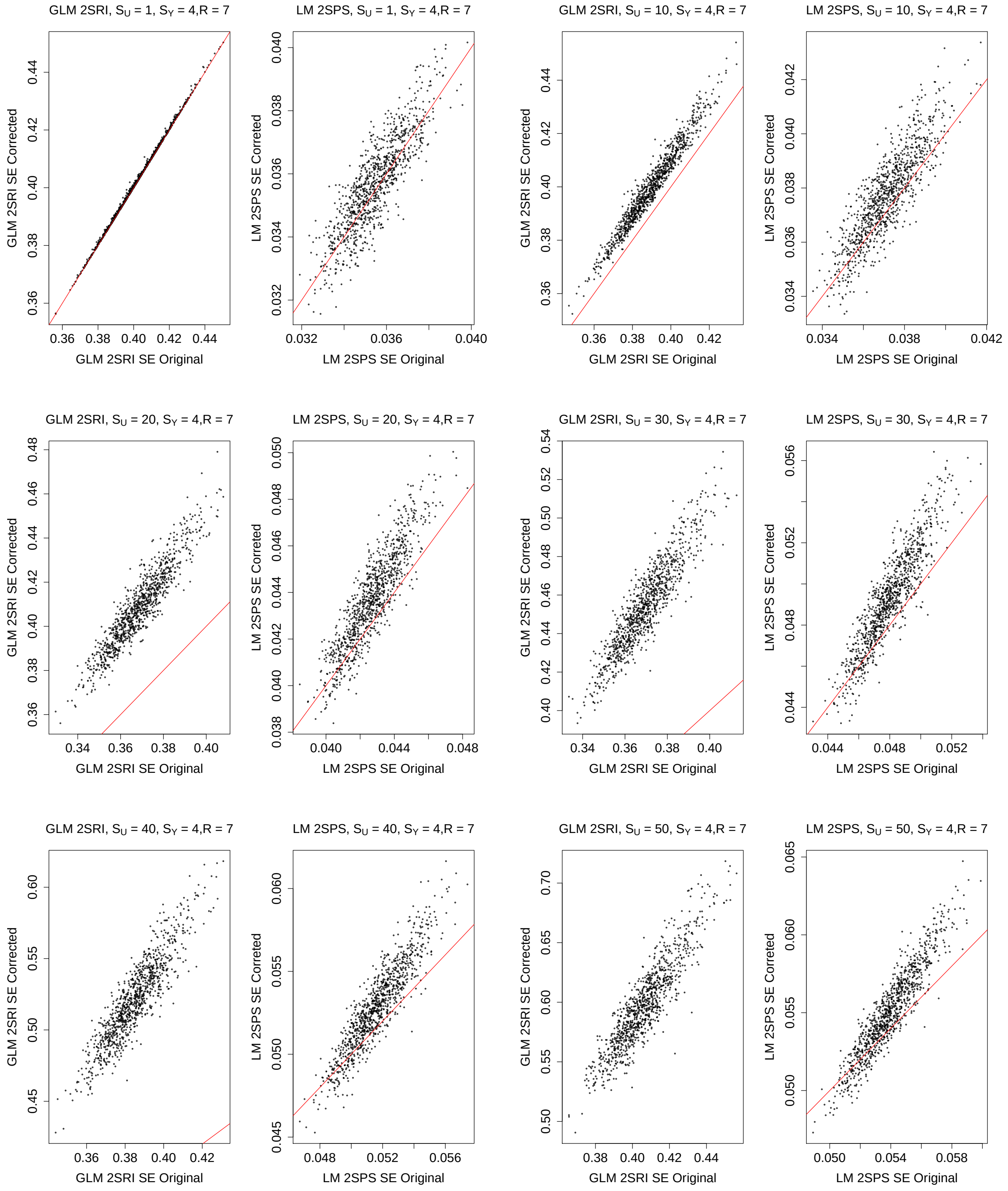

Figure S80: Simulations with the one-sample approaches: comparison of the original and corrected standard error estimates, $S_Y = 5, R = 7$.

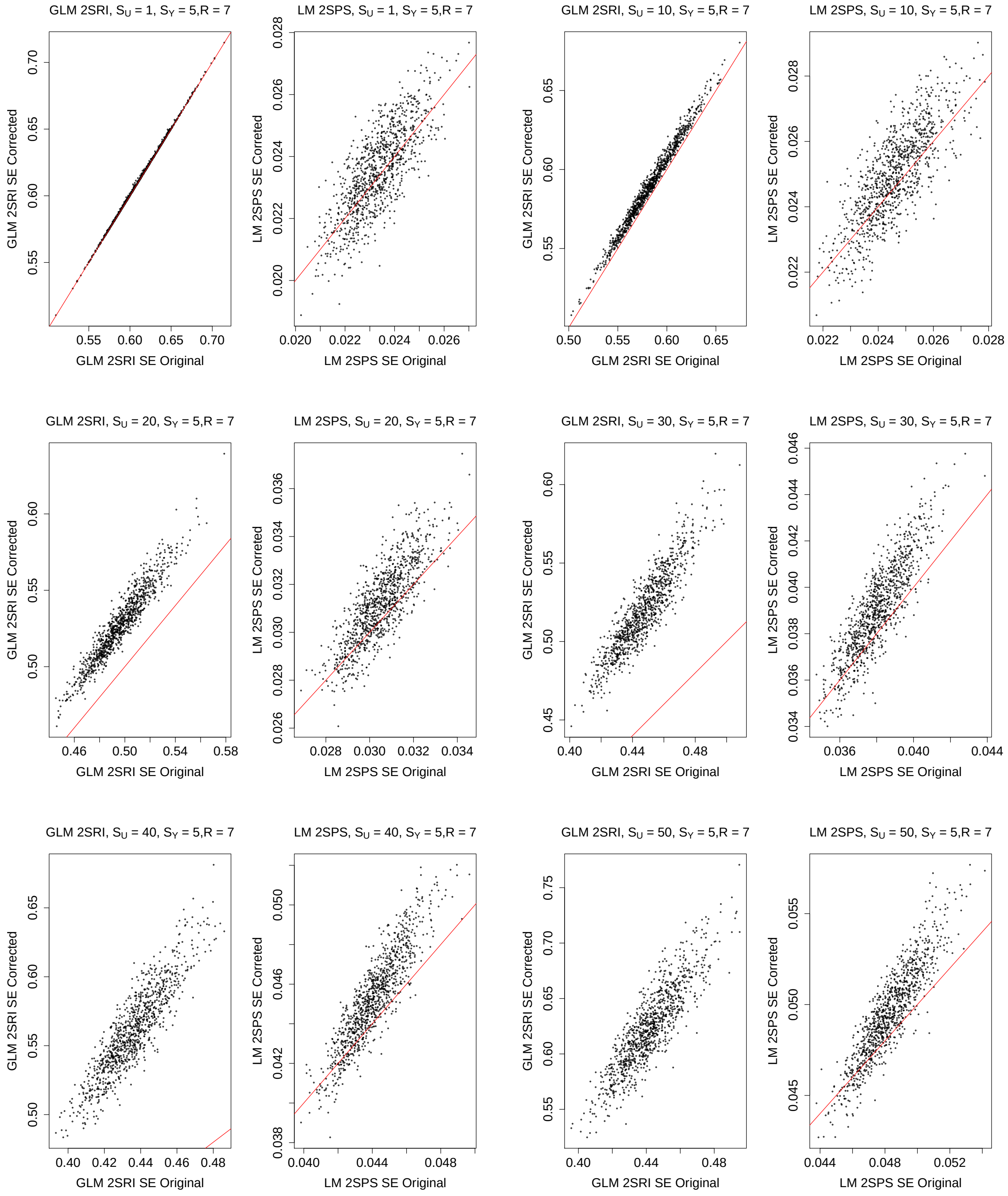

Figure S81: Simulations with the one-sample approaches: comparison of the original and corrected standard error estimates, $S_Y = 0, R = 9$.

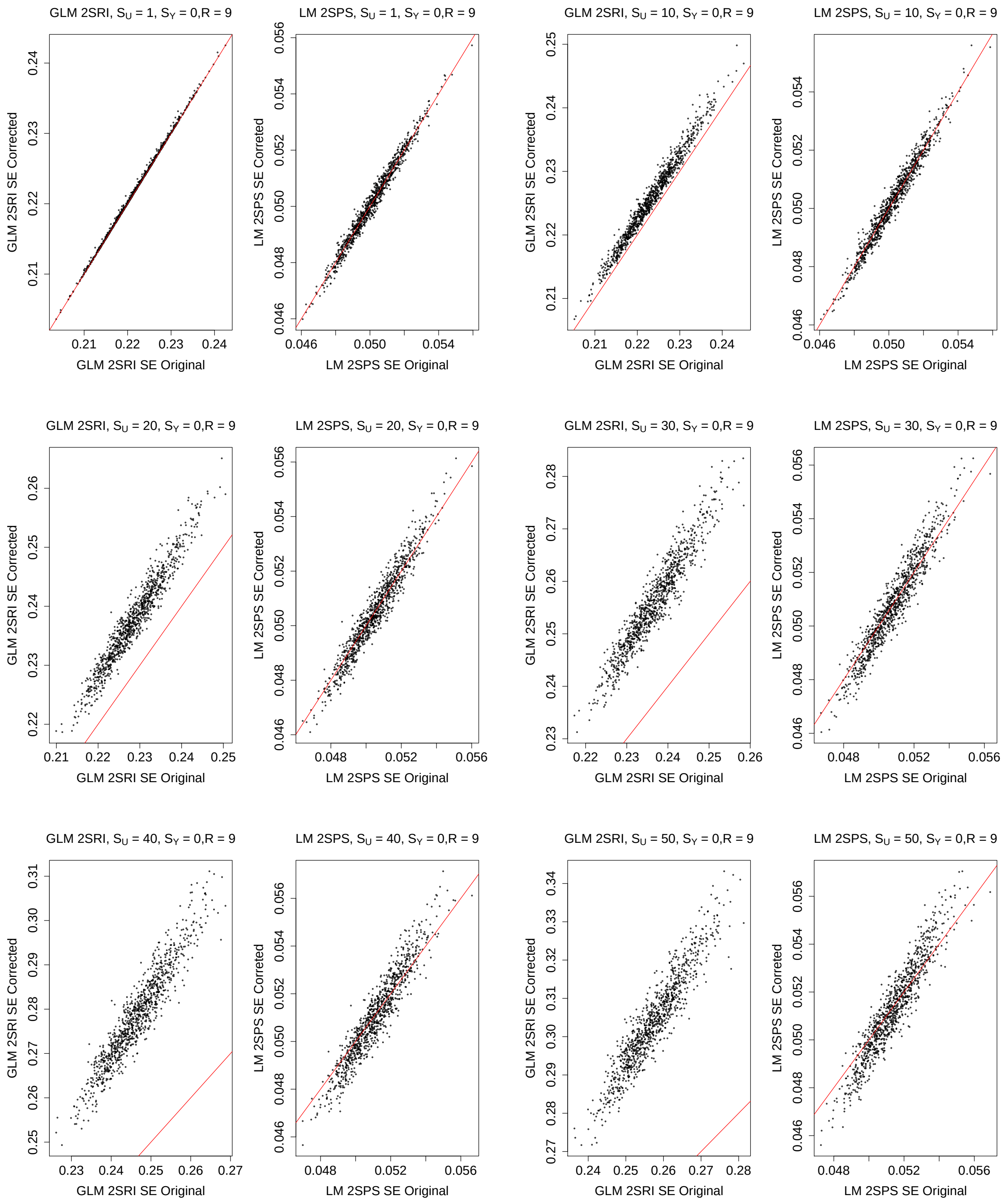

Figure S82: Simulations with the one-sample approaches: comparison of the original and corrected standard error estimates, $S_Y = 1, R = 9$.

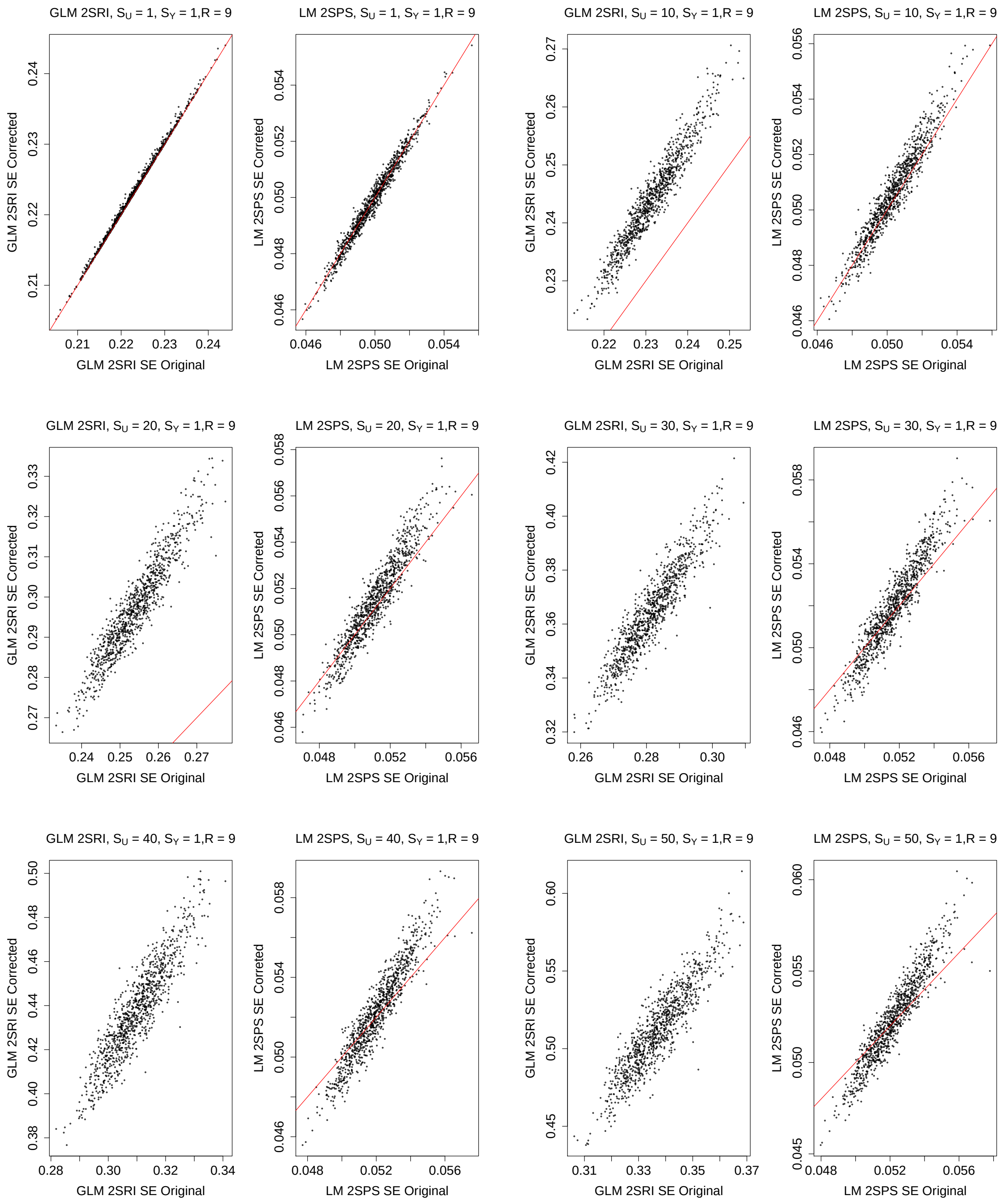

Figure S83: Simulations with the one-sample approaches: comparison of the original and corrected standard error estimates, $S_Y = 2, R = 9$.

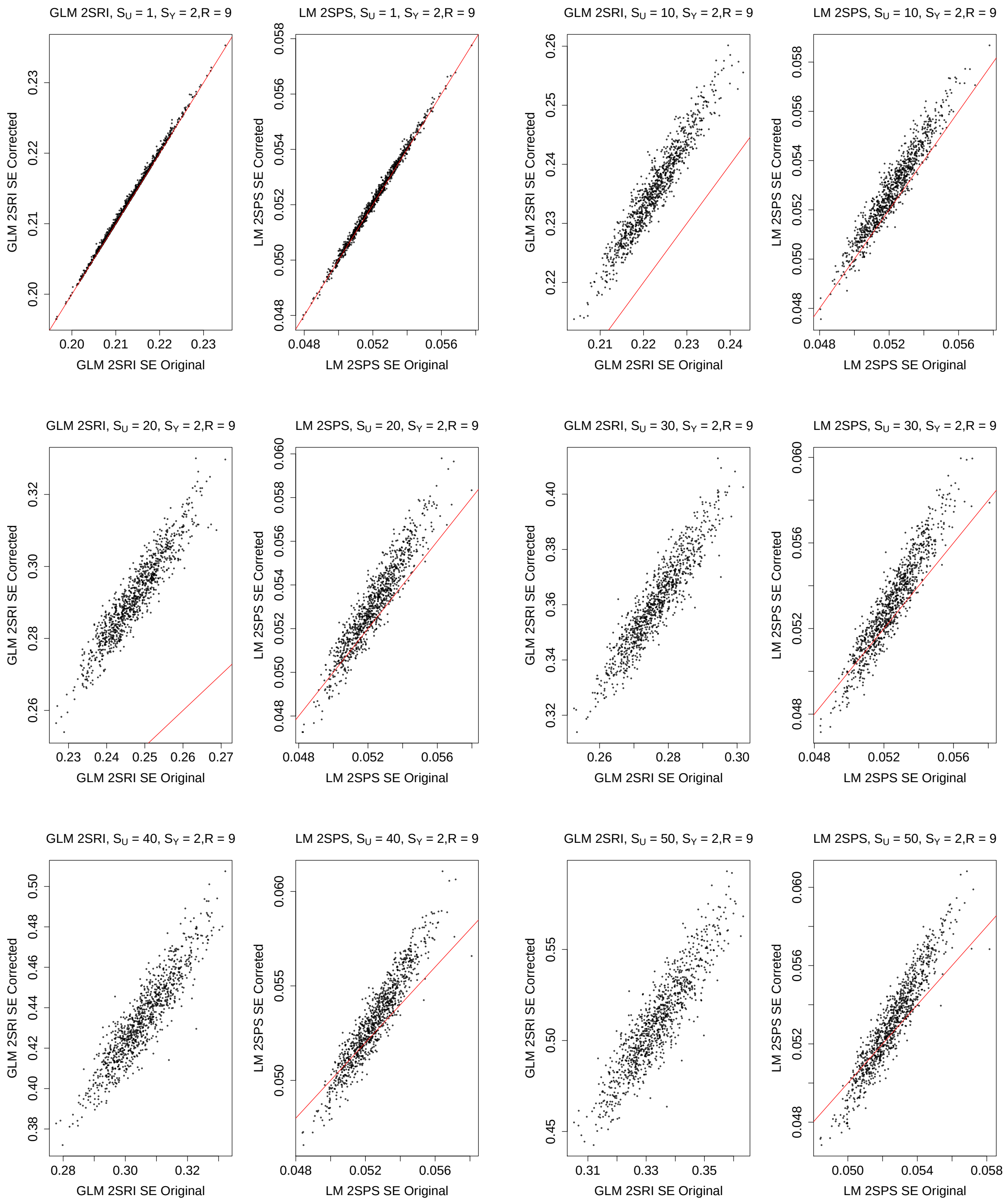

Figure S84: Simulations with the one-sample approaches: comparison of the original and corrected standard error estimates, $S_Y = 3, R = 9$.

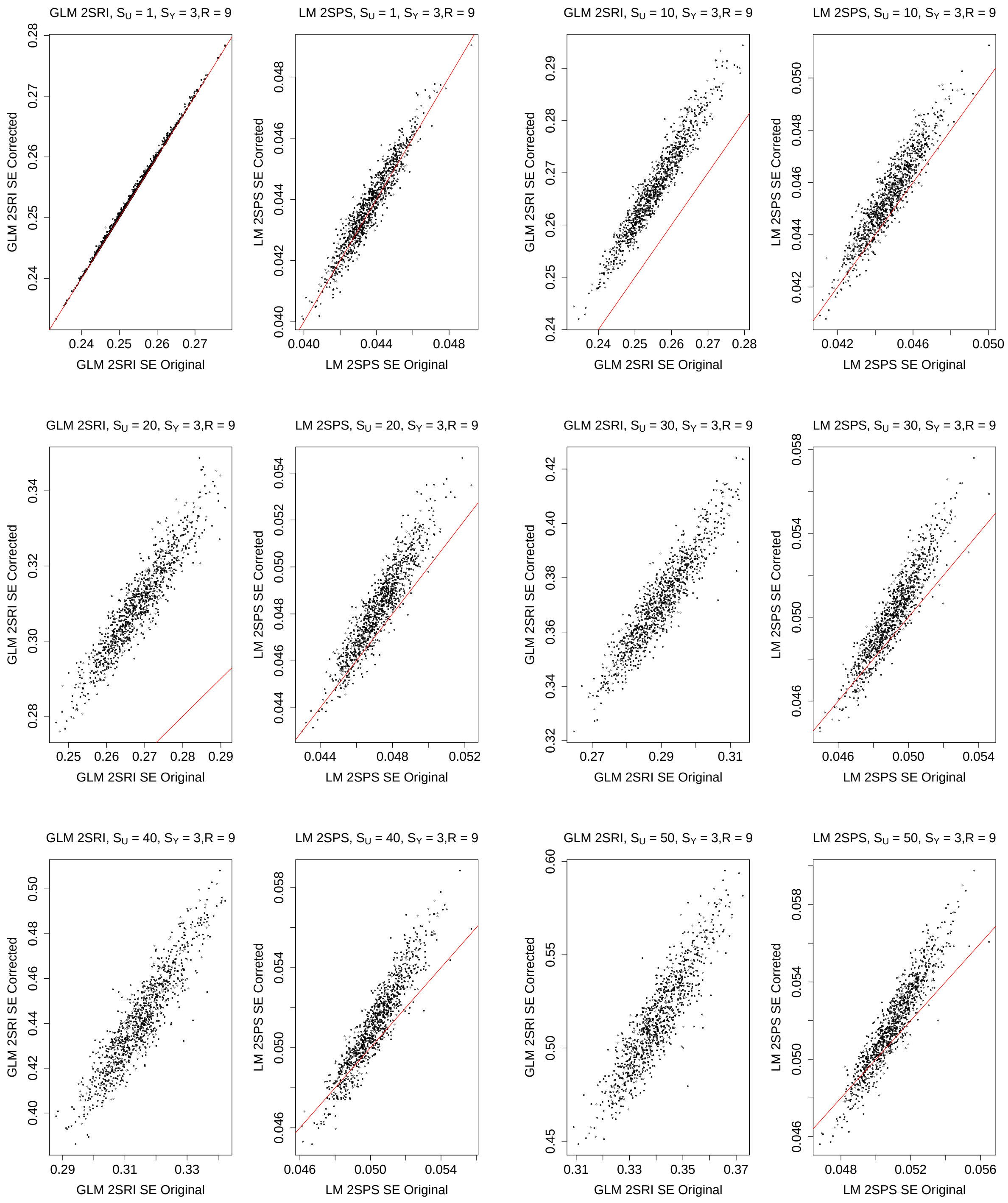

Figure S85: Simulations with the one-sample approaches: comparison of the original and corrected standard error estimates, $S_Y = 4, R = 9$.

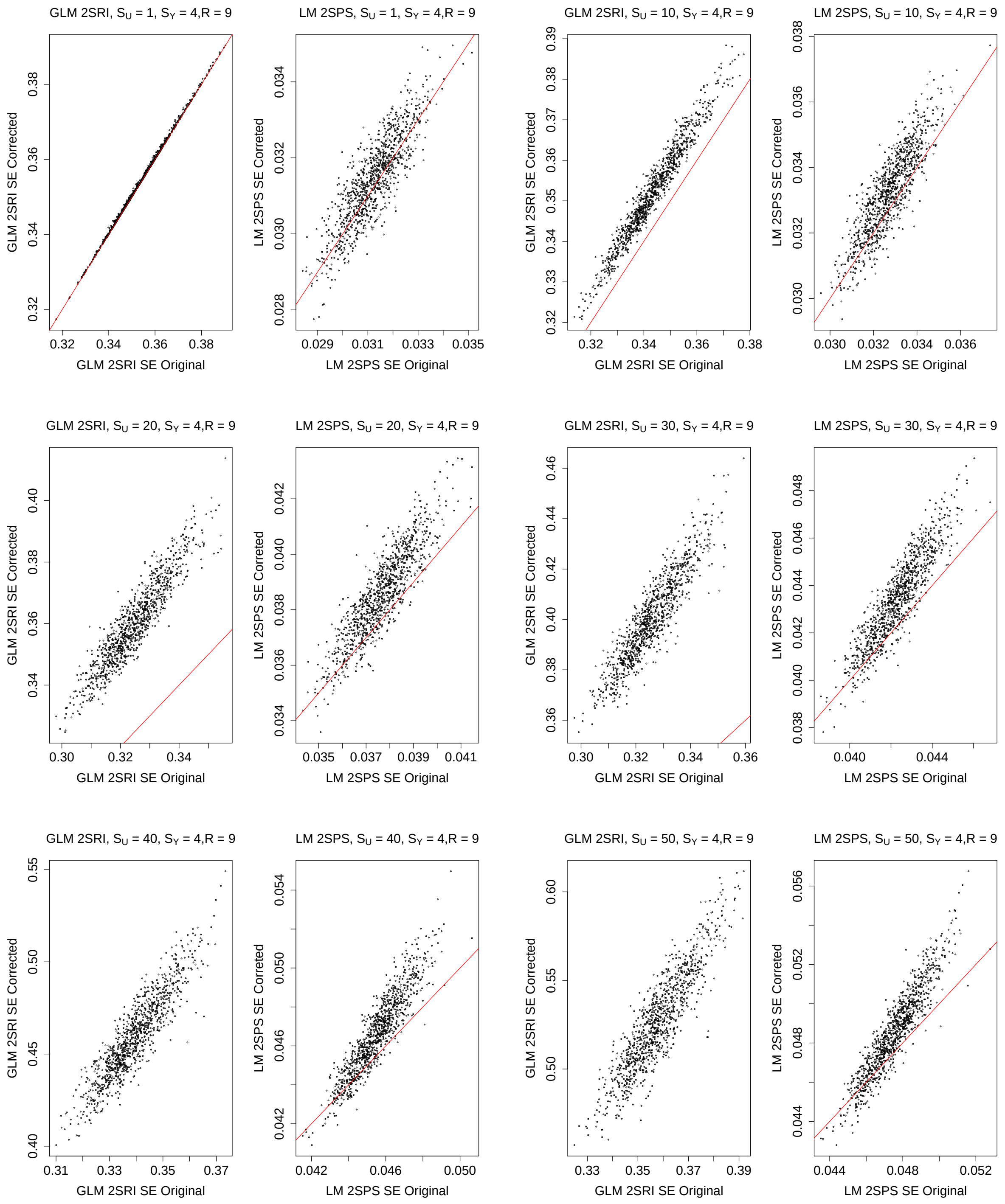

Figure S86: Simulations with the one-sample approaches: comparison of the original and corrected standard error estimates, $S_Y = 5, R = 9$.

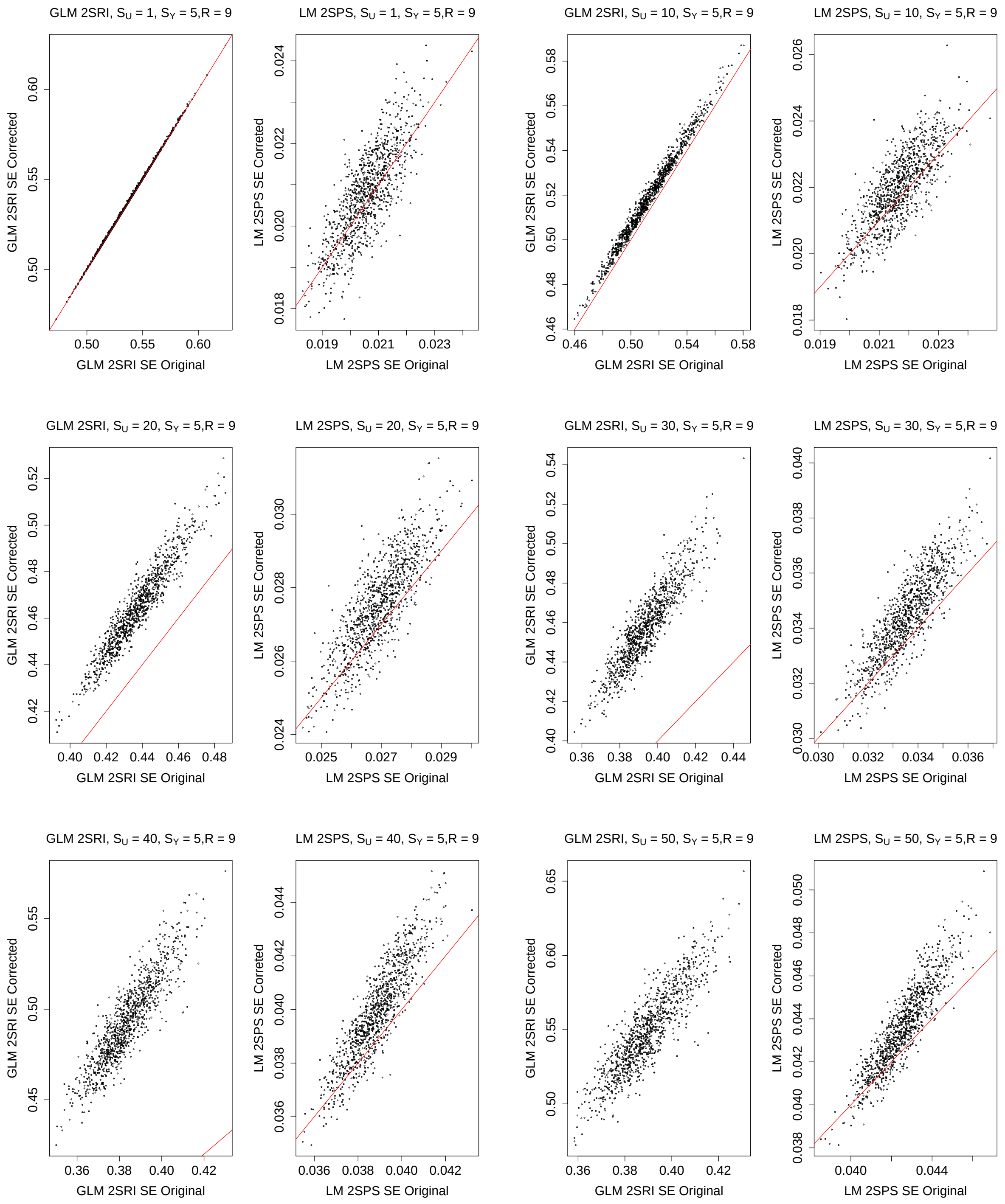
